# Cancer Hallmarks Define a Continuum of Plastic Cell States between Small Cell Lung Cancer Archetypes

**DOI:** 10.1101/2021.01.22.427865

**Authors:** Sarah Maddox Groves, Abbie Ireland, Qi Liu, Alan J. Simmons, Ken Lau, Wade T. Iams, Darren Tyson, Christine M. Lovly, Trudy G. Oliver, Vito Quaranta

**Affiliations:** Department of Biochemistry, Vanderbilt University, Nashville, Tennessee, USA; Department of Oncological Sciences, Huntsman Cancer Institute, University of Utah, Salt Lake City, UT 84112, USA; Department of Biostatistics and Center for Quantitative Sciences, Vanderbilt University Medical Center, Nashville, Tennessee, USA; Epithelial Biology Center and Department of Cell and Developmental Biology, Vanderbilt University School of Medicine, Nashville, Tennessee, USA; Division of Hematology-Oncology, Department of Medicine, Vanderbilt University Medical Center, Nashville, TN, USA; Vanderbilt-Ingram Cancer Center, Vanderbilt University Medical Center, Nashville, TN, USA

## Abstract

Small Cell Lung Cancer (SCLC) tumors are heterogeneous mixtures of transcriptional subtypes. Understanding subtype dynamics could be key to explaining the aggressive properties that make SCLC a recalcitrant cancer. Applying archetype analysis and evolutionary theory to bulk and single-cell transcriptomics, we show that SCLC cells reside within a cell-state continuum rather than in discrete subtype clusters. Gene expression signatures and ontologies indicate each vertex of the continuum corresponds to a functional phenotype optimized for a cancer hallmark task: three neuroendocrine archetypes specialize in proliferation/survival, inflammation and immune evasion, and two non-neuroendocrine archetypes in angiogenesis and metabolic dysregulation. Single cells can trade-off between these defined tasks to increase fitness and survival. SCLC cells can easily transition from specialists that optimize a single task to generalists that fall within the continuum, suggesting that phenotypic plasticity may be a mechanism by which SCLC cells become recalcitrant to treatment and adaptable to diverse microenvironments. We show that plasticity is uncoupled from the phenotype of single cells using a novel RNA-velocity-based metric, suggesting both specialist and generalist cells have the capability of becoming destabilized and transitioning to other phenotypes. We use network simulations to identify transcription factors such as MYC that promote plasticity and resistance to treatment. Our analysis pipeline is suitable to elucidate the role of phenotypic plasticity in any cancer type, and positions SCLC as a prime candidate for treatments that target plasticity.

## Introduction

Small cell lung cancer (SCLC) is a lethal malignancy of the airway epithelium, originating from pulmonary neuroendocrine cells (PNECs), and possibly other related cell types.^1–5^ SCLC tumors have long been considered homogeneous due to histological appearance as a carpet of uniform “small blue round cells,”^6, 7^ and to the virtually ubiquitous biallelic inactivation of tumor suppressors Rb and TP53.^8^ However, in recent years, accumulating molecular and functional evidence has led to the identification of distinct intratumoral SCLC transcriptional subtypes across several model systems, including cell lines, human tumors, and genetically engineered mouse models (GEMMs).^5, 9–13^ Phenotypic heterogeneity, both genetic and non-genetic, is intensively studied across cancer types because of its perceived impact on progression, acquired resistance, and relapse.^14–25^ Studies of intratumoral heterogeneity are especially relevant for SCLC, since cooperativity and transitions among SCLC subtypes have been reported but remain poorly understood.^11, 12, 26^

The phenotypic heterogeneity of SCLC cells has been provisionally classified into four consensus subtypes defined by enrichment for the transcription factors (TFs) ASCL1 (A), NEUROD1 (N), YAP1 (Y), or POU2F3 (P).^5, 9–13^ We also identified a fifth subtype, A2, which is driven by ASCL1 but is clearly distinct from the SCLC-A neuroendocrine subtype.^27, 28^ However, stark delineation of bulk cell lines or tumors into single subtypes has proven difficult and may inadequately describe SCLC intratumoral heterogeneity, since it is often the case that either multiple or none of the eponymous TFs are expressed in a population of SCLC cells. For instance, our work using CIBERSORT^29^ decomposition showed all tested SCLC tumors are composed of multiple subtypes, and several studies have reported changes of subtype prevalence during tumor progression or in response to treatment.^11, 27, 28, 30^ Bulk RNA-seq analyses confirm some samples are positive for more than one TF, such as human cell lines and tumors that are positive for both ASCL1 and NEUROD1.^31^ In bulk data, it is unclear if this is due to a mix of discrete subpopulations of cells, or if SCLC cell states are better represented as a continuum of gene expression between subtypes. These layers of heterogeneity suggest that single cell data may be necessary to fully parse subtype prevalence in SCLC cell lines and tumors. While approaches using various methods such as bulk sequencing,^9, 11, 28, 32^ DNA methylation profiling,^33–36^ or CRISPR^5^ experiments have been used to define subtypes of SCLC, the prevalence of subtypes at the single cell level has not been well-studied, and comprehensive gene signatures of each subtype for single cells currently do not exist.

Clustering methods, which identify “prototypical” gene expression profiles of cluster centers, have often been used in other systems to characterize subtypes. These clusters are easily interpretable but are often too rigidly defined in the case of mixed or intermediate samples. One method that is more flexible than clustering and yet remains easily interpretable is Archetypal Analysis (AA)^37, 38^ **(Figure 1A)**. Archetypal analysis has been used on systems ranging from cancer to development in order to explore heterogeneous gene expression by finding archetypes, or “pure subtypes,” in gene expression space that best explain the heterogeneity seen in the samples.^37–46^ Using AA on SCLC cells from cell lines and tumors gives us the flexibility to understand how plastic cells may pass through intermediate states,^26, 47^ which cannot be described in a discrete-clustering framework, to transition between subtypes.

**Figure 1:**
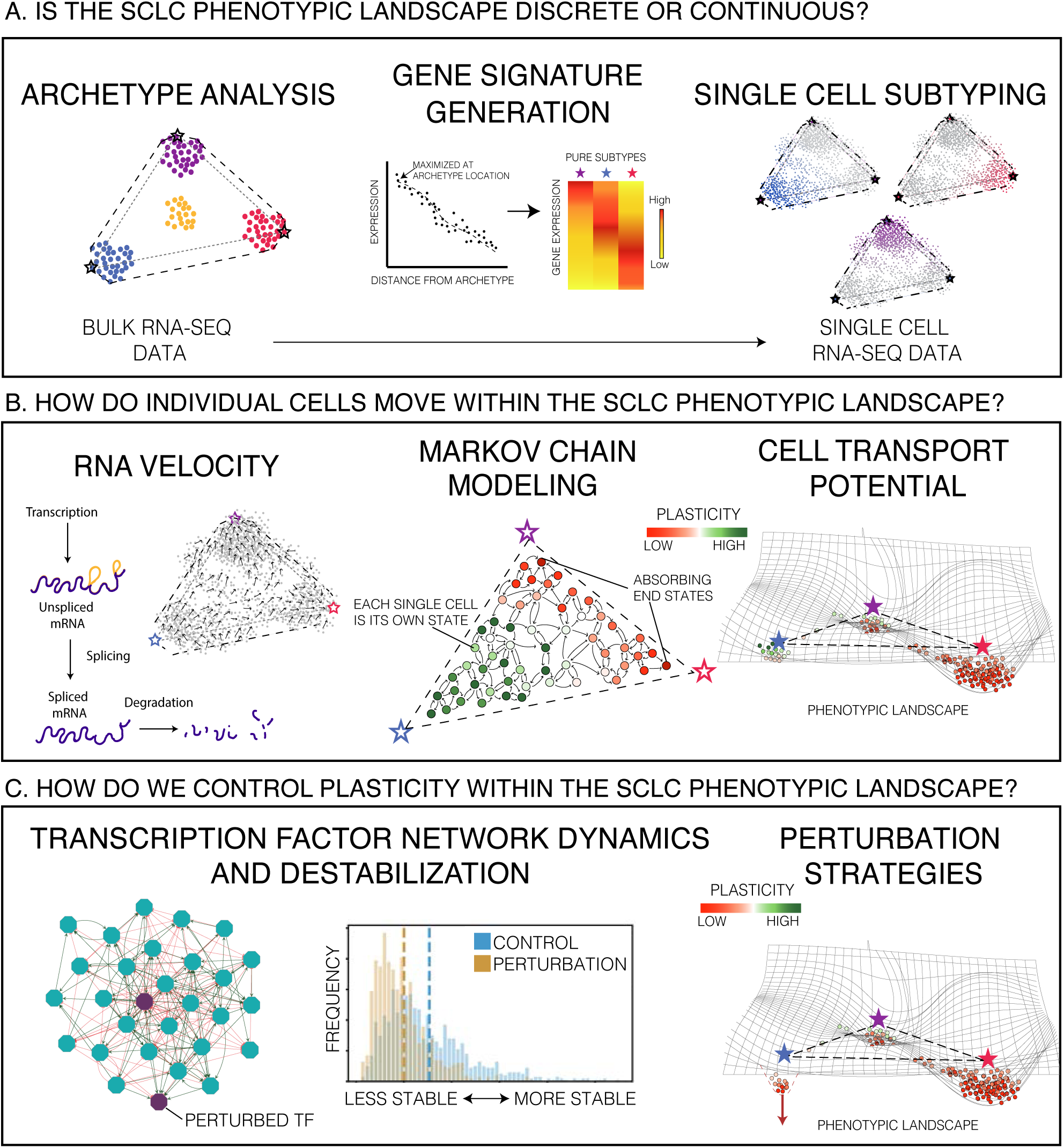
Analysis pipeline. A. *Is the SCLC phenotypic landscape discrete or continuous?* Archetype analysis (AA) and single cell subtyping are used to determine the geometry of the gene expression space for SCLC phenotypes. AA defines tasks optimized by each archetype that cells must tradeoff between. Single cell RNA sequencing data is then placed within this framework using a novel subtyping method. B. *How do individual cells move within the SCLC phenotypic landscape?* Using RNA velocity, a Markov Chain model defines likelihood of transitions between cell states in the landscape. Cell transport potential (CTrP) is calculated as the expected distance of travel for each cell in gene expression space. C. *How do we control plasticity within the SCLC phenotypic landscape?* Using TF network analysis with BooleaBayes, we investigate the structure of the network and inferred rules of interaction between nodes. Network simulations show likely paths of transition under normal dynamics of the system, which can be used to efficiently trade-off between archetype tasks.

Furthermore, in several other systems low-dimensional geometry of data has recently been attributed to evolutionary tradeoffs between multiple functional tasks.^38, 45^ When cancer cells with limited resources (e.g. metabolic constraints) must optimize fitness in the face of multiple competing tasks, such as proliferation and migration,^48, 49^ they fill a polygonal shape between archetypes in gene expression space. Each archetype optimizes a single task, and cells between archetypes fall somewhere within the Pareto front, which is the set of gene expression profiles that cannot optimize all tasks at once.^38^ We analyzed the low-dimensional polytope of single cell data from SCLC cell lines within this context and found the main trade-offs SCLC cells include proliferation, invasion, and immune regulation, which mirror the cancer hallmarks proposed by Hanahan et al.^50, 51^ and the corresponding universal cancer tasks proposed by Alon and coworkers.^45^ Where each cell falls with respect to the archetypes determines how specifically it optimizes a single task (specialists near an archetype), or how it has generalized in order to complete several tasks (near the center of the polytope or along an edge/face between two/three tasks) (**Figure 1A**). If the proportions of tasks needed changes rapidly, such as during tumor evolution and metastasis, a population of generalists that exists along a continuum of states may have an advantage. We show here that SCLC cell lines and tumors comprise continuums with both specialists and generalists.

SCLC has been shown to contain low amounts of non-tumor cells relative to other tumor types,^8^ raising the question of how SCLC compensates for the lack of trophic support usually provided by other non-cancer cell types in the tumor microenvironment. One hypothesis is that SCLC cells can easily diversify and shift between phenotypes to fulfill these different tasks.^52^ Using a novel metric to quantify plasticity, Cell Transport Potential, we found that SCLC cells from human cell lines diversify across the archetype space; in the cell lines studied, we found subpopulations of high and low plasticity within each sample. Similarly, SCLC tumors tended to have high-plasticity non-neuroendocrine cells that could shift phenotypes to generalists and other cell types to optimize specific tasks. We visualized plasticity within archetype space to determine how SCLC cells can traverse the landscape (**Figure 1B**). Importantly, our continuous subtyping and plasticity framework allows for characterization of these phenotypic transition paths, including characterization of TF network dynamics over transition paths, which cannot be adequately captured by a 4-TF framework (**Figure 1C**).

Overall, our work advances the field’s understanding of SCLC heterogeneity and plasticity by unveiling the prevalence of cells that fall in between extreme subtypes, thus demonstrating a need for a more flexible method of phenotype characterization. We provide a theoretical basis for the existence of intermediate “generalist” cells, which have not been captured in previous subtyping efforts using single TFs, by enumerating tasks that SCLC tumors must trade-off between to thrive. We propose a new quantitative definition of cell plasticity, Cell Transport Potential, and use this method to determine that SCLC subtypes are capable of manipulating their plasticity according to the tasks required of them and the context in which they grow. Lastly, we show that TF network modification and dynamics may be key in understanding ubiquitous SCLC tumor relapse and acquired resistance.

## Results

### Archetype analysis defines a phenotypic continuum for SCLC subtypes

Phenotypic heterogeneity of Small Cell Lung Cancer (SCLC) is reflected in the large number of established SCLC cultured cell lines that span morphologies (classic vs. variant), gene expression profiles, and treatment response.^10, 53, 54^ Therefore, to characterize SCLC transcriptional heterogeneity we analyzed the gene expression variance of a bulk RNA-seq dataset comprising 120 SCLC cell lines (**Supplemental Figure 1**). Resulting SCLC clusters were not compact and well-separated, as determined by the Dunn Index (**Figure 2A**, **Supplemental Figure 2A and B**). In addition, several cell lines fell in-between clusters, possibly because they comprise two or more canonical subtypes.^31^ Therefore, we analyzed the bulk RNA-seq cell line dataset with Archetype Analysis (AA), a method that allows for a more flexible characterization of SCLC gene expression space.^55^

**Figure 2:**
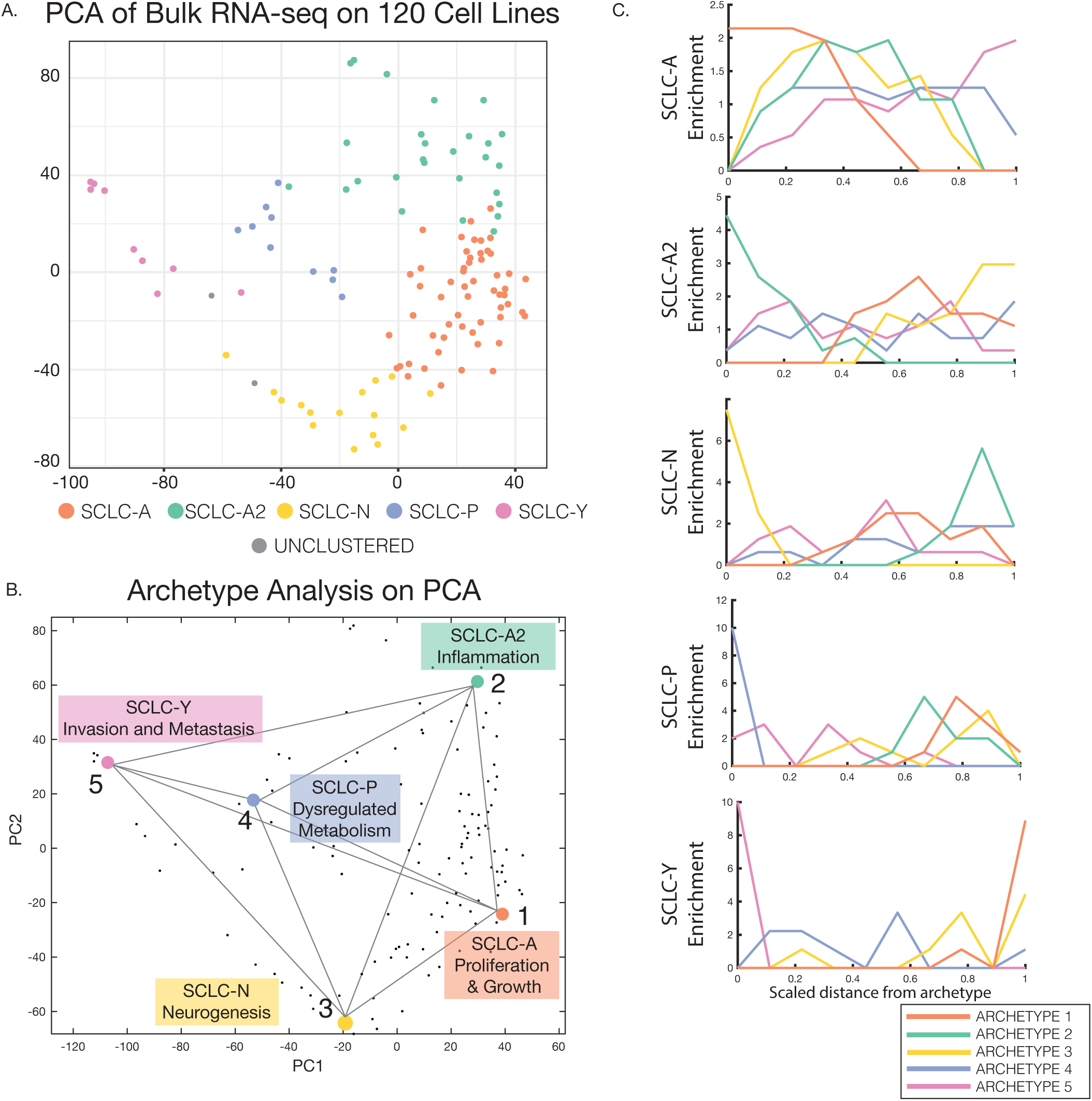
PCA and Archetype Analysis on SCLC Cell Line Bulk RNA-Seq Data. A. PCA of bulk RNA-seq on 120 cell lines showing the first two components. Cells are colored according to subtype clustering from **Supplemental Figure 2A**. B. PCHA shows 5 archetypes fit the cell line data well (p = 0.034). Each archetype location is enriched in one or more cancer-related tasks and corresponds to a previously defined subtype (A = orange, A2 = teal, N = yellow, P = blue, or Y = pink). C. Subtype label enrichment by distance from each archetype. Data was binned according to distance from each archetype (x-axis), and enrichment of each subtype label (y-axis) was computed using a hypergeometric test. Enriched subtypes are highest at X=0, in the bin closest to one of the archetypes, and lowest near all other archetypes. Each archetype shows enrichment in one of the five SCLC subtypes from literature.

Briefly, AA approximates the cell-phenotype space as a low dimensional shape called a convex polytope. The vertices of this multi-dimensional shape represent archetypes (**Figure 1A**), constrained to be linear mixtures of some set of data points.^38^ To determine the optimal number and location of the archetype vertices in gene-expression space, we applied the Matlab package *ParTI*.^55^ Considering the variance each archetype was able to explain, we found that 5 archetypes best fit the data (**Figure 2B**, p-val = 0.034, t-ratio test, see Methods and **Supplemental Figure 2C**). By bootstrapping the location of the vertices, the archetypes in gene expression space were robust and not dependent on any extreme points in the dataset (**Supplemental Figure 2D**). Within this 5-dimensional polytope, data points corresponding to bulk RNA-seq from each of the 120 cell lines were then represented as linear combinations of the archetypes (**Figure 2B**). In summary, AA explained SCLC cell line heterogeneity in bulk transcriptomics data as a low-dimensional phenotypic continuum between 5 archetype vertices (**Figure 2B**).

To reconcile these archetypes with the canonical SCLC consensus subtypes,^11, 28^ we computed the enrichment of SCLC subtype labels **(Supplementary Figure 2A)** in cell lines closest to each archetype versus the remaining cell lines (hypergeometric test, q < 0.1, see Methods). After binning the data into ten bins by distance to archetype, we found that each archetype was enriched in cell lines from one of the five SCLC subtypes (**Figure 2C, Supplemental Table 1**). For example, canonical SCLC-A-labeled cell lines were enriched in the bin closest to archetype 1 (orange line), but no SCLC-A cell lines were found in bins closest to the other four archetypes (**Figure 2C**). Likewise, bins closest to the remaining 4 archetypes each contained cell lines from the remaining 4 canonical subtypes (**Figure 2C**). Thus, the AA-derived 5-dimensional SCLC phenotypic continuum integrated well with the canonical SCLC subtypes, with the distinct advantage that cell lines with a bulk transcriptome that does not adhere to any of the canonical subtypes can now be understood as intermediate between archetypes, rather than being forced into discrete clusters unrepresentative of their phenotype. Furthermore, cell lines previously ill-defined due to lack of expression of one of the eponymous TFs can be classified in the phenotypic continuum based on distance from archetype.

### Trade-offs between cancer hallmark tasks define the continuous phenotypic space of SCLC

In AA, a phenotypic continuum can arise when each archetype corresponds to a functional task. Multi-objective evolutionary theory suggests that in tissues where multiple tasks are required, such as the intestinal crypt and liver,^38, 40, 44, 55^ cell trade-offs between these tasks form the low-dimensional polytope shape of the data. In the case of cancer cells in a tumor, such trade-offs presumably occur among tasks that promote tumor growth and survival, such as cancer hallmarks.^45, 50, 51^ To assign possible cancer tasks, we evaluated enrichment of gene ontologies at each SCLC archetype location (**Figure 2B, Supplemental Tables 2-4**).^56–58^ We used the molecular signatures database (MSigDB) to evaluate hallmark gene sets^57^ and ontology biological processes, and the Cancer Hallmark Genes database (CHG^59^) to evaluate ten cancer hallmarks.^50, 51^ For each archetype, we then considered as significant the gene sets that were maximal in the ten percent of cell lines closest to that archetype compared to the remaining cell lines (Bonferroni-Hochberg-corrected q < 0.1). We used a leave-one-out procedure for each gene in the gene sets to avoid circularity for when a gene is also used to define the archetype location.^55^

Archetype 1 corresponded to the SCLC-A Subtype (**Figure 2B and C**) and was enriched in neuronal differentiation genes (**Table 1**). While no hallmark gene sets from CHG were significantly upregulated in this archetype, we found evidence of lung cell-specific and neuronal type-specific proliferation and growth in enriched GOs. The highly proliferative nature of these cells was consistent with the large proportion of SCLC-A cells in primary SCLC tumors and high sensitivity to DNA damaging agents, and the neuronal-specific GO enrichment reflected their neuroendocrine nature.^53, 60, 61^ Archetype 2 (SCLC-A2) was also driven by neuronal programs but was enriched for the cancer hallmarks *evading immune destruction* and *tumor-promoting inflammation*. As shown in **Table 1**, the SCLC-A2 archetype is also enriched in response to the environment and chemical homeostasis functions. Together, optimization of these tasks may allow SCLC-A2 cells to quickly and effectively respond to and interact with the tumor microenvironment by sensing and responding to immune infiltration and signals from other stromal cells.

**Table 1:**
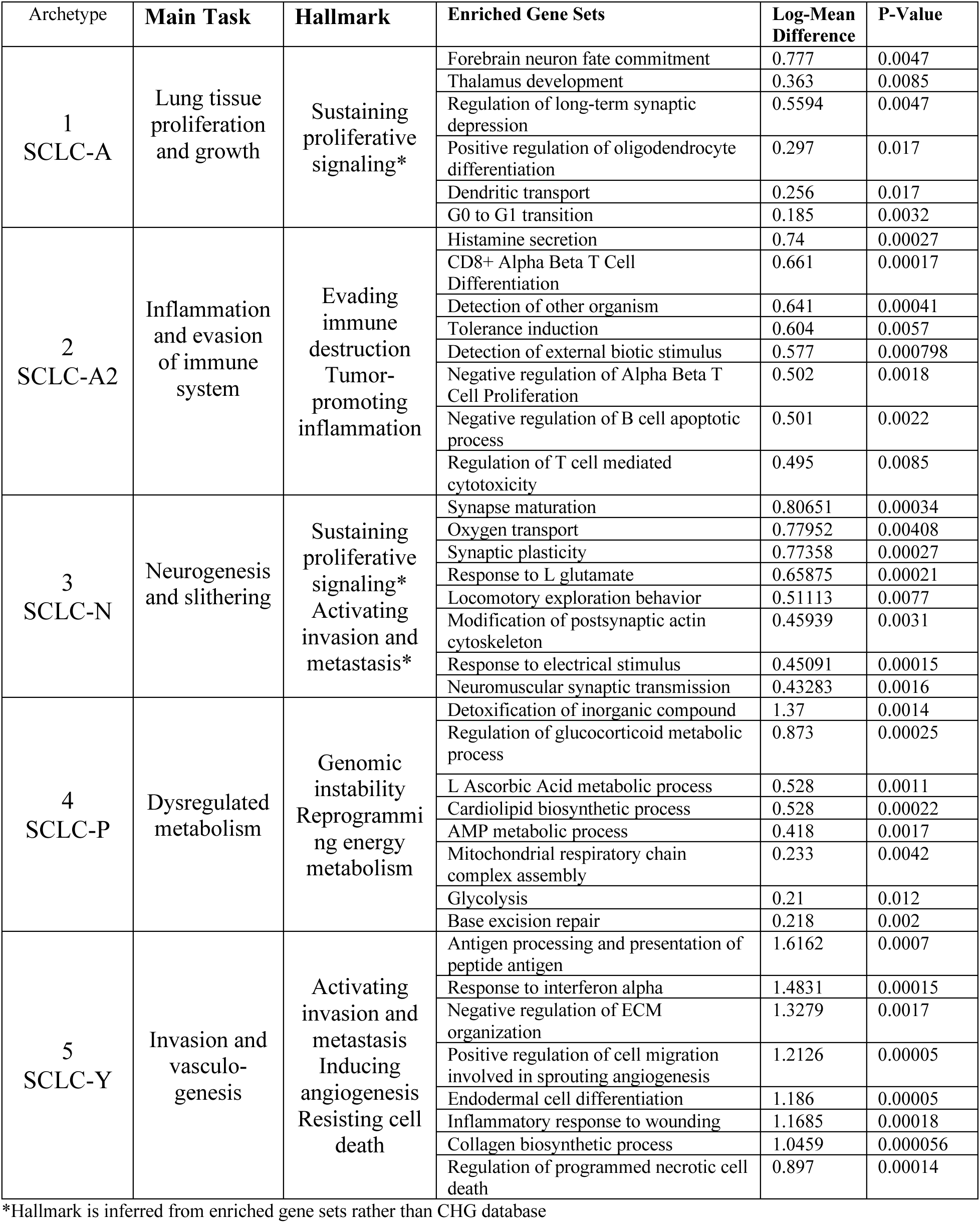
SCLC Archetype Gene, Gene Set, and Task Enrichment. This table details gene sets from MSigDB that are enriched in cell lines closest to each archetype by Mann-Whitney test on binned data (FDR q <0.1). Only features that are highest in the bin closest to archetype are considered.

Archetype 3 (SCLC-N) was enriched in neurogenesis terms, including synapse and actin cytoskeleton terms. Specific to neuronal cell types, these may enhance tumor survival by specifying a protruding, axon-like morphology. Such a morphology is related to the slithering observed in PNECs^62^, whereby cells transiently downregulate adhesion genes and use axon-like protrusions to migrate. Thus, Archetype 3 may be relevant to invasion and metastasis, yet another cancer hallmark. Archetype 4 (SCLC-P) was enriched in the cancer hallmarks of *genomic instability* and *dysregulated energy metabolism*. Enrichment in glycolysis, mitochondrial respiratory chain, and other metabolic terms corroborated this task. Archetype 5 (SCLC-Y) was the only archetype enriched in all ten cancer hallmarks, suggesting it may be the key to understanding the aggressiveness of SCLC. Specific to this archetype, we found enrichment in inducing *angiogenesis* and *invasion* hallmarks. Cells closest to this archetype vertex were enriched in tissue remodeling functions such as regulation of extracellular matrix (ECM), endodermal differentiation, cell migration, and inflammation, further confirming the cancer hallmark tasks we ascribe to this archetype. Furthermore, this archetype was optimized for *resisting cell death*, as shown by enrichment in anti-apoptotic genes (**Table 1**), consistent with the role of SCLC-Y cells in tumor relapse.^11, 32^

These findings indicated that the vertices of the SCLC phenotypic continuum derived from AA were well aligned with the known biological properties of SCLC tumors and subtypes, supporting the validity of this approach. Thus, it is expected that cell lines closer to the archetypes would behave as “specialist” populations that specialize in the corresponding task, while cell lines in between archetypes may comprise multiple specialist subpopulations or contain intermediate “generalist” cells. Distinguishing between these two possibilities cannot be done with bulk RNA-seq and requires projection of these bulk-derived archetypes onto single-cell data, as we did in the next section.

### Projection of SCLC archetype signatures onto single cells shows a continuum of phenotypic states in human cell lines

For cell lines falling in-between archetypes (**Figure 2A, B**), two possibilities could be envisioned: they may comprise multiple subpopulations each optimized for a different task, or they may have an intermediate, “generalist” phenotype with suboptimal task performance. To distinguish between these possibilities, we used the archetype locations to generate archetype gene expression signatures capable of identifying phenotypes at the single-cell level (**Supplemental Figure 3A**). We found genes that were enriched in cell lines closest to each archetype (Mann-Whitney Test, q < 0.1, **Supplemental Table 5**). We then selected genes with the largest enrichment in cell lines binned closest to each vertex versus the remaining cell lines, in order to ensure that these genes were maximized in the bin closest to the archetype (see Methods, **Supplemental Figure 3B, Supplemental Table 6**).

Several genes were able to distinguish between the SCLC-A and the SCLC-A2 archetypes, including ELF3, GRP, and ISL1, while others, such as ASCL1 and FLI1, were shared by both. This corroborated our finding^28^ of two distinct transcriptional programs. For example, in the SCLC-A archetype, but not A2, SOX1 may promote neuronal stemness since this TF acts as an oncogene in glioblastoma, suppresses cell migration in non-neuroendocrine lung cancers, and maintains an active neural progenitor state in normal development.^63–66^ The SCLC-N signature, which included NEUROD1 and other NEUROD family genes (NEUROD2 and NEUROD6), was enriched for the biological process “nervous system development” (GO:0007399, p = 0.0014), consistent with the neuroendocrine nature of this subtype.^12^ MYC absence in the N gene signature was notable, but most likely due to upregulation across multiple archetypes (**Supplemental Figure 3C**), consistent with recent literature suggesting MYC plays an important role in the transitions between subtypes N and Y, and possibly P. ^11, 12, 47, 67^

POU2F3 was significantly overexpressed in the tuft cell-like SCLC-P archetype signature, as expected. Several other genes previously associated with tuft cells and SCLC-P^5^—SOX9, GFI1B, and AVIL—were enriched in this archetype signature as well. Lastly, in the SCLC-Y subtype signature, YAP1 expression was enriched, as well as multiple genes regulated by the transcription factors RARG and SOX2 (adjusted p = 0.055 and 0.034, respectively), as determined using ENRICHR.^68, 69^ These two factors are overexpressed in lung squamous cell carcinoma,^70^ consistent with the non-NE nature of SCLC-Y. SOX2 has also been implicated in lineage plasticity^71^ and SCLC progression.^72, 73^ The top two genes enriched in the SCLC-Y archetype, LGALS1 and VIM, are associated with a mesenchymal phenotype and have previously been implicated with SCLC chemoresistance.^36, 74–77^

To examine enrichment of these signatures at the single-cell level, we sequenced a panel of 8 cell lines chosen to maximally span the phenotypic continuum (**Figure 3A**). We then projected the single cell expression data into “archetype space” by the signature matrix using least-squares approximation (see Methods, **Supplemental Figure 4**). We thus derived archetype scores for each single cell sampled and investigated the subtype composition of each cell line with respect to these scores. To visualize these five archetype scores for each cell in two-dimensional plots, we used a Locally Linear Embedding (LLE, **Figure 3B**). **Figure 3B** shows that each cell line occupied a distinct region in archetype space, as expected from the bulk transcriptomes shown in **Figure 3A**. Furthermore, for each of the cell lines, the single cells did not comprise several discrete specialist populations, but instead formed a continuum between archetype subsets.

**Figure 3:**
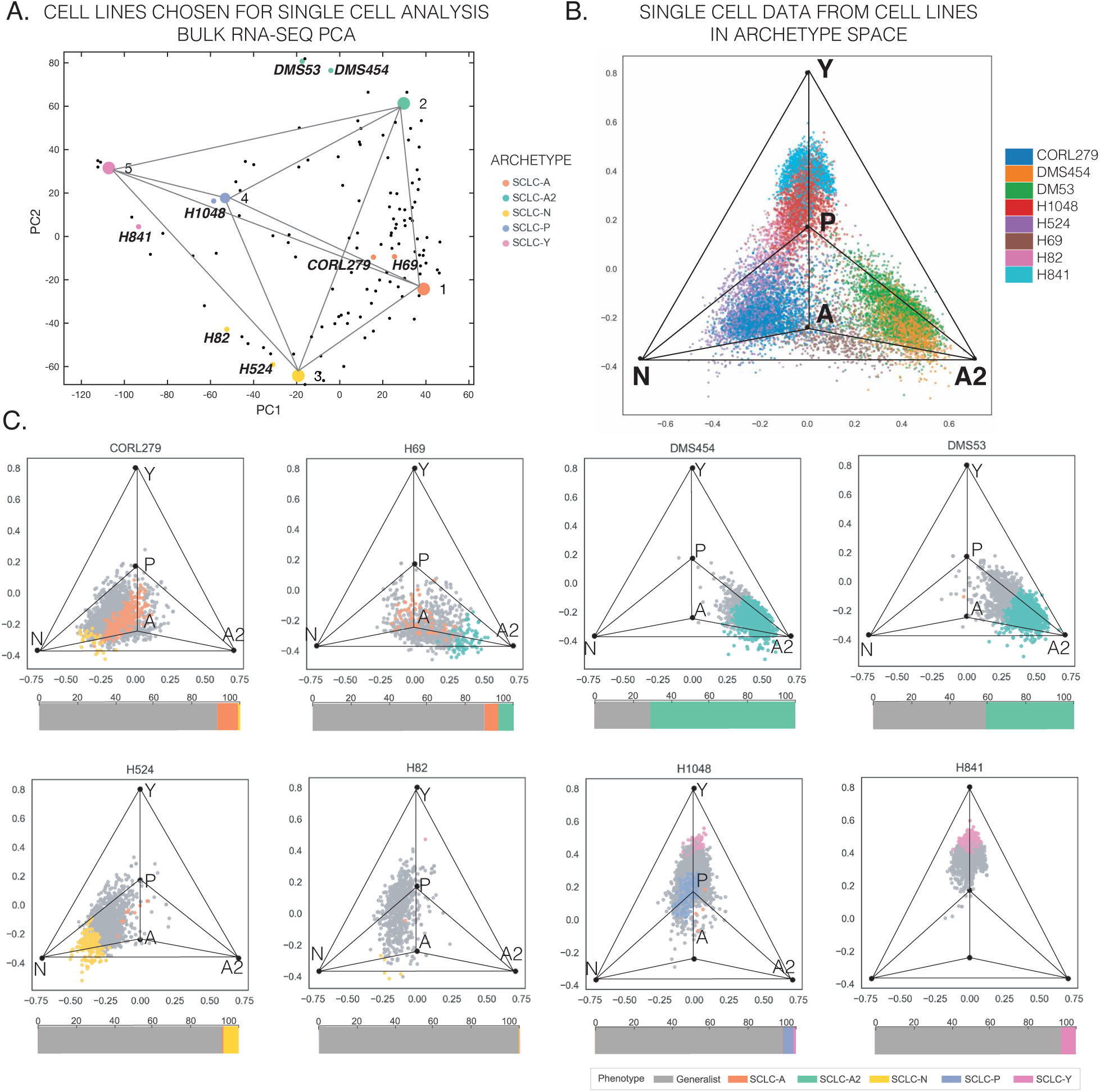
Single Cell Subtyping on 8 SCLC Cell Lines that Span the Phenotypic Space. A. Cell lines for scRNA-seq were chosen to span the phenotypic space of SCLC. Two cell lines from each neuroendocrine subtype (A, A2, and N) were chosen, and one from each non-neuroendocrine subtype (P and Y) was chosen. B. Locally linear embedding (LLE) of subtype composition for each cell line. As detailed in Methods, cells were scored using a least squares approximation method and embedded in two dimensions using LLE. Each cell line fills a different region of SCLC phenotypic space. C. LLE of subtype scores by cell line. Vertices in each figure represent one of the 5 archetypes. Cells with a maximum score greater than 0.5 were assigned to the corresponding subtype. Cells with a mix of positive scores between 0.1 and 0.5 were assigned a label of “generalist,” and cells with scores less than 0.1 for all archetypes were assigned a label of “None.” LLE allows for a reduction in dimensionality from five subtype scores to two dimensions and shows that each cell line spans a continuum of generalists between at least two subtypes.

We then used the archetype scores to evaluate the distance of each single-cell population (i.e., a cell line) to the archetypes to determine the dominant tasks being optimized. Cells closest to each archetype, defined by a score greater than 0.5 for a single archetype (see Methods), were considered specialists for that archetype task. Cells with an intermediate phenotype, where multiple archetype tasks were represented, were considered generalists, and cells without any high scores (>0.1) were considered unclassified (“None”). The ubiquity of single cells with a mixed phenotype in each cell line confirmed the need for a continuous framework (**Figure 3C**). Each of the cell lines formed a continuum between subsets of SCLC archetypes, with varying proportions of specialists and generalist cells (**Figure 3C**). For example, CORL279, which is positive for both ASCL1 and NEUROD1 at the bulk expression level, had a large proportion of generalists and formed a continuum between the A and N archetypes. The cell line H82, which falls between SCLC-N and SCLC-Y (**Figure 3A**), had the highest proportion of generalist cells. Virtually all cells could be assigned to specialists or generalists, with negligible numbers of cells left unclassified.

These results indicated that applying archetypal signature scores to SCLC cell lines, some of which fall in-between archetypes in bulk RNAseq transcriptomics (**Figure 3A**), reveals to what extent they are a composite of specialist (close to a single archetype) and generalist (mixed phenotype) single cells. While archetypal signatures are flexible enough to uncover such mixed cells, the framework does not ensure *a priori* that mixed generalist cells will exist. In other words, such a framework could also describe populations of entirely-specialist cells, suggesting the generalist cells we detected here are truly of an intermediate phenotype.

### A phenotypic continuum of specialist and generalist cells is detected in SCLC tumors

To determine whether a continuum of cell states exist in tumors as well, we applied our archetypal signature score to single-cell gene expression data from SCLC human tumors (**Supplemental Figure 5**) and GEMMs (**Supplemental Figure 6 and 7**).

In human tumors (**Figure 4A**), archetype signature scoring shows that Tumor 1, a bronchoscope biopsy from an SCLC relapse, was enriched in A and A2 cells. Instead, Tumor 2, a treated, limited-stage (1B) SCLC tumor with a large cell neuroendocrine carcinoma (LCNEC) component, was enriched for SCLC-Y. In both cases, a subpopulation of generalist cells spanned the A, A2, P, and Y archetypes to different degrees, again supporting the presence of a cell-state continuum. Furthermore, generalist cells with intermediate phenotypes were present in every sample tested.

**Figure 4:**
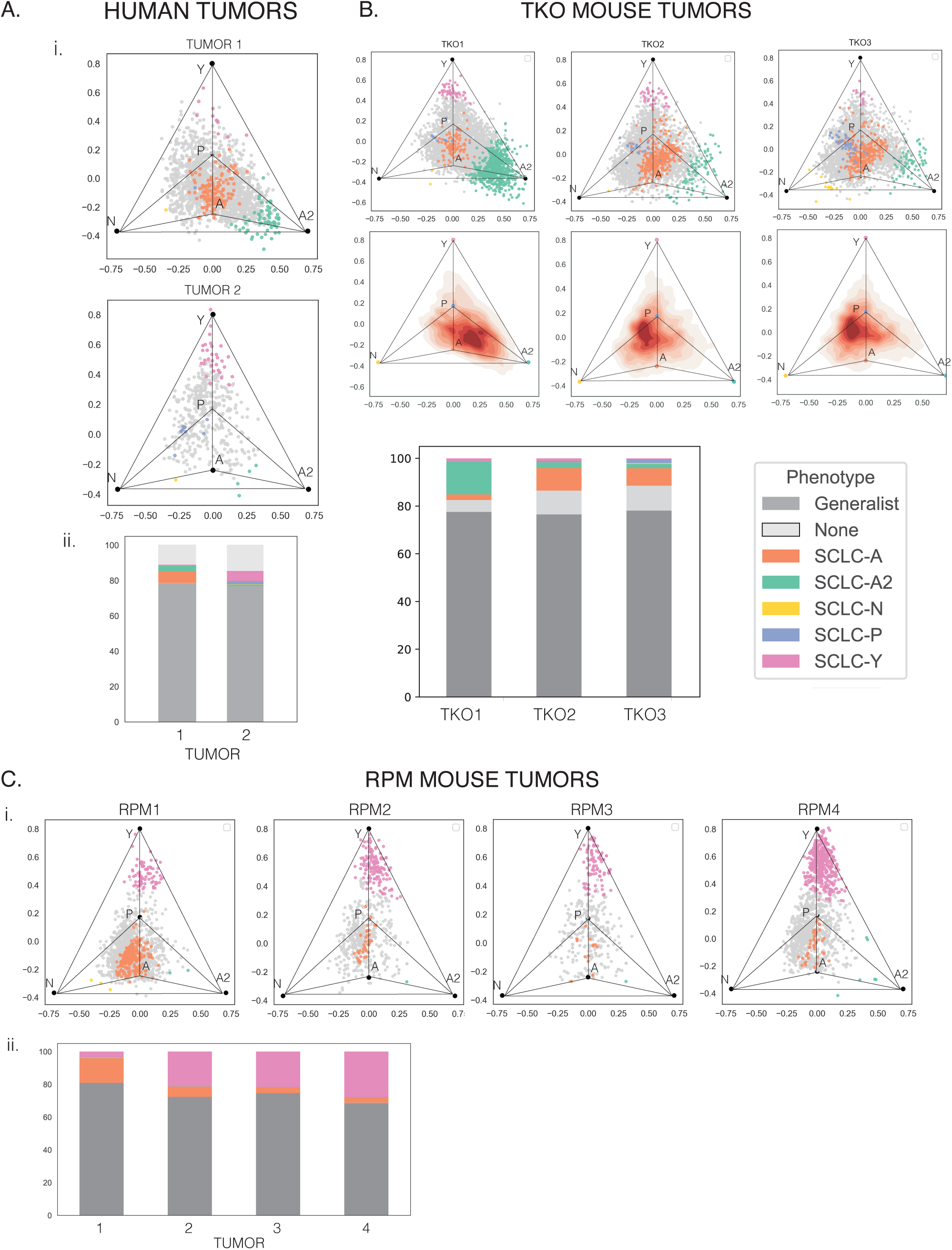
Single Cell Subtyping on Human and Mouse Tumors. A. Similar to Figure 3C, LLE plots for each human tumor. Stacked bar plots show overall composition of each sample, including generalist and unclassified cells. The human tumors show distinct subtype composition; Tumor 1 comprises SCLC-A and SCLC-A2 cells and NE generalists, while Tumor 2 is mostly SCLC-Y cells and non-NE generalists. These two tumors demonstrate the high level of inter-tumoral and intra-tumoral heterogeneity of SCLC. B. LLE plots for each TKO tumor. These three tumors are largely similar, with varying proportions of SCLC-A, SCLC-A2, and SCLC-P specialists. TKO1 has the largest proportion of SCLC-A2 cells and the fewest unclassified cells. TKO2 and TKO3, which are derived from the same mouse, have very similar profiles, with slight differences in the proportion of SCLC-N, SCLC-P, and SCLC-Y cells. C. LLE plots for each RPM tumor. Each RPM tumor is composed of SCLC-A and SCLC-Y specialists, with varying proportions of generalist cells. RPM1 has the largest proportion of SCLC-A cells, concordant with results from the original paper suggesting this tumor falls early in pseudotime inferred from the trajectory between SCLC-A, SCLC-N, and SCLC-Y.^47^ RPM 2-4 have more SCLC-Y cells, which is expected for tumors that fall later along the same trajectory.

In primary (Triple Knockout Models TKO1 and TKO2) and metastatic (TKO3) mouse tumors from the Sage laboratory,^28^ archetype signatures revealed a large proportion of SCLC-A2 subtype cells in TKO1. In contrast, TKO2 and TKO3, originated in the same mouse, had higher proportions of SCLC-A (**Figure 4B)**. Each tumor had a sizable portion of non-neuroendocrine cells as well. We next applied our analysis to Myc-driven mouse tumors (Rb1^fl/fl^; Trp53^fl/fl^; Lox-Stop-Lox [LSL]-Myc^T58A^, RPM) **(Figure 4C)**. The specialists in RPM tumors, in contrast to TKO tumors, were predominantly SCLC-Y, with varying proportions of SCLC-A cells. In all mouse tumors analyzed, regardless of relative archetype composition, a large proportion of cells were generalists. This suggested that intermediate cell states are a staple of GEMM tumors, however generated, further supporting the notion of a cell-state continuum.

Taken together, single-cell gene expression data indicated that SCLC cell lines, human tumors, and GEMMs each formed a continuum of expression at the single-cell level, rather than as mixes of discrete (specialist) subpopulations near the archetypes.

### Novel plasticity metric, CTrP, captures potential for cell state transitions

A continuum of phenotypes often arises when cells traverse diverse regions of gene expression space to adaptively fulfill function.^38, 44, 55, 78, 79^ In this dynamic continuum, intermediate generalists may represent either stem-like cells acting as a source for specialists at archetype vertices or cells in transition from one archetype vertex to another. To determine which SCLC cells can traverse the phenotypic continuum, or whether there exists a specific stem-cell like state from which all traversing cells originate, we quantified the plasticity of individual cells. Briefly, we computed RNA velocity^80^ by modeling spliced versus unspliced counts of individual genes using scVelo^81^ **(Supplemental Figure 8A)**. This assigned a velocity vector to each individual cell in gene expression space, where each dimension (and thus each component of the vector) is a gene. To visualize cell state dynamics, we projected the velocity vectors into archetype space using the same method for single cell gene expression profiles described above. Based on average splicing kinetics, the time-step between measured gene expression and extrapolated expression based on the velocity is around 3-4 hours.^80^ To extrapolate beyond this, we used RNA velocity and the relative locations of sampled data points to generate a transition matrix that defines a Markov Chain model, where each sampled point is a state in the model (**Supplemental Figure 8B)**. Finally, we inferred plasticity by calculating an expected distance of transition for each single cell, which we called Cell Transport Potential (CTrP, **Supplemental Figure 8C**).

CTrP should be zero for steady states (or end states) in the system, and higher for cells that are able to change their gene expression profile by a large amount to reach a steady state. In this way, CTrP should capture the average (expected) change in gene expression each cell is capable of, a proxy for the height (potential) of a cell state in a phenotypic landscape. By averaging potential transitions for each cell, we are necessarily deconvoluting two distinct characteristics of plastic stem-like cells: (1) the average change in expression expected increases with stemness, which we term CTrP, and (2) the variance in possible cell state transitions increases with stemness, which we term pluripotency and is outside the scope of the manuscript (see Methods and Discussion for more).

We tested this metric in two biological differentiation systems where the ground truth is known, as highlighted in Supplemental Methods (**Supplemental Figure 9**). In both systems, CTrP scoring was capable of capturing potential for cell state transitions, with classic stem cells and lineage progenitors having high plasticity and terminally differentiated cells showing low or no plasticity.

### CTrP quantifies high plasticity of generalists and SCLC-Y cells in SCLC cell lines

We then turned to analyzing cell state dynamics within human cell lines using our CTrP analysis. We found that the RNA velocity field of each sample varies widely, suggesting that each cell line we measured contained different proportions of transitioning and phenotypically stable subpopulations **(Figure 5A and Supplemental Figure 10A)**. Some of the high-velocity regions show directed, coherent movement, such as in the cell lines H1048, H841, H524, and DMS454, while the others are more stochastic and less directed. In the cell lines with coherent velocity, we found that only SCLC-Y specialists (in H841 and H1048) consistently moved towards generalists; in other words, Y cells act more like sources for the rest of the population. Conversely, H524 cells trend from intermediate phenotypes towards the SCLC-N archetype, suggesting the N phenotype acts as a sink in this cell line. The cell line populations showed little movement on average across the phenotypic continuum (small black arrows in center figure of **Figure 5A**), suggesting each sample is close to a dynamic equilibrium, where individual cells move within the archetype space but the overall cell density change across the landscape is close to zero. This was expected for relatively stable cell lines grown in controlled environments.

**Figure 5:**
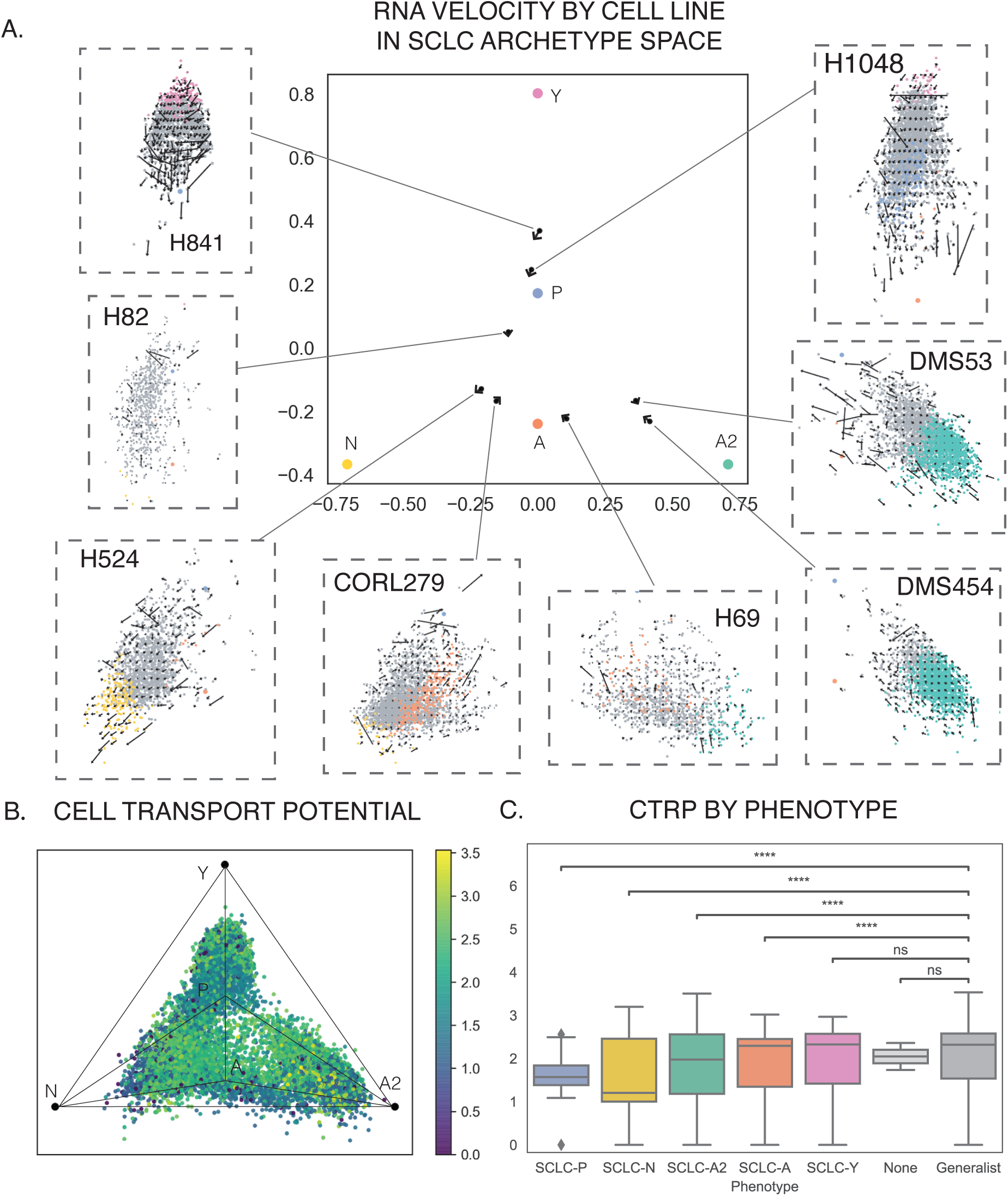
Plasticity of Human Cell Lines. A. RNA velocity by cell line in SCLC archetype space. Using the same LLE from Figure 3, single cells from human cell lines are embedded into archetype space, defined by 5 SCLC archetype vertices. Center plot shows average velocity for each cell line, with length of arrow corresponding to magnitude of the average velocity vector. All cell lines show low average velocity, suggesting little movement of the population within archetype space, consistent with the relative consistency of cell line populations over time. Each cell line is also shown individually, overlaid with a velocity grid that averages velocity within small neighborhoods of the graph. While some cell lines, such as H841 and H1048, show small amounts of directed movement (away from SCLC-Y, in both cases), several other cell lines show little coherence of movement across archetype space. B. Cell transport potential (CTrP) for human cell lines. CTrP is a measure of plastic potential for individual cells based on an RNA-velocity-derived Markov Chain model. Higher CTrP, shown by color of data points in archetype space, signifies a higher expected distance of travel in gene expression space. Overall, SCLC-Y and generalist cells have higher CTrP, though the full range of CTrP values can be found in virtually every region (archetype specialists or generalists) of the archetype space (**Supplemental Figure 10**). C. CTrP by phenotype. Generalists, unclassified cells, and SCLC-Y specialists show significantly higher plasticity than the remaining phenotypes. Overall, plasticity values are relatively low, ranging from around 3.5 at high-plasticity root cells to 0 at absorbing end states that are present in every specialist type (**Supplemental Figure 10**) (t-test, **** p-value < 0.0001, ns = not significant).

When we calculated CTrP, we found that all cell lines spanned degrees of plasticity, with most showing distinct high and low plasticity subpopulations **(Figure 5B and Supplemental Figure 10B and C)**. Cells with high CTrP that are likely to traverse the continuum of states exist in multiple regions in **Figure 5B**, suggesting that there is not a single stem cell phenotype for SCLC cell lines but rather a “stem-like” functional state that can span SCLC archetype space. However, the presence of distinct root cells for most phenotypes (**Supplemental Figure 10B**) showed that it is possible different high CTrP states are subtype-specific progenitors already committed to the cell line subtypes and analyzing cell line data would miss the existence of a true SCLC stem cell state. When we compared CTrP to archetype scores from **Figure 3**, we found a negative correlation between plasticity and archetype scores, suggesting generalist cells with lower archetype scores are more plastic (**Supplemental Figure 10D**). Indeed, generalists (and unclassified cells) were significantly more plastic than all other cell types except SCLC-Y cells, which may reflect SCLC-Y cells’ source-like nature described above (**Figure 5C**). Overall, CTrP analysis of SCLC cell lines showed that: i) generalists across the continuum are likely to be high-plasticity transitioning cells; ii) SCLC-Y specialists may act as a plastic source, if present; and iii) other specialists are more likely to act as end states with low plasticity.

### SCLC tumors show SCLC-Y cells are most plastic *in vivo*

We next applied our plasticity pipeline to the SCLC tumors mentioned previously (**Figure 6**). Each of the two SCLC human tumors showed no easily discernible pattern of RNA velocity in archetype space (**Supplemental Figure 11**). However, the transport potential of each tumor clearly indicated that cells close to SCLC-Y have high CTrP on average compared to the rest of the tumor cells (**Figure 6A**). The clarity gained by our CTrP analysis over RNA velocity alone demonstrated its utility in understanding cancer cell plasticity. Furthermore, these results suggested highly plastic SCLC-Y cells may play a role in tumor relapse by regenerating intermediate generalists that are adaptable to external perturbations. However, this does not preclude the possibility that other human SCLC tumors have different plasticity landscapes.

**Figure 6:**
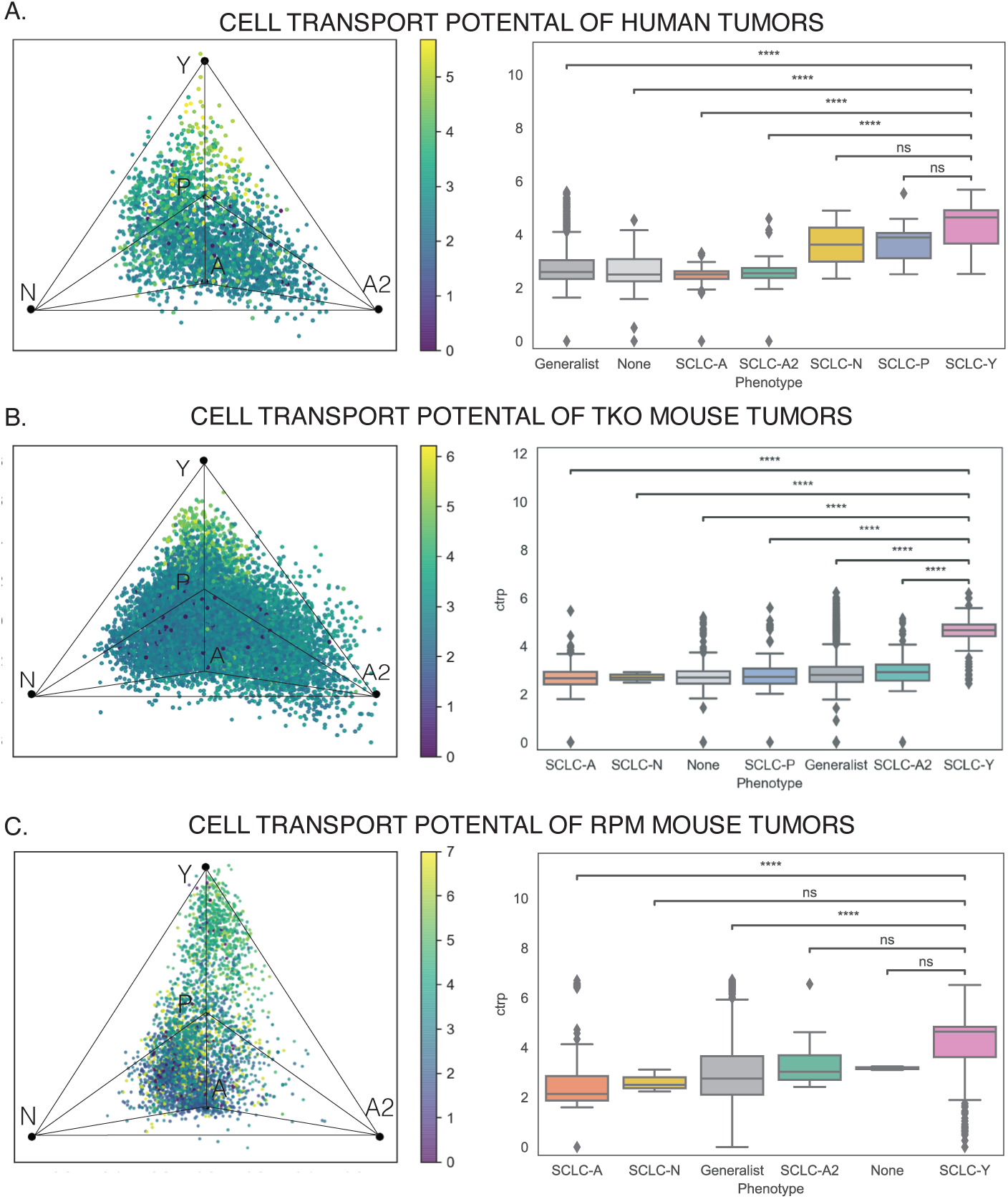
Plasticity of Human and Mouse Tumors. A. CTrP analysis of human tumors. Human tumors show little directed movement across the archetype space according to RNA velocity (**Supplemental Figure 11**), but CTrP analysis shows clear dependence of plasticity on subtype. SCLC-Y cells have higher plasticity than most other cell types, followed by SCLC-P and SCLC-N specialists. B. CTrP analysis of TKO mouse tumors. Similar to human tumors, TKO mouse tumors do not have coherent RNA velocity fields (**Supplemental Figure 12**), but CTrP shows clear dependence of subtype, with SCLC-Y cells significantly more plastic than all other cell types. C. CTrP analysis of RPM mouse tumors. RNA velocity fields show slight movement towards the SCLC-N archetype in some of the RPM tumors (**Supplemental Figure 13**). CTrP analysis reveals that SCLC-Y cells are the most plastic (t-test, **** p-value < 0.0001, ns = not significant).

In GEMMs tumors, RNA velocity also did not report easily identifiable patterns (**Supplemental Figures 12 and 13**). Conversely, in all three TKO mouse tumors, CTrP clearly identified a subpopulation of highly plastic SCLC-Y cells (**Figure 6B**). Similarly, four RPM tumors showed that SCLC-Y specialists have the highest plasticity on average (**Figure 6C**). However, the SCLC-Y specialists ranged from high-plasticity cells to end states with CTrP equal to zero, suggesting that plasticity is not an immutable characteristic of subtype.

In summary, SCLC-Y cells were the most plastic in all tumors and most cell lines analyzed, regardless of source (mouse or human) or genetic background (TKO or RPM).

### MYC drives high-plasticity neuroendocrine SCLC cells towards non-neuroendocrine generalist states through TF network regulation

To determine whether plasticity is a property exclusive to SCLC-Y cells or applies to other subtypes as well, we analyzed a single-cell time-course of RPM progression from Ireland et al^47^ (**Figure 7** and **Supplemental Figure 14**). Early-stage tumor cells from Ad-Cgrp-Cre-infected RPM mice were macro-dissected, placed in culture, and harvested over six timepoints for sequencing. Applying our archetypal signature to the time series data, we predictably found a shift from neuroendocrine subtype cells to non-neuroendocrine, consistent with the results of the original paper.^47^ The plots in **Figure 7A** showed that earlier timepoints (days 4-7) were mostly composed of SCLC-A, -A2, and -N phenotype cells (>50%), which formed a continuum of specialists and generalists. Day 7 was slightly enriched in generalists compared to day 4 and had fewer SCLC-A and SCLC-A2 specialists, which was representative of cells moving away from NE archetypes between these two timepoints. By day 11, cells had transitioned towards the SCLC-Y archetype, with a high proportion of non-NE generalists. We found that, from day 14 on, the cells seemed to be close to equilibrium, with relatively stable proportions of generalists between the SCLC-A, SCLC-A2 and SCLC-Y archetypes, as well as a substantial proportion of unclassified cells that slightly increased with each timepoint (20-30%, **Figure 7B**; interactive 3D plots can be found in the supplement).

**Figure 7:**
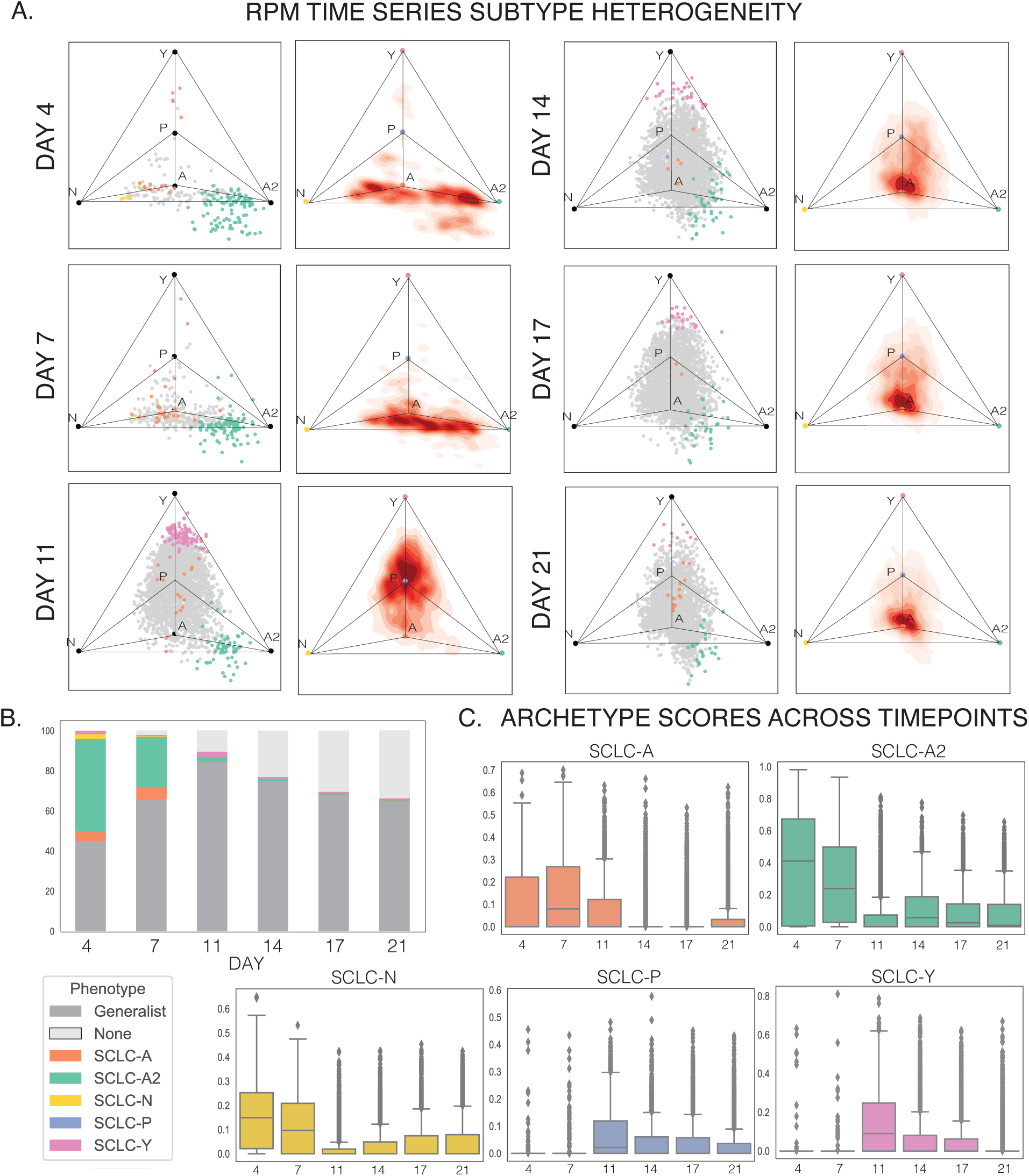
Single Cell Subtyping on RPM Time Series A. LLE plots of archetype scoring for RPM time series. Subtype composition drastically changes over the RPM time course, with early time points dominated by SCLC-A/A2/N mixed (generalist) cells and later time points increasing in SCLC-Y and unclassified cells. Contour plots show a shifting population of generalists that moves from NE regions between SCLC-A, -A2, and -N archetypes towards SCLC-Y, and finally toward a semi-stable equilibrium between SCLC-A and SCLC-Y. B. Stacked bar plots show overall composition change from NE specialists and generalists to non-NE cells, with an increasing proportion of unclassified cells after day 11. C. Archetype scores across time points show that early time points have higher SCLC-A, -A2, and -N scores. Day 11 shows the highest average SCLC-Y scores, consistent with the LLE plots in A. By days 14-21, cells have low but relatively consistent levels of all subtype scores, consistent with the increase in generalist and unclassified cells in B.

This trajectory over the time course could be explained by either cell state transitions and/or selection. If cells were traversing the phenotypic continuum, transitioning from an initial NE phenotype to non-NE phenotypes, we would expect high plasticity in early time-points that decreased across the time course. By contrast, if cells were not transitioning, the shift in subpopulation proportions between day 4 and day 21 (**Figure 7A**) would require that (undetected) non-NE cells from early time points were highly proliferative and able to outgrow NE subpopulations. In this case, we would observe low plasticity for each subtype. To distinguish between these possibilities, we applied our plasticity pipeline. Based on RNA-velocity, we observed significant movement across phenotypic space, particularly around day 11 (**Figure 8A and Supplemental Figure 14**). By calculating plasticity using CTrP (**Figure 8B**), we found that cells from the first two timepoints had the highest plasticity, and plasticity decreased over the six time points with end states only found in the last three timepoints. We therefore conclude that cell state transitions caused the shift in subpopulation composition from day 4 to day 21. Furthermore, these analyses indicated that plasticity is a property that can be acquired by virtually any subtype, since in the RPM time series SCLC-A2, SCLC-N, and NE generalists in early time points had high CTrP (**Figures 7A and 8B**), while SCLC-Y cells were the most plastic in the other tumors analyzed (**Figure 6**).

**Figure 8.**
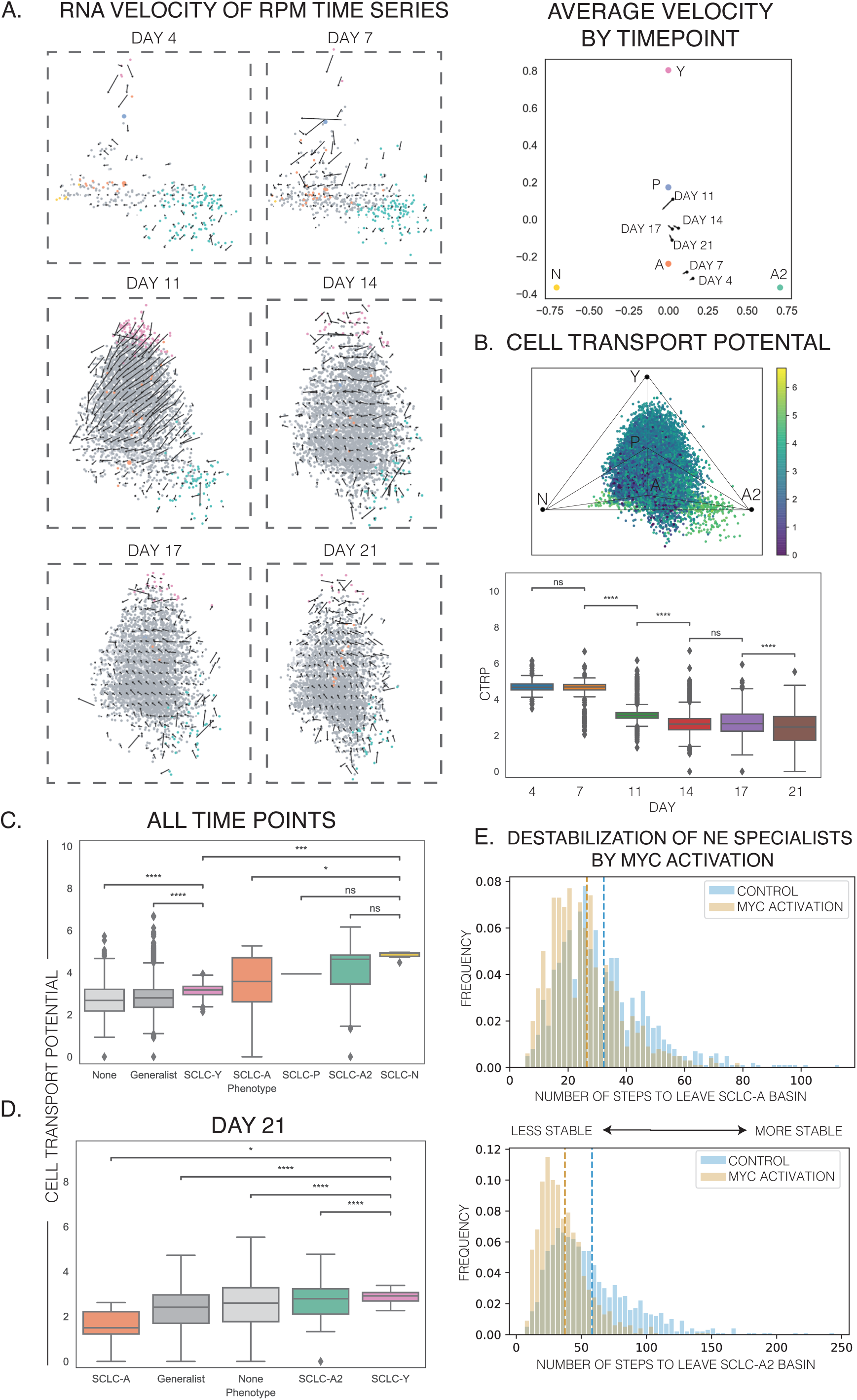
A. (Left) RNA velocity of RPM time series data. In the first two timepoints (days 4 and 7), the cells do not show significant directed movement across archetype space. By day 11, gene expression for a large proportion of the population of cells increases in the SCLC-N signature, shown as movement toward the SCLC-N archetype. Days 14-21 show changes to the direction of this average movement, with velocity that is redirected towards the SCLC-Y archetype by the end of the time series. (Right) Average velocity of time series by time point. Days 4 and 7 show small average velocity vectors from SCLC-A2 towards SCLC-N, which greatly increases by day 11. Day 11 shows the highest average velocity of any time point. Days 14 to 21 show less average movement directed towards the SCLC-Y archetype. B. (Top) Cell transport potential (CTrP) for all cells across RPM time series. Cells closer to neuroendocrine archetypes A and A2 have higher plasticity, which correspond to earlier time points. (Bottom) CTrP decreases over time, consistent with cells that transition from earlier neuroendocrine phenotypes with high plasticity to later non-neuroendocrine phenotypes with low plasticity. End states, where CTrP is 0, arise in the last three time points and tend to fall between A, A2 and N as generalist cells (**Supplemental Figure 15C**). Plasticity does not significantly change between days 4 and 7 and days 14 and 17 (t-test, **** p-value < 0.0001, ns = not significant). C. CTrP by phenotype of all six time points combined. SCLC-N and SCLC-A2 cells have the highest plasticity on average. SCLC-Y cells have significantly lower plasticity than these neuroendocrine specialists, and generalists and unclassified cells show the lowest plasticity on average, though generalists also show the greatest variability (t-test, * p-value <0.05, *** p-value < 0.001, **** p-value < 0.0001, ns = not significant). D. When restricted to cells from the last time point (day 21), the plasticity of each phenotype is more similar to *in vivo* RPM tumors (Figure 6C). SCLC-Y cells in the last time point show significantly higher plasticity than other cell types, with SCLC-A cells showing the lowest CTrP values on average. E. *In silico* destabilization of neuroendocrine specialists by MYC activation. Using BooleaBayes simulations, we determined the effect of MYC activation on the stability of different phenotypes. By counting the number of steps a random walk takes to leave the basin of attraction for a specific state, we can quantify the destabilizing effect of any perturbation. We repeated the random walks for 1000 iterations to generate a distribution of the number of steps it takes to leave each basin. Fewer steps to leave suggests that the perturbation has a destabilizing effect, while more steps to leave suggests a stabilizing effect. As shown, SCLC-A and SCLC-A2 states are significantly destabilized by constitutive MYC activation, suggesting it is capable of increasing plasticity in RPM tumors.

When we further investigated the relationship between subtype and plasticity, we found that SCLC-N cells showed higher plasticity compared to other cell types (**Figure 8C)**. This may indicate that the SCLC-N state is an unstable cell state, likely existing as a transient intermediate state during the MYC-driven transition from SCLC-A/A2 to SCLC-Y. While SCLC-Y cells had lower plasticity than other specialist states in this time course, we found that generalists and unclassified cells had significantly lower plasticity than SCLC-Y (**Figure 8C**). Therefore, neuroendocrine cells from the first time points, particularly plastic SCLC-A2 and SCLC-N cells, transitioned over the time course to SCLC-Y states and then to generalist states.

Based on this plasticity analysis of RPM tumor cells, we conclude that MYC was capable of influencing the distribution of SCLC cell states by increasing CTrP in neuroendocrine specialists. However, this increase in plasticity for SCLC-A2 and SCLC-N cells was only observed in early time points along the time course. Indeed, by day 21 of the time course, SCLC-Y cells were once again the most plastic subpopulation (**Figure 8D**), which was consistent with the high plasticity seen in SCLC-Y cells in RPM mouse tumors *in vivo* (**Figure 6C**). This may reflect the re-equilibration of SCLC subtypes seen by day 14 (**Figure 7A and B**). Therefore, we conclude that the plastic potential of each phenotype shifted over the time course, stabilizing by day 21 to result in the plasticity landscape seen in RPM tumors *in vivo*.

Finally, we sought a mechanistic explanation for the marked ability of MYC to modulate plasticity of cell states. Because each SCLC archetype corresponds to a semi-stable TF network attractor state,^28^ we reasoned that degrees of plasticity for a specific cell state may primarily vary by modulation of TF-TF interactions in the network. Unlike transient upregulation of MYC, RPM tumors constitutively express MYC, and we therefore hypothesized that RPM tumors exhibit modified TF-TF interactions in the network and shifted dynamics of cell state transitions from the human cell lines, human tumors, and TKO mouse tumors. We therefore introduced MYC into a previously derived SCLC network and applied our BooleaBayes^28^ algorithm to determine its effect on subtype stability (See Methods and **Supplemental Figure 16**). Constitutive MYC activation destabilized the SCLC-A and SCLC-A2 states, decreasing the number of steps it takes to leave those attractor’s basins (**Figure 8E**). This result indicated that SCLC plasticity could be modulated at the epigenetic level. Specifically, modifications of network dynamics, e.g., by MYC, could increase the plasticity of neuroendocrine specialists such that these cells transition towards non-neuroendocrine generalists.

## Discussion

SCLC is a heterogeneous cancer comprising neuroendocrine and non-neuroendocrine subtypes, classified by eponymous transcription factors. In this study, our goal was to understand dynamics amongst the canonical subtypes A, A2, N, P and Y, since plasticity is likely to play a key role in supporting SCLC aggressive features.^26, 28^ However, in analyzing SCLC datasets from diverse sources we found that the current subtype classification is insufficient to fully capture SCLC phenotypic heterogeneity, because large numbers of SCLC single cells, regardless of source (cell lines or human and mouse tumors), fall in-between subtype clusters (**Figure 2A, 3, 4**). Furthermore, a discrete subtype space hinders analysis of continuous dynamics between SCLC single-cell phenotypes, which may take place during progression, drug response and relapse. Here, to overcome limitations of discrete subtype clustering, we proposed an alternative view of SCLC heterogeneity, based on SCLC archetypes that define a phenotypic continuum (**Figure 1**). While correspondence between gene signatures of archetypes and canonical subtypes was largely maintained, the archetype-bounded phenotypic continuum paradigm presented several advantages that better represent SCLC cell state space: i) the transcriptional profile of each single cell can be evaluated based on distance from archetype gene signatures (e.g., a cell between archetypes N and Y is assigned a properly apportioned intermediate N/Y phenotype); ii) it allows us to quantify plasticity of intermediate phenotypes, by tracing transition paths between archetypes and identifying regions of high SCLC cell plasticity; iii) cell transitions are rooted in multi-objective evolutionary theory such that movement across the continuum fulfills the goal of trading off between cancer hallmarks tasks, providing a functional interpretation of SCLC phenotypes; and, iv) the continuous framework allows us to identify TFs that drive plasticity and suggest strategies for targeting plastic SCLC cells, which we propose should be a high priority for effective SCLC treatment.

Our analysis, inspired by the work of Alon and co-workers,^40, 44, 45, 55, 82^ used a novel pipeline to quantify the role of heterogeneity and plasticity in SCLC tumors, which can easily be applied to new samples and datasets as they become available. In previous methods for subtyping SCLC, single cells that did not closely adhere to subtype signatures (e.g., co-expressing multiple eponymous TFs such as ASCL1 and NEUROD1,^31^ or expressing none of them) were difficult to classify. In contrast, archetype analysis easily described these single cells, positioning them in the phenotypic continuum at appropriate distances from the relevant archetype vertices. In this continuum of gene expression space, cells closest to a vertex can be considered specialists, i.e., performing the specific functional tasks that gene ontology enrichment assigns to that archetype. Cells located between archetypes can be considered generalists, i.e., performing multiple tasks sub-optimally. Our results suggested that both SCLC cell lines and tumors comprise specialist and generalist cells, with cells traversing the continuum to optimize fitness of the population. It is tempting to speculate that such levels of adaptability may be responsible for the extremely aggressive features of SCLC tumors, as mentioned below. For instance, an altered balance in favor of the more adaptable generalist cells may be expected in tumors immediately after treatment, a scenario that could be tested experimentally in GEMM or CDX tumors. These dynamics may begin to explain the exceptional initial response to standard of care in SCLC tumors (as specialist cells may succumb to treatment), inevitably followed by relapse (driven by generalist cells).

Using gene set enrichment analysis, we identified the tasks optimized by each specialist cell type, such as vasculogenesis for archetype Y, proliferation for archetype A, and inflammation for archetype A2. This palette of biological tasks is in agreement with recent reports indicating that lung tumors are capable of building their own microenvironment, as well as with the low amounts of non-tumor cells usually found in SCLC tumors compared to other tumor types.^8, 26, 83, 84^ A specific example is trophic support provided by archetype Y cells via vasculogenic mimicry, which was recently described in experimental CDXs.^85, 86^ Overall, the high proportion of adaptable generalists we found across SCLC samples is consistent with the aggressiveness of SCLC and may partially explain its ability to rapidly invade and metastasize, by quickly adapting and thriving in inhospitable microenvironments.

An unanswered question in the field is whether SCLC phenotypes are plastic, i.e, capable of state transitions.^11^ The dispersal of SCLC cells across the phenotypic continuum suggests that cell state transitions occur, and therefore we interrogated cell dynamics using RNA velocity.^80^ We defined a novel metric, Cell Transport Potential (CTrP), which quantifies the expected change in transcriptomic state for a cell in a dynamic population. This metric is akin to the height of a cell state in a phenotypic landscape, relative to cells in the system that are not predicted to change cell state at all (endpoints with CTrP = 0). Another form of plasticity that is often convolved with potential is the concept of pluripotency, which is akin to the local depth of the basin in which a particular cell state resides. In other words, CTrP describes how “far” a cell is likely to go in phenotype space (here, defined by gene expression), while pluripotency describes the ability to reach multiple different endpoints. Further analysis is needed to quantify pluripotency, which will most likely require additional experiments such as lineage tracing in scRNA-seq. We may find upon further analysis that cells can possess high plasticity of one type (e.g. CTrP) and not the other (e.g. pluripotency). For example, it may be possible that early SCLC-A/A2 cells in the RPM time course have high CTrP but later SCLC-Y cells have increased pluripotency.

CTrP is dependent on the underlying genetics that determine the shape of the phenotypic landscape, the particular cellular state in which a cell resides, and any external conditions that may transiently “tilt” the landscape. For example, in the RPM time series, it may be the case that MYC promotes plasticity of SCLC-A/A2 cells to non-neuroendocrine states, such that early states have high CTrP, but only under the specific condition of constitutive MYC expression. We argue here that plasticity is not an immutable quality of a location in phenotypic space (i.e., a gene expression profile), but a functional state that may, for example, characterize the SCLC-Y archetype in treated human tumors and characterize the SCLC-A/A2 archetype in *ex vivo* RPM tumor progression. While SCLC-Y cells had high plasticity in all of the tumors tested, further analysis is needed to clarify how they acquired this characteristic. It may be that SCLC-Y cells acquire stemness that is used by the tumor to invade surrounding tumors and subsequently transdifferentiate to seed metastases. Appropriate experimentation should be able to clarify this important point, which negates the existence of a single, immutable stem-like cell type in SCLC tumors, in favor of “stemness” as a functional state that can be adopted by any cell. Recently a PLCG2-expressing stem-like subpopulation was reported in a survey of human SCLC tumors.^87^ This stem-like cell may not be inconsistent with a diverse, stem-like functional state, since it was reported to be present across A, N, and P subtypes. Similarly, previously identified tumor propagating cells (TPCs) common to all SCLC tumors may fall into this stem-like functional state.

We speculate that the plasticity of SCLC cells may derive from dysregulation of the innate plasticity in normal PNECs, believed to be the cell of origin for SCLC.^1, 2, 88^ After injury to the lung epithelium, PNECs can de-differentiate to perform repair tasks and regenerate club cells.^3, 89–91^ This ability to reversibly (de)differentiate under injury suggests that non-genetic mechanisms allow cells to traverse phenotypic space from specialist PNECs, whose main task is chemosensory hypoxia detection and response,^92–95^ to generalist transit-amplifying populations,^89, 91^ to specialist club cells, whose main task is secretion of protective proteins.^96–100^ SCLC cells can likewise transition from neuroendocrine to non-neuroendocrine subtypes through generalist phenotypes, as shown in the tumors analyzed here. Furthermore, in SCLC tumors and cell lines, generalist cells exhibit variable plasticity levels: in all cases, generalists span high plasticity states and low plasticity end states, and the average level of plasticity of generalists in each case varies with respect to that of specialist cells (**Figures 5 and 6**). SCLC generalist cells therefore appear to be a highly adaptable cell state, poised to become plastic in response to perturbations, such as treatment or changes in microenvironmental conditions. These cells may therefore correspond to the transit-amplifying population of de-differentiated PNECs that arise after club cell injury.^91^

The current standard-of-care for SCLC, i.e., chemotherapy and/or radiation, is predicated upon targeting highly proliferative cells. However, this treatment inevitably results in resistant relapse. The large amount of highly plastic cells we detected in SCLC cell lines and tumors (**Figure 5, 6, and 8**) suggest that plasticity may drive resistance in SCLC. The phenotypic continuum also shows that plasticity enables SCLC cells to trade off cancer hallmark-related tasks, which translates to high level of adaptability to diverse microenvironments. Thus, plasticity may also be responsible for SCLC aggressive traits, such as local invasion and early metastatic spread. Plasticity itself, then, should be considered as an effective target for SCLC treatment. For example, we found that human tumors have plastic SCLC-A and SCLC-A2 cells that can transition to a non-neuroendocrine subtype. A strategy to restrict this movement of SCLC-A cells could be silencing MYC, a destabilizing factor for ASCL1-driven phenotypes (i.e., it increases their plasticity) previously implicated in SCLC acquired resistance. We therefore propose a paradigm shift for SCLC treatment towards targeting plasticity. Inhibition of phenotype switching has been suggested as a target for therapy in other cancer types, such as breast cancer and melanoma.^20, 71, 101–104^ For SCLC, we point to epigenetic strategies that can be derived from analyses of TF network dynamics, such as the MYC inhibition strategy proposed in **Figure 8**. Given the primary role of TFs in driving SCLC phenotype, as well as transitions across the phenotypic continuum, SCLC should be a prime candidate for epigenetic therapy targeted to plasticity.

## Methods

### Bulk SCLC Cell Line RNA-seq Data Preprocessing

Bulk RNA sequencing expression data on SCLC cell lines was taken from two sources: 50 cell lines were taken from the Cancer Cell Line Encyclopedia,^105, 106^ and 70 cell lines (not including H69 variants) were taken from cBioPortal^107, 108^ deposited by Dr. John Minna (2017). 29 cell lines overlapped between the datasets, so a “c” (CCLE) or “m” (Minna) was used to denote the source of each cell line. Each dataset was filtered and normalized independently and then batch corrected together. For each source, genes and cell lines with all NAs were removed, as well as mitochondrial genes. The counts data is then normalized by library size and transcript abundance to TPM values. The two datasets were combined using overlapping genes and log-transformed, and genes with low expression across all samples were removed (cutoff of log(TPM) >= 1). The two datasets were then batch corrected using the *sva* R package, which includes a COMBAT-based integration method.^109, 110^ SVA, or surrogate variable analysis, uses a null model and a full model to derive hidden variables, such as batch, that may contribute to gene expression variance across samples. The four SCLC TF factors that define broad subtypes— ASCL1 (A), NEUROD1 (N), YAP1 (Y) and POU2F3 (P)— were used to align the two datasets to each other. Batch-corrected distributions of gene expression are shown in **Supplemental Figure 1**. The resulting dataset contained 120 samples and 15,950 genes.

For labelling cell lines by subtype cluster (**Supplemental Figure 2A**), hierarchical clustering with the Spearman distance metric was calculated using the R function *hclust* and cluster cutoffs were determined manually. Cell lines previously characterized as SCLC-A^28^ comprised two branches of the dendrogram separated by SCLC-N cell lines, most likely due to the dual positivity of ASCL1 and NEUROD1 in some SCLC-A cell lines. Cell line H82 (from both data sources) was considered “unclustered,” as it has previously^28^ been considered an SCLC-N cell line (with positive expression of NEUROD1) but was clustered with SCLC-Y cell lines here. As shown in **Figure 3A**, H82 falls in between SCLC-Y and SCLC-N archetypes, corroborating our conclusion that clustering cannot adequately describe all cell lines.

To quantify the goodness of fit for clustering, internal validation metrics were computed using the R package *clValid* (**Supplemental Figure 2B**).^111^ Data was clustered into k=[2,6] clusters using *hierarchical*, *k-means*, and *pam* methods. Optimal scores for the Dunn Index were calculated with two clusters using hierarchical clustering. Because we know from previous literature that more than two SCLC subtypes exist, this suggests that the additional subtypes are not well separated.

Principal Component Analysis (PCA) was run on the bulk RNA-seq dataset using the R *prcomp* function. The R package *factoextra* was used to visualize percentage explained variance for each principal component up to 30. The elbow method on explained variance per component was used to choose 12 principal components for downstream analysis **(Supplemental Figure 2C)**. The top 12 principal components were able to explain ∼50% of the variance in the dataset, suggesting a low-dimensional representation of the data was possible.

### Archetypal Analysis using PCHA

Using an AA method known as Principle Convex Hull Analysis (PCHA),^37^ we found a low-dimensional Principal Convex Hull (PCH) for the dataset. The convex hull is the minimal convex set of data, which can be thought of as a rubber band that “wraps” around the dataset (**Supplemental Figure 3A, top**). The PCH is a subset of the convex hull that comprises a set of vertices, or archetypes, still able to capture the shape of the data. By comparing the convex hull to the PCH, one can determine how well a low-dimensional shape fits the dataset, and thus ascertain the optimal number of vertices, or archetypes, that define the shape.

Archetypal Analysis was done using the Matlab package *ParTI.*^55^ To determine the best k, where k equals the number of archetypes, the function *Parti_lite* was used, varying k from 2 to 12. As suggested by ParTI based on the explained sample variance for each number of archetypes (**Supplemental Figure 2C**), 5 was chosen for k, which is consistent with the current literature on SCLC subtypes. The function *ParTI* was then used for the full analysis with parameters: dim = 12 (dimensions) and algNum = 5 (PCHA^59^) to find the location of the archetypes in gene expression space. Empirical p-values were computed by comparing the ratio of the polytope volume to that of the convex hull (t-ratio), the shape made with the constraint that all data must be contained inside it. This t-ratio is then computed on bootstrapped shuffled samples 1000 times to generate a background distribution to determine if the shape of the data is significantly different from a cloud of points. The p-value is the fraction of random t-ratios that are equal to or higher than the observed t-ratio. This p-value = 0.034, suggesting that 5 archetypes fit the data well. Errors were then calculated on each archetype location by sampling the data with replacement and calculating the archetypes on the bootstrapped data sets (1000 times, **Supplemental Figure 2D**). Error on the archetypes gives an idea of the variance in archetype position expected, and a smaller variance suggests the archetype is robust to outlier samples.

Enrichment of subtype labels was determined using the *ParTI* function *DiscreteEnrichment*. Cell lines were binned into 10 bins according to distance from each archetype. For each subtype label (from hierarchical clustering above), the percentage of labels in the bin closest to the archetype was compared to the percentage in the rest of the data using a hypergeometric test. Enrichment was considered significant, as it was for all five subtypes, if the bin closest to the archetype was maximal for that label and the FDR-corrected p-value for the hypergeometric test was significant (Benjamini-Hochberg, q < 0.1). These results are shown in **Supplemental Table 1**.

### Gene expression and ontology enrichment at archetype locations

To find genes and gene sets enriched by each archetype, we tested the enrichment of each feature on the bin closest to archetypes versus the rest of the data. For each archetype, data was separated into 10 bins according to the distance from the archetype (12 samples in each bin). The *ParTI* function *ContinuousEnrichment* was used to analyze gene expression of all 15,950 genes (**Supplemental Table 5**) and gene sets downloaded from MSigDB on October 14, 2020: Hallmark gene sets (H; h.all.v7.2.symbols) and biological process ontology gene sets (C5; c5.go.bp.v7.2.symbols), as well as Cancer Hallmark Gene Sets **(Supplemental Tables 2-4)**. Gene sets are transformed into a matrix using the function *MakeGOMatrix* such that each row is a sample, each column represents an MSigDB category, and expression values represent the average expression of genes belonging to that MSigDB category. Similar to discrete feature enrichment above, data was binned into 10 bins according to distance from each archetype. The expression of each feature was compared between the closest bin to each archetype and the rest of the data using a Mann-Whitney test (FDR-corrected p-value, q<0.1).

### Gene signature matrix generation

After testing each gene with a Mann-Whitney test as described above, genes that are not maximal close to an archetype or with a p-value higher than 0.05 are considered insignificant and are removed from the analysis. Remaining genes are assigned to the archetype for which the mean difference (log-ratio) of log-transformed gene expression in the closest bin to the archetype compared to rest is highest (**Supplemental Figure 3A, middle**). The matrix (with size [G, n], where G = total number of genes and n = number of archetypes) is populated with the archetype gene expression profiles (i.e. the average location in gene expression space after bootstrapping the archetype analysis). To reduce the size of the gene matrix and choose the most salient genes for each archetype, an algorithm is used to optimize the condition number, or stability, of the matrix (**Supplemental Figure 3A, bottom**). The condition number of the matrix is the value of the asymptotic worst-case relative change in output for a relative change in input. Minimizing this value ensures that genes that do not well-distinguish between the archetypes are not included in the matrix. With genes sorted by mean difference for each archetype, the top g genes are chosen for each archetype, with g ranging from 20 to 200. For each g, the condition number of the matrix is calculated using the Python function *cond* from *numpy.linalg* using the 2-norm (largest singular value, p = 2). The gene signature matrix size with the lowest condition number, which includes g* genes, is chosen. For our archetypes, g* = 21, so the resulting size of the gene signature matrix is [g* x n, n] = [105, 5] (**Supplemental Figure 3B** and **Supplemental Table 6**). This method can be extended to other sorted lists of genes, such as genes sorted by adjusted p-value in an ANOVA test between archetypes. For SCLC, the gene signature included the four consensus TFs: ASCL1, NEUROD1, POU2F3, and YAP1 (**Supplemental Figure 3C**). Myc is shown in **Supplemental Figure 3C** as an example of a gene that has been found to be important for SCLC phenotype characterization but is not included in the gene signature because it does not define a single archetype, with expression that spans several archetypes (SCLC-N, SCLC-P, and SCLC-Y).

### Single cell RNA sequencing: experimental methods for cell lines

Eight SCLC human cell lines from the bulk data above were chosen for single cell RNA-sequencing. We chose two cell lines from each neuroendocrine subtype (A: H69 and CORL279, A2: DMS53 and DMS454; N: H82 and H524) and one cell line from each non-neuroendocrine subtype (P: H1048; Y: H841) (**Figure 2A**). This approximates the distribution of subtypes seen in bulk tumor data, where most tumors are largely neuroendocrine. ^11^ We also aimed to pick cell lines that ranged in their distance from their “assigned” archetype, to better understand intermediate samples as compared to ones close to an archetype location.

In preparation for single cell RNA-sequencing, cells were washed with PBS three times, and then the cells were counted and concentration was adjusted to 100 cells/μL. Droplet-based single cell encapsulation and barcoding was performed using the inDrop platform (1CellBio), with an in vitro transcription library preparation protocol.^112^ After library preparation, the cells were sequenced using the NextSeq 500 (Illumina). DropEst pipeline was used to process indrops scRNA-seq data and to generate count matrices of each gene in each cell.^113^ Specifically, cell barcodes and UMIs were extracted by dropTag, reads were aligned to the human reference transcriptome hg38 using STAR,^114^ and cell barcode errors were corrected and gene-by-cell count matrices and three other count matrices for exons, introns and exon/intron spanning reads were measured by dropEst. Spliced and unspliced reads were annotated and RNA expression dynamics of single cells were estimated by velocyto.^80^

### Single cell RNA sequencing: experimental methods for tumors

The two human SCLC tumors were collected in collaboration with Vanderbilt University Medical Center. Tumor #1 was a relapsed tumor collected via bronchoscopy with transbronchial needle aspiration of a left hilar mass. Tissue was washed in an RBC lysis buffer, passed through a 70 μm filter, and washed in PBS. Cells were dissociated with cold DNAse and proteases and titrated every 5-10 minutes to increase dissociation. Cells were then filtered with a 40 μm filter, washed, and droplet-based single cell encapsulation and barcoding was performed using the inDrop platform (1CellBio), as described above (*Single cell RNA sequencing: experimental methods for cell lines*). The tumor was then sequenced on a BGI MGI-seq. Human tumor #2 was stage 1B SCLC tumor with a mixed large cell neuroendocrine component treated with etoposide and cisplatin and was surgically removed via right upper lobectomy. Tumor was immediately placed in cold RPMI on ice for dissociation. Droplet-based single cell encapsulation and barcoding was then performed using the inDrop platform (1CellBio), as described above (*Single cell RNA sequencing: experimental methods for cell lines*). Cells were prepared for sequencing using TruDrop and sequenced on Nova-seq.

### Single cell RNA sequencing: preprocessing

Single cell RNA-seq counts matrices were analyzed using the Python packages *Scanpy* and *scVelo.*^81, 115^ For the cell lines (**Supplemental Figure 4**), the data was subset randomly to 20% of the original data for computational efficiency, leaving 17298 cells. We batch corrected the cell line samples using scanorama,^116^ with default parameters and the “approximate” solver, because this method can integrate multiple samples even if they do not all share a common subpopulation. Using the function *correct_scanpy* with return_dimred = True, the corrected dataset was used to generate PCA dimensionality-reduced embeddings for visualization. Un-batch-corrected data was used for downstream analyses such as subtyping, neighborhood identification and RNA velocity, and plasticity calculations.

For all of the datasets—8 cell lines (**Supplemental Figure 4**) vs 6 RPM time points & 7 GEMMs (**Supplemental Figure 5**) vs 2 human tumors (**Supplemental Figure 6**)—cells were filtered using *Scanpy’s* filtering and normalization functions with a threshold of 200 minimum genes per cell and 3 minimum cells per gene, and log-transformed. We used *Scanpy’s score_genes_cell_cycle* function to regress out cell cycle effects. For all samples, we predicted cell doublets using Scrublet.^117^

### Bulk gene signature subtyping of single cells

In order to apply the gene signature matrix (**Supplemental Table 6**) to single cells, we first subsetted the data to the genes in the archetype signatures. Due to dropouts, the intersection of genes from the signatures and the single cell data may be less than the full signature (105 genes), and we refer to this intersection as “shared genes.” We scale the gene signature and the single cell gene expression data by the L2 norm for each archetype or cell, respectively, to remove differences caused by different platforms (bulk vs. single cell sequencing). Each archetype’s and each cell’s gene expression vector were therefore scaled to have a length of 1, so that the archetype space has basis vectors of length 1. For RNA velocity analyses, we scaled the velocity vectors by the associated gene expression norm for each cell, so that the velocity vector was scaled the same way each cell’s gene expression vector was scaled. We then used least squares approximation to transform each cell into archetype space by solving *Ax = b*, where *A* is the signature matrix (shared genes *g* by subtypes *s*) and *b* is the single cell matrix (shared genes *g* by cells *c*). This is solved using *numpy.linalg.lstsq* to generate a “pattern matrix” *x* (subtypes *s* by cells *c*). Cells with a maximum signature score greater than 0.5 were considered specialists for that subtype. Cells with a maximum signature score of less than 0.1 (i.e., all five archetype scores are less than this threshold) were considered unclassified. An example in two dimensions is shown in **Supplemental Figure 4D**. These cells with low scores may be due to cells that have transformed into phenotypes far from the defined SCLC archetype space. Notably, the number of unclassified cells in cell lines and GEMMs (tumors) was negligible, while the RPM time series had increasing amounts over time, and the human tumor with an LCNEC component had a large proportion of unclassified cells.

### Visualization of Single Cells in Archetype Locally-Linear Embedding

To visualize subtype scores for individual cells, we use Locally-Linear Embedding (LLE)^118^ using the Python package *scikit-learn*. The LLE model estimates potentially nonlinear global structure using locally-linear fits to the data. For each sample, we use an LLE model with two components and the “modified” method to fit and transform 5 simulated cells, one for each archetype (e.g. [1,0,0,0,0] for an SCLC-A cell, [0,1,0,0,0] for SCLC-A2, etc.), into two dimensions, allowing for an archetype diagram where each vertex is an archetype signature. We use the same model to transform the archetype scores of the data into this space. To transform velocity vectors into this space, we use 5-dimensional archetype scores for velocity (described below) of each cell. Velocity grids are generated by binning the data according to the 2-dimensional LLE locations of the data and averaging over each bin.

### RNA Velocity Calculation and Analysis

We interrogated the dynamics of SCLC cells and tumors by analyzing RNA velocity.^80^ RNA velocity uses a splicing model to predict directionality and magnitude of gene expression change in the near future for each cell sampled. We used the Python package *scVelo* to infer movement through the SCLC phenotypic continuum for each cell line independently.^81^ A neighborhood graph (adjacency matrix) and first-order moments were calculated using scvelo.pp.neighbors and scvelo.pp.moments, respectively. We used ScVelo’s deterministic velocity method with samples grouped independently (using *groupby*) within sets of data where the samples are independent of each other: human cell lines, human tumors, and GEMMs (TKO and RPM tumors), while RPM time series samples were calculated together. This allowed us to compute velocity independently for appropriate samples, while still using the neighbors in each full dataset for downstream analyses. We then computed velocity graphs, confidences (coherence of velocities), and velocity lengths.

### Cell Transport Potential Calculation and Analysis

While plasticity has been discussed in the context of differentiation^89, 119–127^ and cancer^17, 21–23, 25, 47, 71, 120, 123, 128–134^ many times, it has not been rigorously quantified for individual cells. In quantifying plasticity, we wanted to capture the local likelihood of phenotypic transition (that is, change in gene expression profile) for each transcriptional state sampled. Furthermore, we would like to take into account size and variance of the phenotypic change. Colloquially, “plastic” cells, such as stem cells, generally are considered plastic because they have at least one of these characteristics: they are poised to change their gene expression profile by a large amount (differentiation potency), and/or they are able to change into multiple different end states (multipotency). We argue below that these types of plasticity are captured by the theoretic notion of a phenotypic landscape. The phenotypic landscape can be quantified using a population balance analysis, as shown in work from Klein and coworkers.^135^ The fundamental equation for this analysis begins by considering the dynamics of cell density across a landscape as a function of cell flux through that landscape and cell accumulation and loss due to proliferation or death.

Because multiple dynamics could potentially explain the same cell density distribution, we constrain the solution using a set of reasonable assumptions, as is done in Weinreb et al.^135^ For example, we make the assumption that hidden features, or any cell measurements besides transcriptomics such as DNA methylation, do not impact cell fate over multiple state transitions through the landscape. In a more complete model that includes these complementary ‘omics, we would be able to deconvolve multiple superimposed dynamic processes, which could explain the incoherent velocity directionality sometimes seen in RNA velocity analysis.^80, 81, 135^ Instead, we must assume that the transcriptomics state dynamics are fully captured at each state by splicing dynamics at the point of sampling, and that a cell’s fate can be predicted based on its multi-step path through other defined gene expression states.^135–137^ This assumption guarantees that measured transcriptomics can fully encode a probability distribution of cell states, and thus that the process we are modeling is Markovian. As such, the dynamics of our system can be encoded as a biased random walk that includes both cell state stochasticity and deterministic movement through the landscape. We also assume that there are no oscillations, such as cell cycling, which is common in trajectory inference methods.^135^ Weinreb et al. found that useful predictions may still be made with this assumption, particularly in cases where oscillations such as changes in cell cycle genes are orthogonal to the process we are interested in.^135^ This implies that the velocity field is the gradient of a potential function and is the key to connecting velocity inference to the theoretical notion of a potential landscape. This potential can be decomposed into two terms: a “transport” term and a “constraint” term.^135^ The deterministic transport term counteracts sources and sinks in the landscape to keep the cell density in dynamic equilibrium and quantifies the average, and it assumes the stochastic diffusion matrix is zero such that there is no diffusion out of high-density areas in gene expression space. As a proxy for this potential term, we calculate the Cell Transport Potential (CTrP). We consider transport to be an average movement over the landscape that ignores stochasticity. For this reason, CTrP is the expected value for the movement of each individual cell. More formally, it is the expected distance of travel for a cell, weighted by the time spent in each other cell state before absorption (reaching an end state). The method is detailed below.

CTrP is a measure of the average distance a cell may travel according to its RNA velocity.^80^ For each independent sample (untreated or treated), we ran the following pipeline:

1. **Using RNA velocity calculated as described above, and for each category (‘treatment’), compute a Markov Chain Model transition matrix**. This is calculated using an adapted version of ScVelo’s transition_matrix function, in which transition probabilities between each two cells, i and j, is calculated from the velocity graph pairwise. Each entry is a probability describing the likelihood of moving from state i to state j, and each row is the probability distribution of transitions from state i. RNA velocity is compared to distances between other cells to get a pairwise cosine correlation matrix (velocity graph). A scale parameter (default 10) is used to scale a Gaussian kernel applied to the velocity graph, restricted to transitions in the PCA embedding. This transition matrix, T, has dimensions nxn, where n = number of cells. It is then normalized to ensure each row adds to 1 (because each row is the probability of cell i transitioning to any other cell j, which should total 100%. Diffusion for T is scaled to 0 (i.e., ignored).
2. **Calculate absorbing states (end states) using eigenvectors**. Eigenvalues are calculated for the transition matrix. Any eigenvalue *l* = 1 (here, with a tolerance of 0.01), is associated with an end state distribution (eigenvector **v**); i.e., T(v) = *l*v implies that a distribution of states v will not change under further transformation (transitions) from T. If the Markov Chain is an absorbing Markov Chain, it will contain both transient states (t = number of transient states, where T(i,i) < 1), and absorbing states (r = number of absorbing states, where T(i,i) = 1). For every absorbing state in the matrix, there will be an associated eigenvalue/vector pair, with *l* =1, because any initial configuration of states will continue to evolve until every cell has reached an absorbing state. Therefore, the multiplicity of *l* = 1 is equal to the number of end states (absorbing states, or irreducible cycles). The associated eigenvectors **v** thus correspond to the absorbing states in the Markov Chain, within the tolerance of 0.01.
3. **Calculate the fundamental matrix.** In an absorbing Markov Chain, it is possible for every cell to reach an absorbing state in a finite number of steps. Let us rewrite T, the transition matrix, as P:

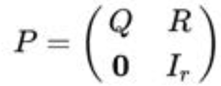

where Q is a t X t matrix, R is a non-zero t x r matrix, 0 is an r X t zero matrix, and I_r_ is an r X r identity matrix. Thus, Q describes the probability of transitioning between transient states, and R describes the probability of transitioning from a transient state to an absorbing state. The fundamental matrix N of P describes the expected number of visits to a transient state j from a transient state i before being absorbed. Because the Markov Chain is absorbing, this number is the sum for all k of Q^k^ (k in (0, inf)): Because Q^k^ eventually goes to the zero matrix (all cells are absorbed), this sum converges for all absorbing chains. Furthermore, each row_i_ of the fundamental matrix describes the expected amount of time spent in state j starting from state i, and thus the row can be thought of as a distribution of weights associated with each state j for each starting state i. N is calculated as written above: the inverse of Q subtracted from the identity matrix. In practice, Numpy’s function numpy.linalg.inv(I-Q) is used to calculate N. If, however, the determinant of I-Q = 0, which occurs when I-Q is a singular matrix, the pseudoinverse is instead calculated using numpy.linalg.pinv(I-Q).
4. **Calculate a distance matrix.** A distance matrix D (n X n) is then calculated using scipy’s function scipy.spatial.distance.cdist. Here, we calculate the Euclidean distance on the PCA embedding of each sample. Because PCA is a linear transformation, it preserves the Euclidean distance metric between states while reducing extraneous noisy dimensions, increasing the efficiency of the calculation. Distance may also be calculated directly on nonlinear dimensionality reduction techniques, such as UMAP and tSNE, but these distances tend to break down for samples that are highly discontinuous (discrete clusters) and should only be applied to continuous data that falls on a single manifold.
5. **Calculate Cell Transport Potential**. Finally, CTrP is calculated as the inner product of fundamental matrix N, and the distance matrix D. This gives an expected distance (sum of distances to j from i, weighted by time spent in j before absorption).

The advantage of this metric over similar techniques, such as pseudotime and other trajectory inference metrics, is that CTrP is an expected distance in linear (PCA) space, which can be compared across samples (assuming they have been embedded in the same PCA). In other words, unitless measures such as pseudotime cannot be compared between samples because their scales are arbitrary; alternatively, CTrP has a scale set across all samples, dependent only on transformations of the data itself (such as log-normalization and PCA).

### Testing CTrP as a Metric for Plasticity

To test this metric, we applied it to two different well-studied differentiation datasets: two developmental time points (P12 and P35) of the dentate gyrus of the hippocampus^138^ and embryonic day 15.5 of endocrinogenesis.^139^ In the dentate gyrus, RNA velocity clearly shows continuous differentiation pathways from radial glia-like cells through neuronal intermediate progenitor cells (nIPC) to some of the cell types, such as neuroblasts. Other differentiated cell types, such as oligodendrocytes (OL), are disconnected from the main “island” in the UMAP. Using our CTrP analysis, we can place these cells in order of plasticity to find that radial glia-like cells and astrocytes have very high plasticity; and microglia, OL, and mature granule cells have low plasticity. This corresponds well with the differentiation program found in Hochgerner et al.^138^ Radial glia-like cells are neuronal stem cells; neuroblasts and nIPCs were identified as transient cell types, which is concordant with their intermediate CTrP levels; and fully mature granule cells show low plasticity. Interestingly, astrocytes showed high plasticity in this analysis, which has been confirmed in several other studies and may be due to the immaturity of the astrocytes in early development sampled here.^138, 140, 141^ For the pancreatic cell differentiation dataset, we found that ductal/endocrine progenitors have high plasticity, as expected, and beta cells represented a terminal state with significantly lower CTrP values (**Supplemental Figure 9**). While alpha, delta, and epsilon cells are also differentiated cell types, our analysis shows these cell states have slightly higher plasticity than beta cells, consistent with the original findings in the paper that at stage E15.5 the majority of progenitors showed velocities toward beta cells. Furthermore, alpha cells may be capable of serving as progenitors for beta cells as well.^142–144^

### Single Cell Network Inference using BooleaBayes

In order to explain our expanded dataset analyzed here, we updated the network structure from Wooten et al. to include MYC and NEUROD1. The updated network structure included 2 new TFs with 43 new edges between them, derived from ChIP-seq databases as detailed previously. Methods for predictions of regulators, stability, and reprogramming strategies are detailed in Wooten et al.^28^

## Supporting information

Supplemental 3D RPM Time Series Figures

## Acknowledgments

We thank members of the Quaranta laboratory for useful suggestions; members of the Lopez laboratory (Vanderbilt University) for critical feedback and support with computation; and Wade Iams, Christine Lovly and the Lovly laboratory, and Jonathan Lehman (Vanderbilt University Medical Center) for their clinical expertise and human tumor preparation for sequencing.

## Conflict of Interest and Funding Sources

VQ received funding from National Institutes of Health National Cancer Institute U54CA217450 (https://csbconsortium.org/) and NIH NCI U01-CA215845 (https://csbconsortium.org/). SMG received funding from NSF fellowship DGE-1445197 (https://www.nsfgrfp.org/). TGO received funding from National Institutes of Health National Cancer Institute 5U01CA231844-03 and 5U24CA213274-04. WTI was supported by the National Institutes of Health (NIH) and National Cancer Institute (NCI) Vanderbilt Clinical Oncology Research Career Development Award (VCORCDP) 2K12CA090625-17, an American Society of Clinical Oncology / Conquer Cancer Foundation Young Investigator Award, and a National Comprehensive Cancer Network Young Investigator Award. CML was supported by a Lung Cancer Foundation of America/International Association for the Study of Lung Cancer Lori Monroe Scholarship and the National Institutes of Health / National Cancer Institute [grant numbers U54CA217450-01, U01CA224276-01, P30-CA086485, UG1CA233259].

CML is a consultant/advisory board member for Pfizer, Novartis, AstraZeneca, Genoptix, Sequenom, Ariad, Takeda, Blueprints Medicine, Cepheid, Foundation Medicine, Roche, Achilles Therapeutics, Genentech, Syros, Amgen, EMD-Serono, and Eli Lilly and reports receiving commercial research grants from Xcovery, AstraZeneca, and Novartis.

## Supplemental Figures

**Supplemental Figure 1:**
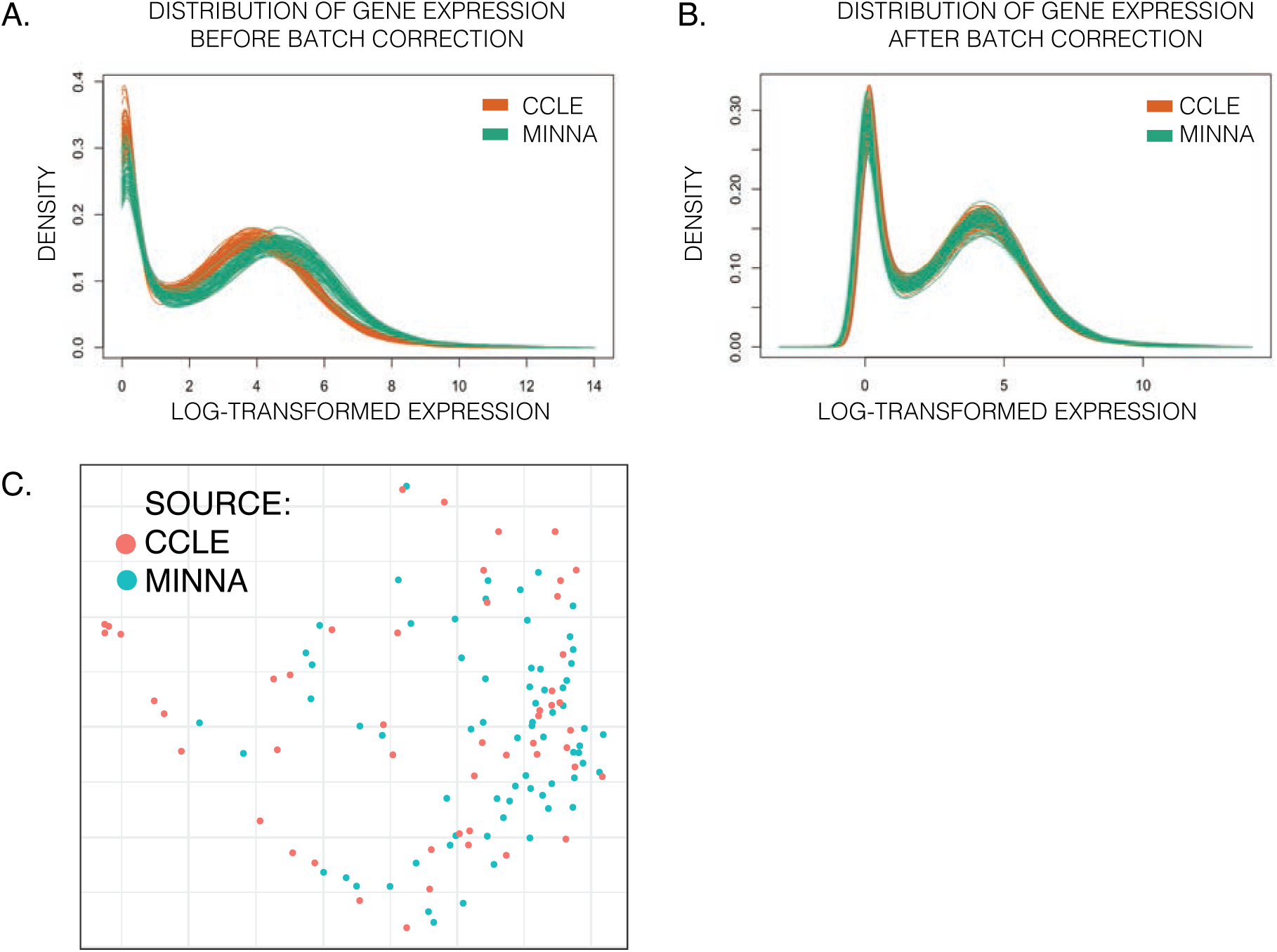
Supplement to Bulk RNA-seq Preprocessing. A. Gene expression distribution before batch correction … B. … and after batch correction for CCLE and Minna datasets. C. Cell line source in PCA.

**Supplemental Figure 2:**
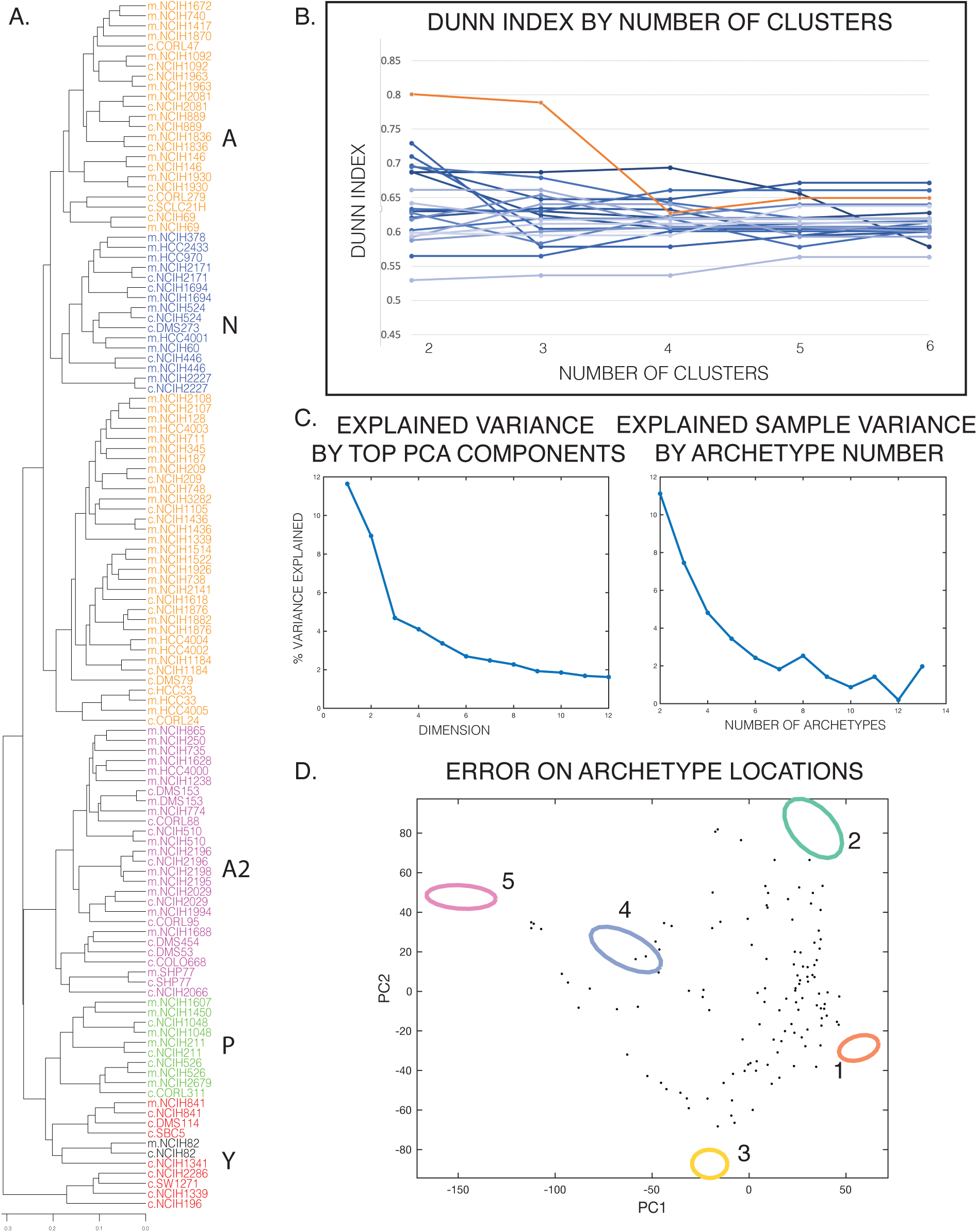
Supplement to PCA and Archetype Analysis. A. Cell line clustering by hierarchical clustering (HC). B. Dunn Index calculated from internal validation tests. Dunn Index is a measure of the minimum inter-cluster Euclidean distance relative to the maximum intra-cluster Euclidean distance. A higher score signifies well-separated clusters. Dunn Index for hierarchical clustering with number of clusters *k* between 2 and 6 is shown for the cell line data (orange line) and twenty randomized datasets of the same size (blue lines). As shown, five clusters for the SCLC cell lines are comparable to that of randomized data, suggesting the clusters are well-separated. C. Explained variance by PCA components and archetypes. (Left) Explained variance by PCA components was used to determine the optimal number of components for downstream analyses. Here, 12 components were chosen based on the elbow of the curve. (Right) Explained sample variance by archetype number shows the amount of variance in the data that can be explained by different numbers of archetypes, where each data point is then some composition of archetype gene expression profiles. Five archetypes were chosen as suggested by *ParTI*. D. Error on archetype locations calculated using a leave-one-out method on the dataset. Relatively small error ranges suggest that the locations of the five archetypes are not dependent on outliers in the data.

**Supplemental Figure 3:**
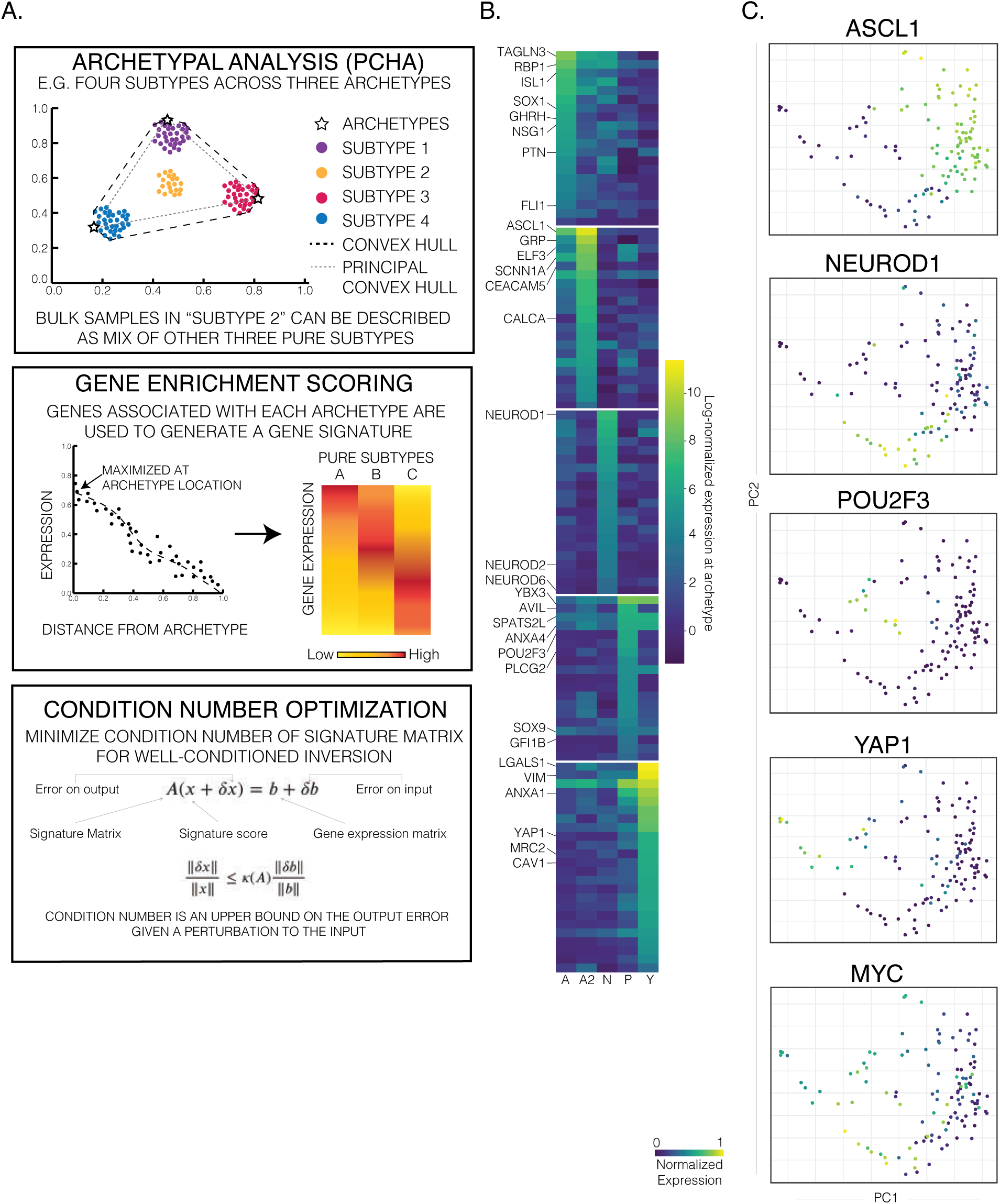
Archetype Analysis and Signature Generation A. Pipeline for AA and signature generation. Archetype analysis is used to define archetype vertices that describe the shape of SCLC cell line RNA-seq data. Gene enrichment at each archetype is used to order genes according to importance in defining archetypes. Condition number optimization determines the best number of genes to include in the archetype gene signature. See Methods for more details. B. Archetype Gene Signature for SCLC. Select genes are highlighted; all genes are reported in **Supplemental Table 6**. C. PCA plots of expression of specific genes (ANYP + MYC). Notably, ASCL1 and NEUROD1 expression overlap on the right side of the figure, and MYC expression is too dispersed to be considered a marker for a specific phenotype. Regions of high YAP1 and POU2F3 expression values are relatively compact, specific to a small region of gene expression space for human cell lines.

**Supplemental Figure 4:**
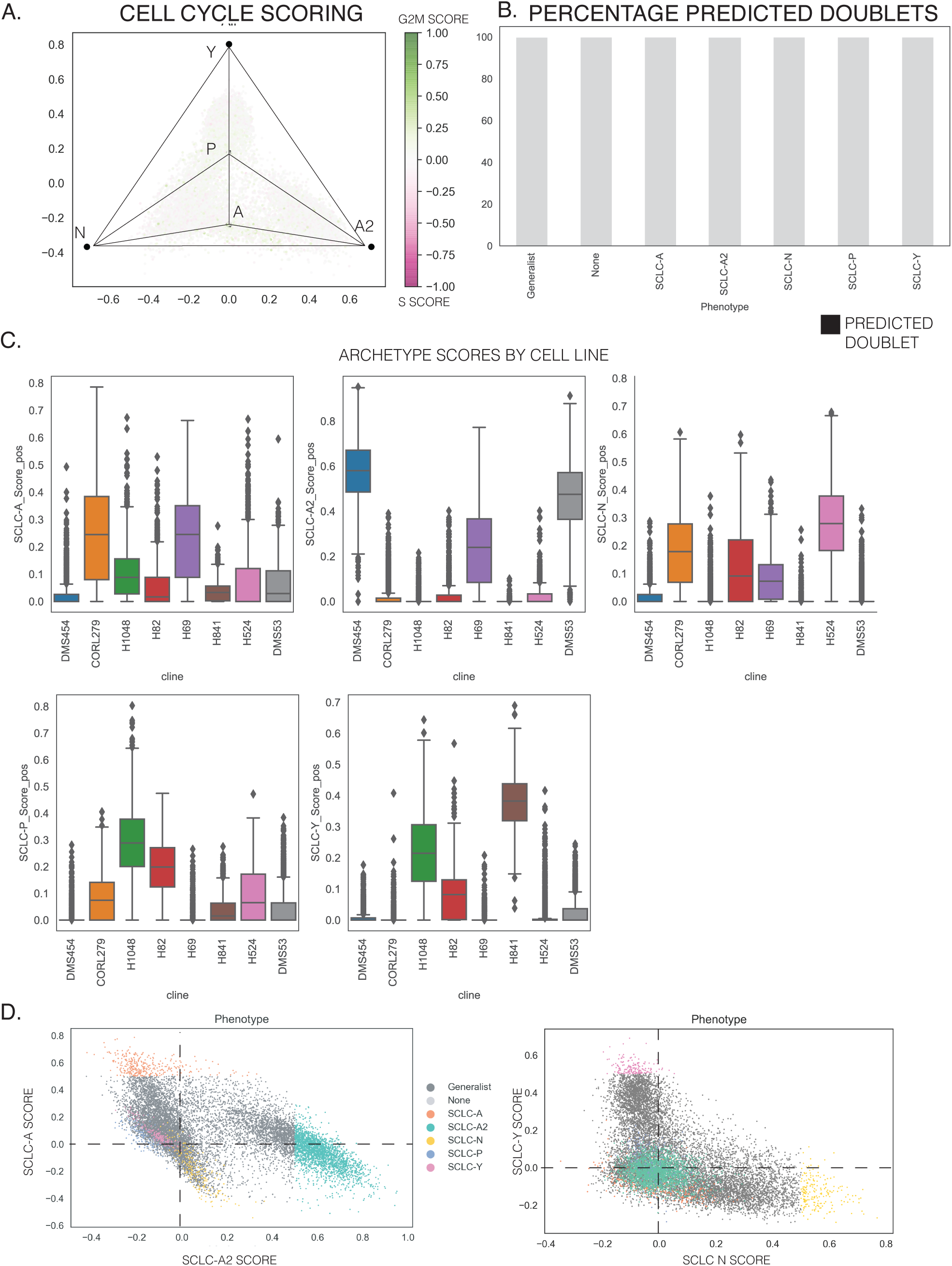
Single Cell RNA-seq on Cell Lines A. G2M and S-associated genes are scored based on average expression compared to average expression of a reference set. Scores are visualized using Locally Linear Embedding (LLE) and show that location of cells within the archetype plot is not dependent on cell cycle, as most cells show very low average expression of G2M or S phase-related genes. B. Predicted doublets using Scrublet are shown as bar plots for each phenotype (Detailed in C and Methods). Very few cells are predicted to be doublets, which are virtually unnoticeable in the bar plot. Critically, generalists are not significantly more likely to be doublets as shown in the table, which would have an intermediate expression profile and confound results. Black = predicted doublets, grey = not predicted to be a doublet. C. Boxplots of archetype scores using least squares approximation described in Methods. Scores generally range between 0 (no similarity) and 1 (cell is at archetype location in reduced gene expression space). Scores for each of the five archetypes range in values across cell lines, with higher archetype score distributions generally corresponding to the expected subtype according to hierarchical clustering (**Supplemental Figure 2A**). A few cell lines, such as H82, show high levels for multiple archetype scores (e.g., N, P, and Y). D. Two example scatterplots of archetype scores show classification of cells into specialists, close to a specific archetype, or generalists, which have scores < 0.5 for all five archetypes. (Left) SCLC-A vs. SCLC-A2 scores show two distinct populations of cells, one along the Y axis (SCLC-A) and the other showing a negative correlation between SCLC-A and SCLC-A2 scores. (Right) SCLC-N vs SCLC-Y scores show a trade-off between the two archetypes, with very few cells high in both scores.

**Supplemental Figure 5:**
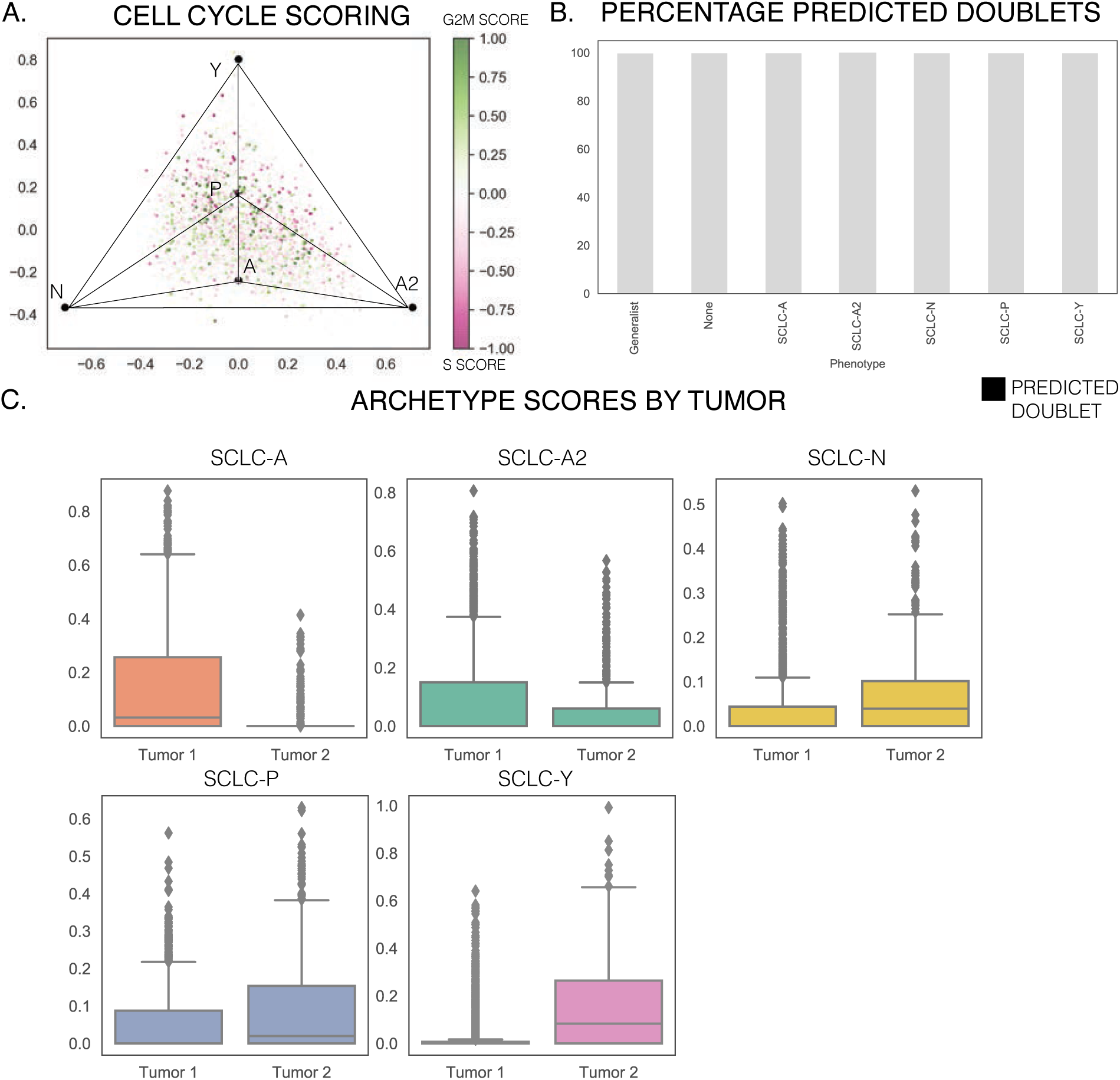
Single Cell Preprocessing on Human Tumors A. Cell cycle scoring using G2M and S phase-associated gene lists. While some cells show higher average expression for G2M or S phase genes compared to human cell lines in **Supplemental Figure 4A**, most cells show low scores. B. Predicted doublets using Scrublet. Similar to human cell lines, very few cells are predicted to be doublets. C. Archetype scores for each tumor. As expected from **Figure 4A**, Tumor 1 has higher scores for SCLC-A and SCLC-A2, while Tumor 2 shows higher scores for SCLC-N, -P, and -Y. Tumor 2 has an LCNEC component, which may contribute to its higher SCLC-Y score, as several studies have shown that SCLC-Y is most similar to non-SCLC lung cancer with lower neuroendocrine scores.^28, 31^

**Supplemental Figure 6:**
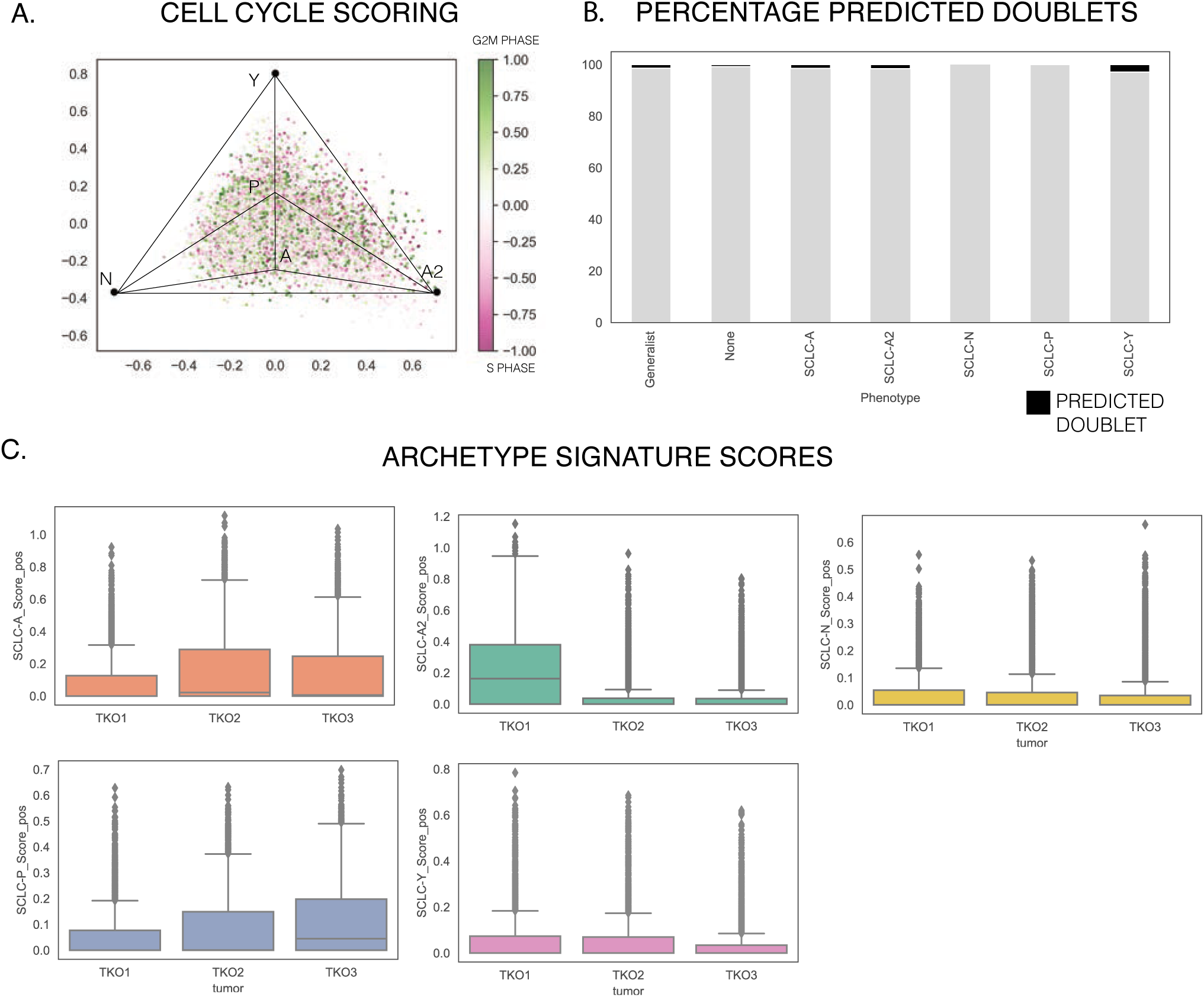
Single Cell Processing on TKO Mouse Tumors A. Cell cycle gene scores for S and G2M-associated genes. Location in archetype space shows no correlation to cell cycle position, which range from S and G2M (cycling) to low scores for both gene sets (not actively cycling). B. Predicted doublets using Scrublet by phenotype. Very few cells are predicted to be doublets, shown in black. Critically, generalists are not significantly more likely to be doublets, which would have an intermediate expression profile and confound results. C. Archetype scores for each timepoint. The score for the SCLC-A2 archetype is higher in TKO1, while SCLC-A and SCLC-P scores are higher in TKO2 and TKO3. SCLC-N and SCLC-Y are low, on average, across all tumors.

**Supplemental Figure 7:**
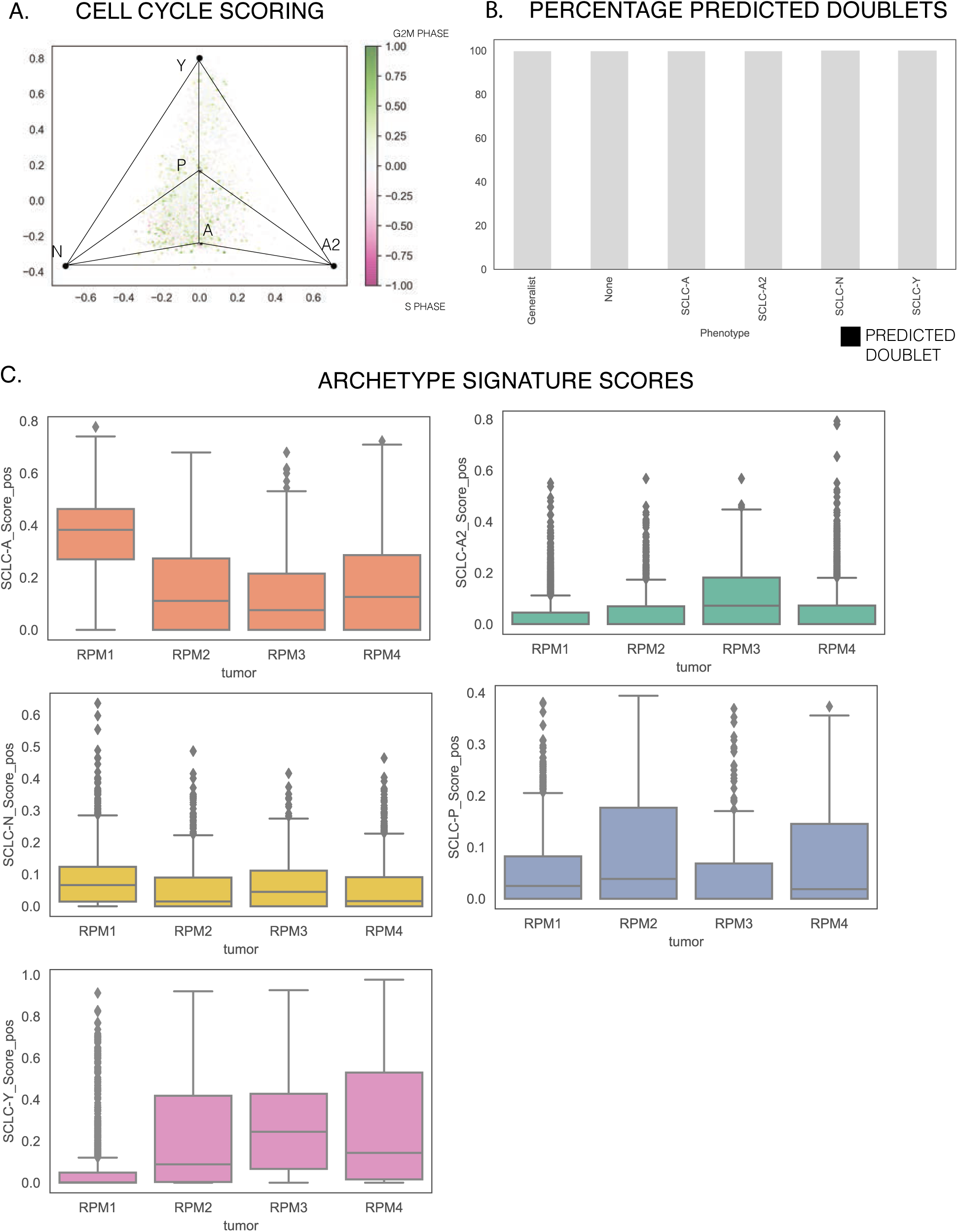
Single Cell Preprocessing on RPM Mouse Tumors A. Cell cycle gene scores for S and G2M-associated genes. A majority of cells are not actively cycling, with low scores for both S and G2M genes. B. Predicted doublets using Scrublet. Similar to human cell lines and tumors, very few cells are predicted to be doublets. C. Archetype signature scores for each tumor. RPM1 has much higher SCLC-A scores, while SCLC-A2 scores are slightly higher in RPM3. SCLC-P is enriched in RPM2 and RPM4, and SCLC-Y is much higher in RPM 2-4 than RPM1. This suggests RPM1 is more neuroendocrine, and RPM 2-4 are more non-neuroendocrine.

**Supplemental Figure 8:**
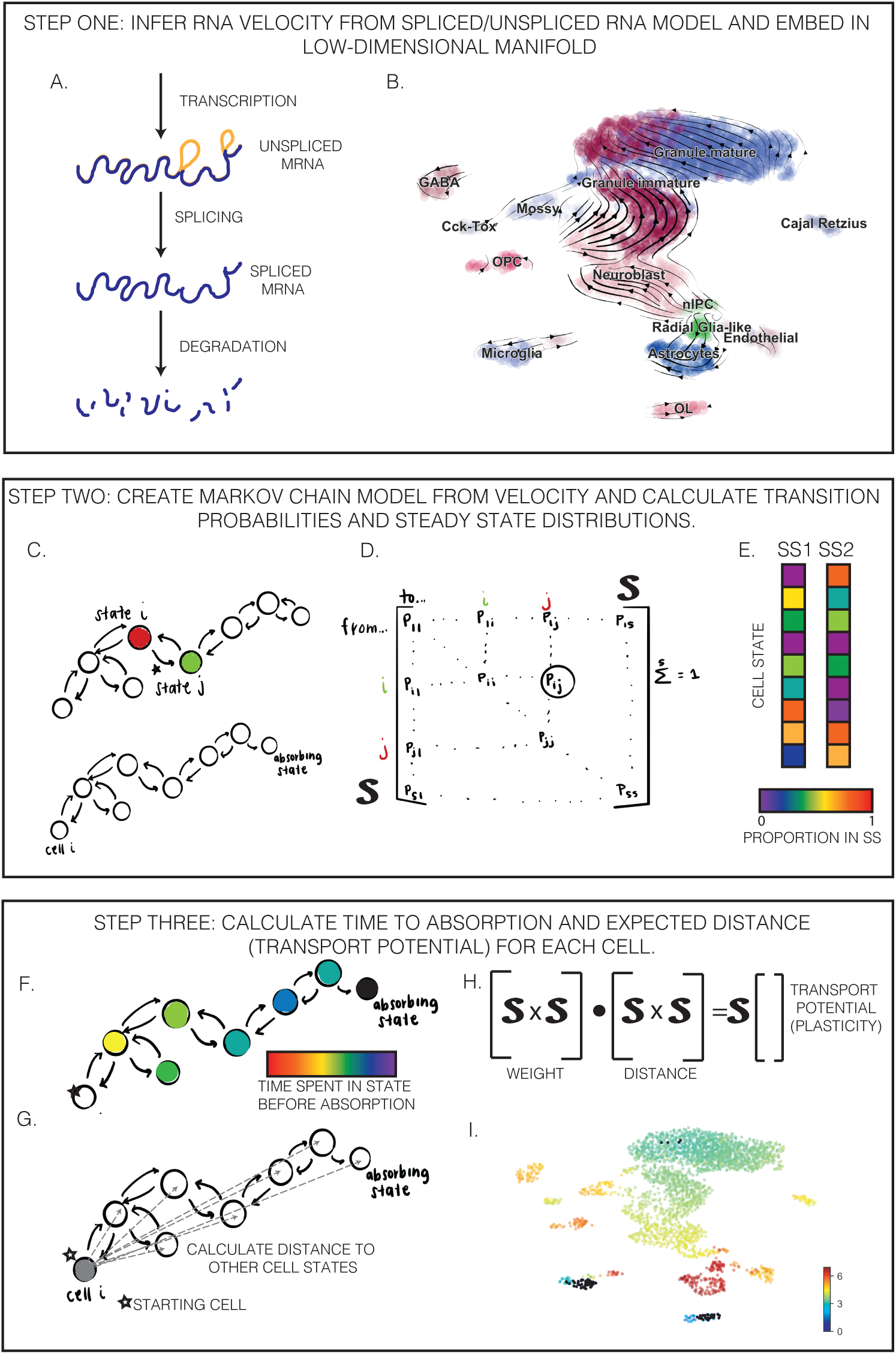
Plasticity Pipeline. Steps for calculating Cell Transport Potential from RNA velocity on single cell data. - Step 1. A. RNA splicing model allows for prediction of future transcriptomic state for individual cells. B. Example of RNA velocity overlaid on neuronal differentiation using scVelo package (See Methods and **Supplemental Figure 9** for more details). - Step 2: C. A Markov Chain model (MC model) is generated in which each sampled cell is its own state and can transition to any other state. Velocity-inferred MC models are generally absorbing chains in which end states (which transition to other states with negligible probability) constitute a steady state of the system. D. Example transition matrix for an MC model. E. Mock steady states of the system, showing the distribution across states in the chain. - Step Three: F. Fundamental matrix of MC is calculated using the inverse of the transition matrix in E. For each state, this matrix defines the time spent in other states before absorption. G. Distances between all states are calculated on low-dimensional manifold (UMAP space) or linear PCA space. H. Transport potential is calculated as the dot product of proportion of time in each state (normalized to sum to 1 for each cell) and distance. I. Plasticity can be visualized on UMAP, as shown here, or other dimensionality reduction embeddings (such as LLE).

**Supplemental Figure 9:**
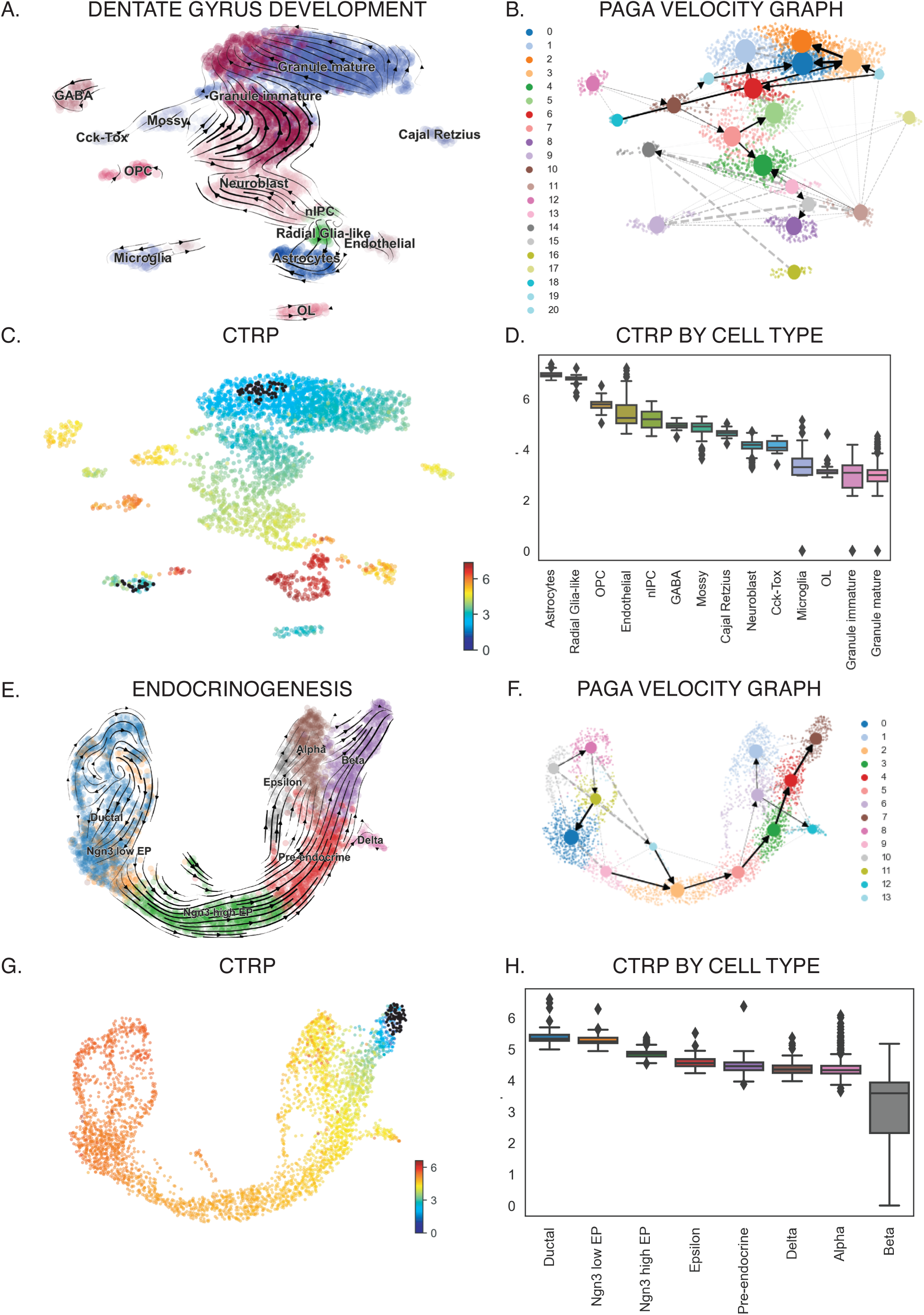
Test cases with known ground truth for CTrP pipeline A. Dentate Gyrus development UMAP with RNA velocity streams overlaid by scVelo. Significant, directed movement can be seen starting by progenitor cell types (Radial Glia-like cells and nIPCs). Differentiation of these cells results in several easily distinguishable clusters representing different cell types. Cell type cluster labels are from scVelo dataset, originally described in Hochgerner et al. (2018).^81, 138^ B. PAGA velocity graph for Dentate Gyrus development. Cells are clustered by the Leiden clustering method, shown by color on plot. Dotted arrows signify clusters with high connectivity, and arrows show significant transitions between clusters. Of note, pink cluster 13 overlaps with progenitor cells and shows transitions to neighboring cell types. C. CTrP analysis shown by color scale, with black dots signifying absorbing end states for Markov Chain model. High plasticity cell types correspond to progenitor cells and immature astrocytes, while low plasticity end states fall in terminally differentiated regions of the UMAP. D. CTrP by cell type, ordered by median value. Progenitor cells show highest CTrP on the left side of the plot, with more differentiated cell types showing decreasing plasticity towards the right side of the plot. E. Endocrinogenesis process showing differentiation of ductal cells into pancreatic beta cells from Bastidas-Ponce et al.^139^RNA velocity streams overlaid on UMAP show trajectory of differentiation. F. PAGA velocity graph shows movement from early progenitors on the left to differentiated pancreatic cells on the right. G. CTrP analysis shown by color scale, with black dots signifying absorbing end states. High plasticity cells fall in the Ductal cell cluster, with decreasing plasticity along the trajectory to beta cells. H. CTrP by cell type, ordered by median value. End states with low plasticity are only found in the beta cell cluster.

**Supplemental Figure 10:**
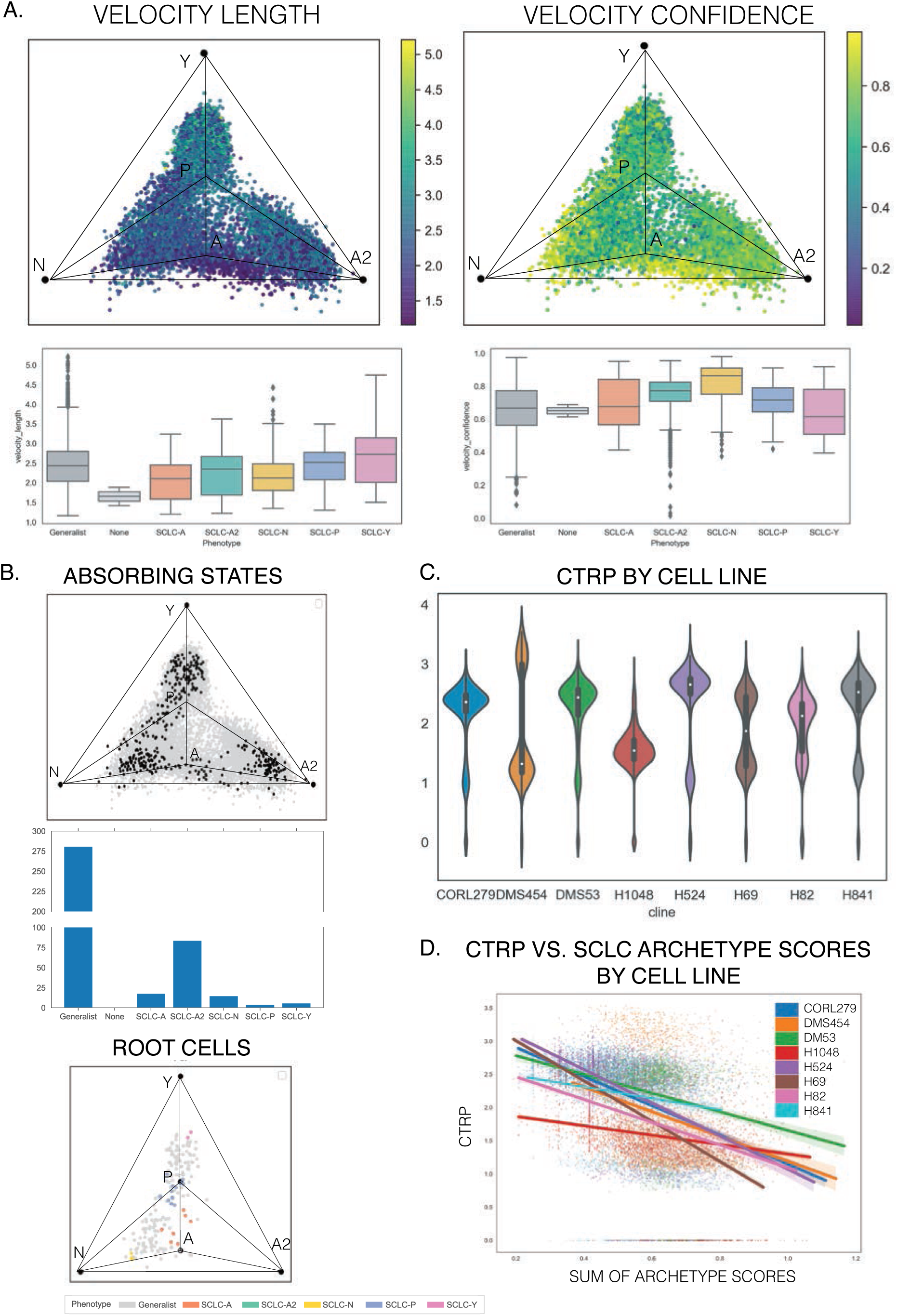
Plasticity analysis for SCLC human cell lines A. (Left) Velocity length shows the magnitude of velocity in gene expression space for each cell. When compared to **Supplemental Figures 11-13**, we can see that these cells have lower velocities, possibly due to their consistent *in vitro* environments. (Right) Velocity confidence shows coherence of velocities within a defined neighborhood for each cell. Higher confidence for a cell suggests that velocities for cells within a neighborhood are similar in direction to the cell. A majority of cells have confidence levels above 0.6. B. (Top) Absorbing states, as defined in Methods and **Supplemental Figure 8B**. Absorbing cells (end points) are shown as small black circles, while transition cells are gray. Large black circles provide archetype reference locations in LLE. (Middle) Distribution of absorbing states by phenotype. Most end states are generalists, reflective of the fact that a large proportion of cells in the cell lines are generalists. Every archetype specialist has associated absorbing states. (Bottom) Root cells from Markov Chain analysis show starting states by phenotype. Each phenotype except SCLC-A2 cells have root cells in human cell lines. C. CTrP by cell line. Most cell lines have two distinct subpopulations of high and low plasticity, and every cell line has end states with plasticity equal to 0. D. Scatterplots of CTrP values versus sum of archetype scores by cell line are shown with lines of best fit. Five archetype scores (one for each vertex: A, A2, N, P, and Y) are summed to give the overall similarity of each cell to SCLC archetype signatures. A negative trend was found for plasticity vs. archetype scores in every cell line, suggesting that cells less similar to SCLC archetypes (generalists and unclassified “None” cells) have higher plasticity.

**Supplemental Figure 11:**
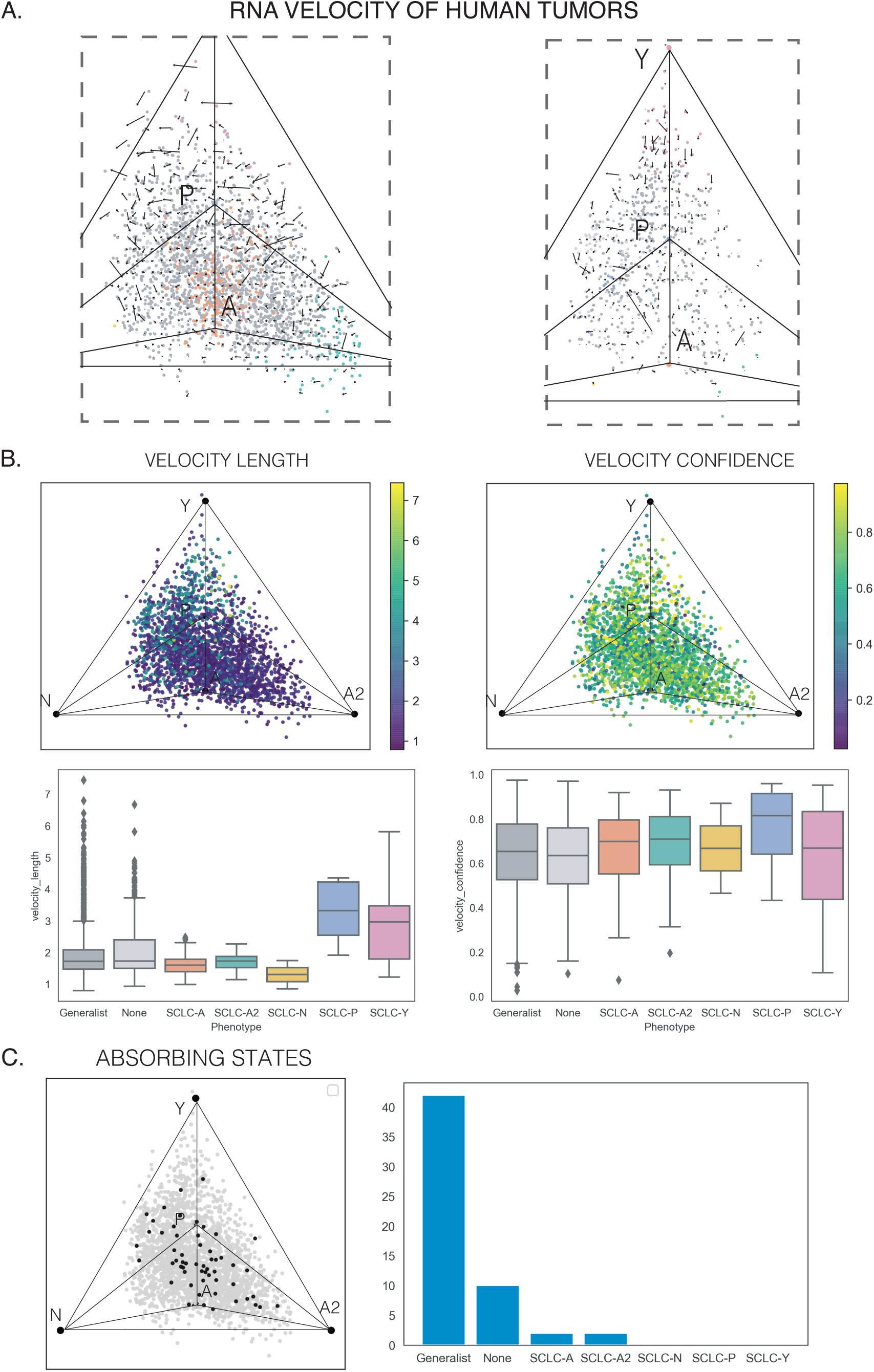
Plasticity analysis for SCLC human tumors A. RNA velocity fields for human tumors. Neither tumor shows coherent directionality across the archetype space. B. (Left) Velocity length showing the magnitude of velocity for each cell shows slight dependence on location in archetype space. SCLC-P and SCLC-Y cells have higher velocity than other cell types, shown in the boxplot below the LLE plot. (Right) Velocity confidence shows little dependence on archetype space location, with a majority of cells having confidence above 0.6. C. Absorbing states shown in the LLE space. End states, shown as small black dots, are found across the archetype space between SCLC-A and SCLC-A2 archetypes. A large proportion of absorbing states are generalists.

**Supplemental Figure 12:**
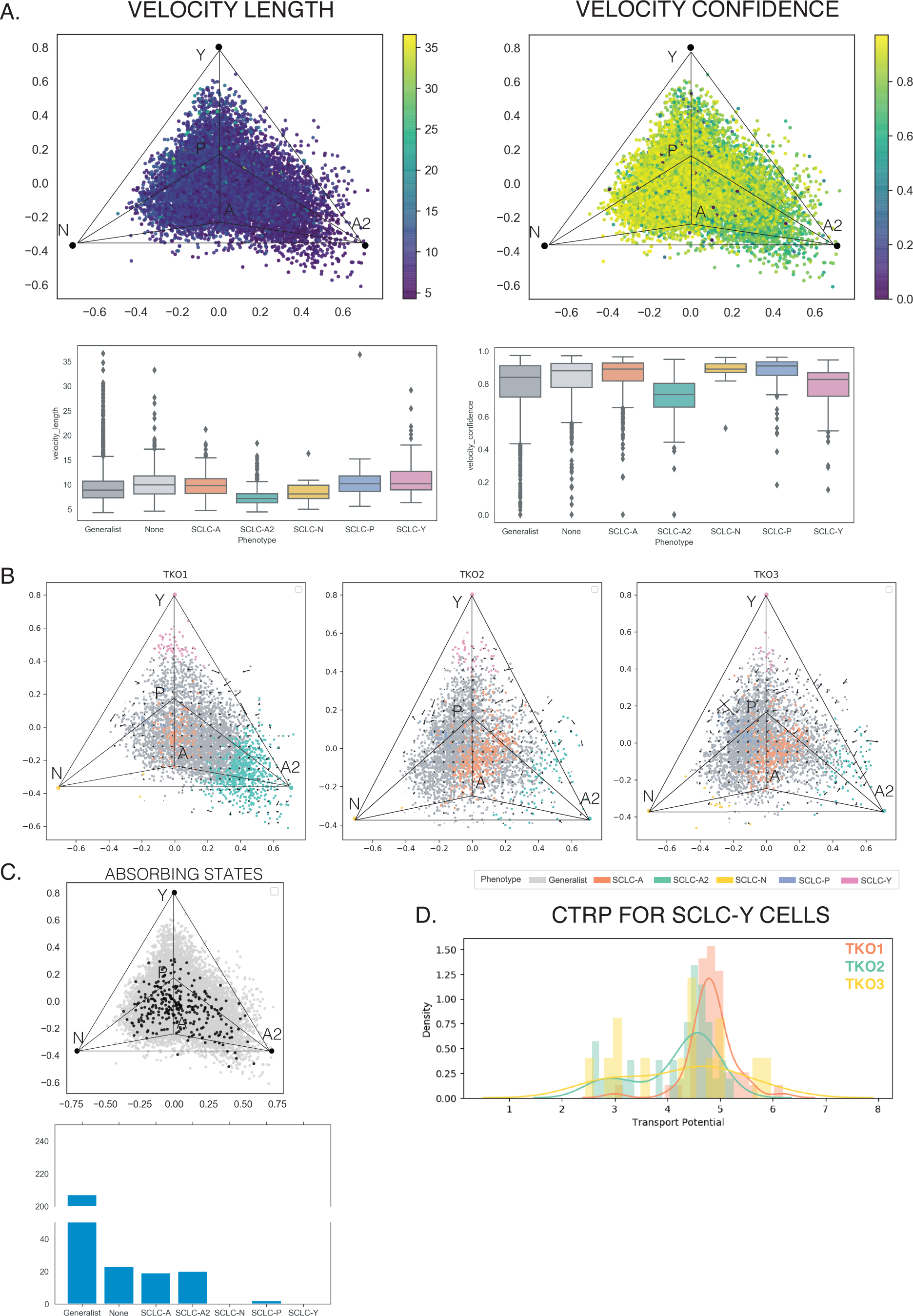
Plasticity analysis for TKO mouse tumors A. (Left) Velocity length shows little dependence on location in archetype space. Similarly, no major patterns emerge between velocity length for each phenotype in the boxplot below the LLE plot. (Right) Velocity confidence shows little dependence on archetype space location, with a majority of cells having confidence above 0.8. B. RNA velocity fields for TKO tumors. None of the tumors shows coherent directionality across the archetype space, with overall low velocity across the archetype space. C. Absorbing states shown in the LLE space. End states, shown as small black dots, are found across the archetype space. A large proportion of absorbing states are generalists and unclassified cells, and SCLC-A, -A2, and -P are all represented. D. CTrP for SCLC-Y cells across tumors. TKO1 has slightly more plastic SCLC-Y cells than the other two tumors. Different distributions between TKO2 and TKO3 demonstrate that, even within the same mouse, each phenotype (i.e., SCLC-Y) is capable of different levels of plasticity depending on unknown factors. We predict that these differences are due to differences in environmental conditions between a primary and metastatic tumor that affect the transcription factor network regulating cell identity.

**Supplemental Figure 13:**
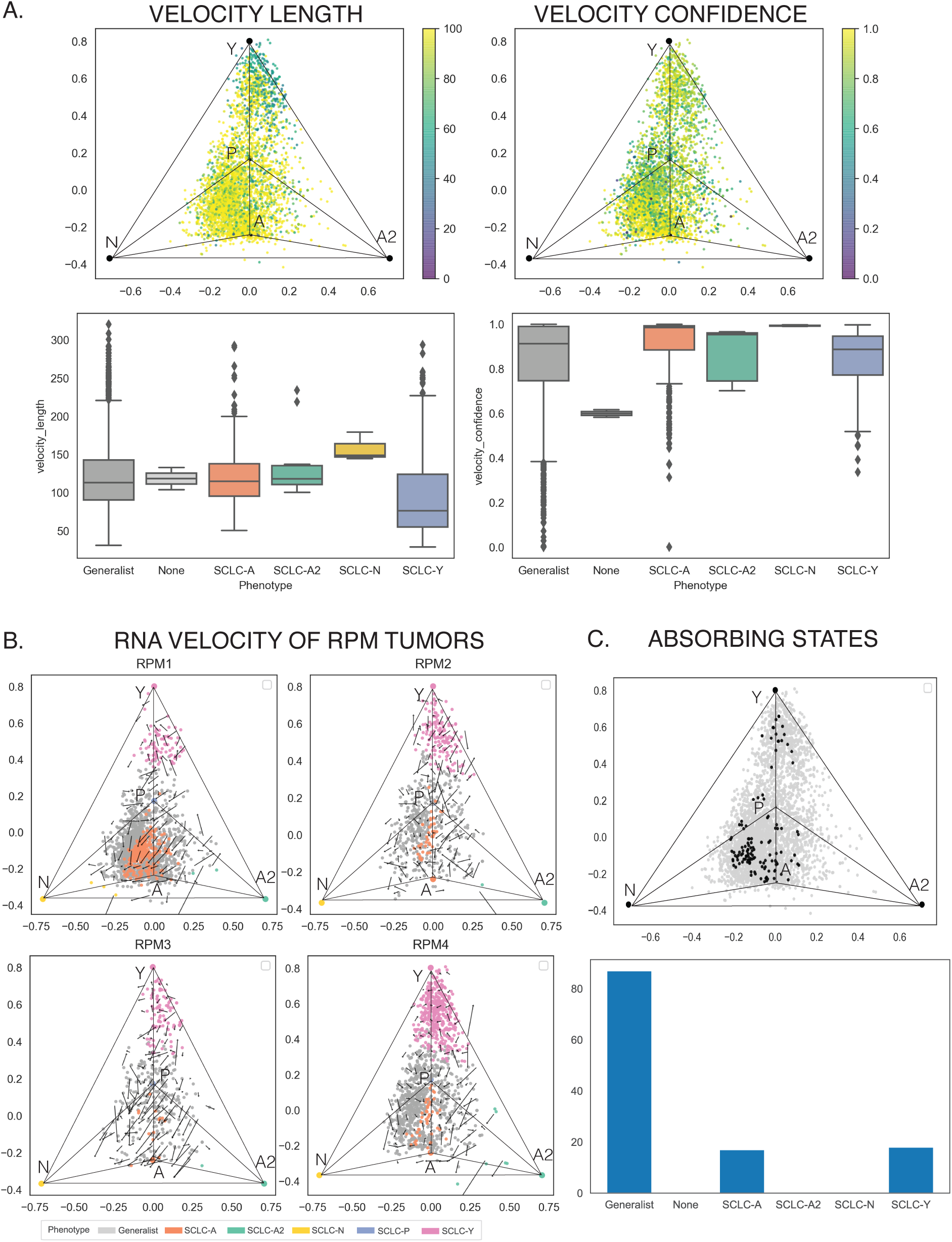
Plasticity analysis for RPM mouse tumors A. (Left) Velocity length showing the magnitude of velocity for each cell shows slight dependence on location in archetype space. SCLC-Y cells have slightly lower velocity than other cell types, shown in the boxplot below the LLE plot. Overall, cells in these tumors have high velocity on average compared to other samples. (Right) Velocity confidence shows little dependence on archetype space location, with a majority of cells having confidence above 0.8. B. RNA velocity fields for human tumors. The tumors show little consistent directionality in the RNA velocity field through archetype space. Some cells show large velocities towards the SCLC-N archetype, though it is unclear when in the pseudotime trajectory introduced by Ireland et al.^47^ these cells are transitioning. C. Absorbing states shown in the LLE space. End states, shown as small black dots, are found across the archetype space between SCLC-A and SCLC-Y archetypes. Again, a large proportion of absorbing states are generalists.

**Supplemental Figure 14:**
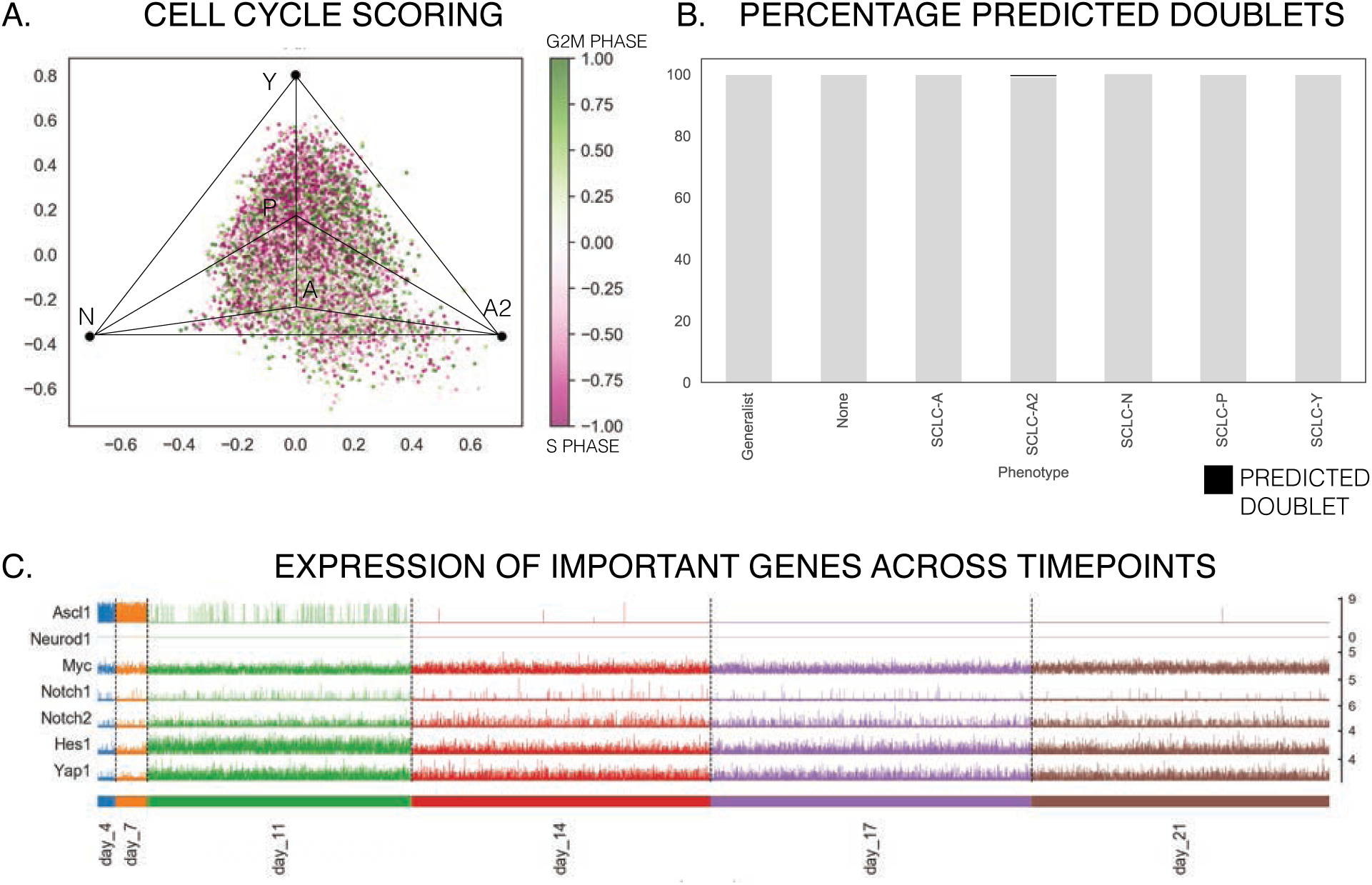
Single cell subtyping for RPM time series data A. Cell cycle scoring using G2M and S phase-associated gene lists. Most cells show high scores for G2M or S phase-associated genes, suggesting the cells are actively cycling. B. Predicted doublets using Scrublet. Very few cells are predicted to be doublets. C. Expression of important genes across timepoints. Genes expected to change across timepoints from Ireland et al. ASCL1 clearly decreases over the time course, as YAP1 and NOTCH pathway genes increase by day 11. MYC expression is relatively constant throughout, consistent with its stable overexpression in the RPM mouse model.

**Supplemental Figure 15:**
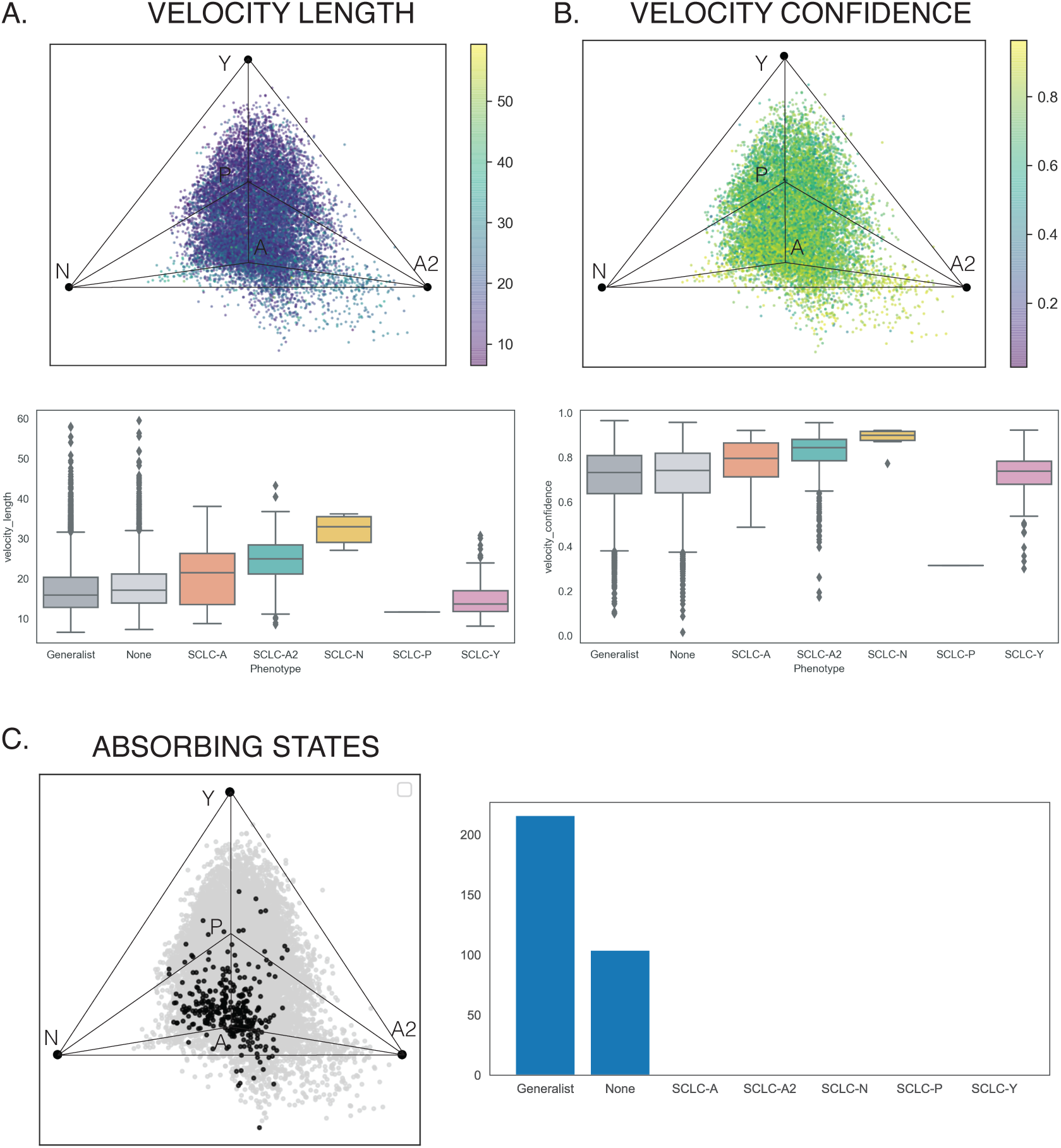
Plasticity analysis on RPM time series data A. Velocity length shows highest velocity cells between SCLC-A, -A2, and -N, consistent with their quickly changing gene expression profiles early on in the time course. B. Velocity confidence, showing coherence of velocity directions, is high across cells and not dependent on location in archetype space. C. Absorbing states, or end points in the Markov Chain model, are shown as small black circles and mostly consist of generalist and unclassified cells.

**Supplemental Figure 16:**
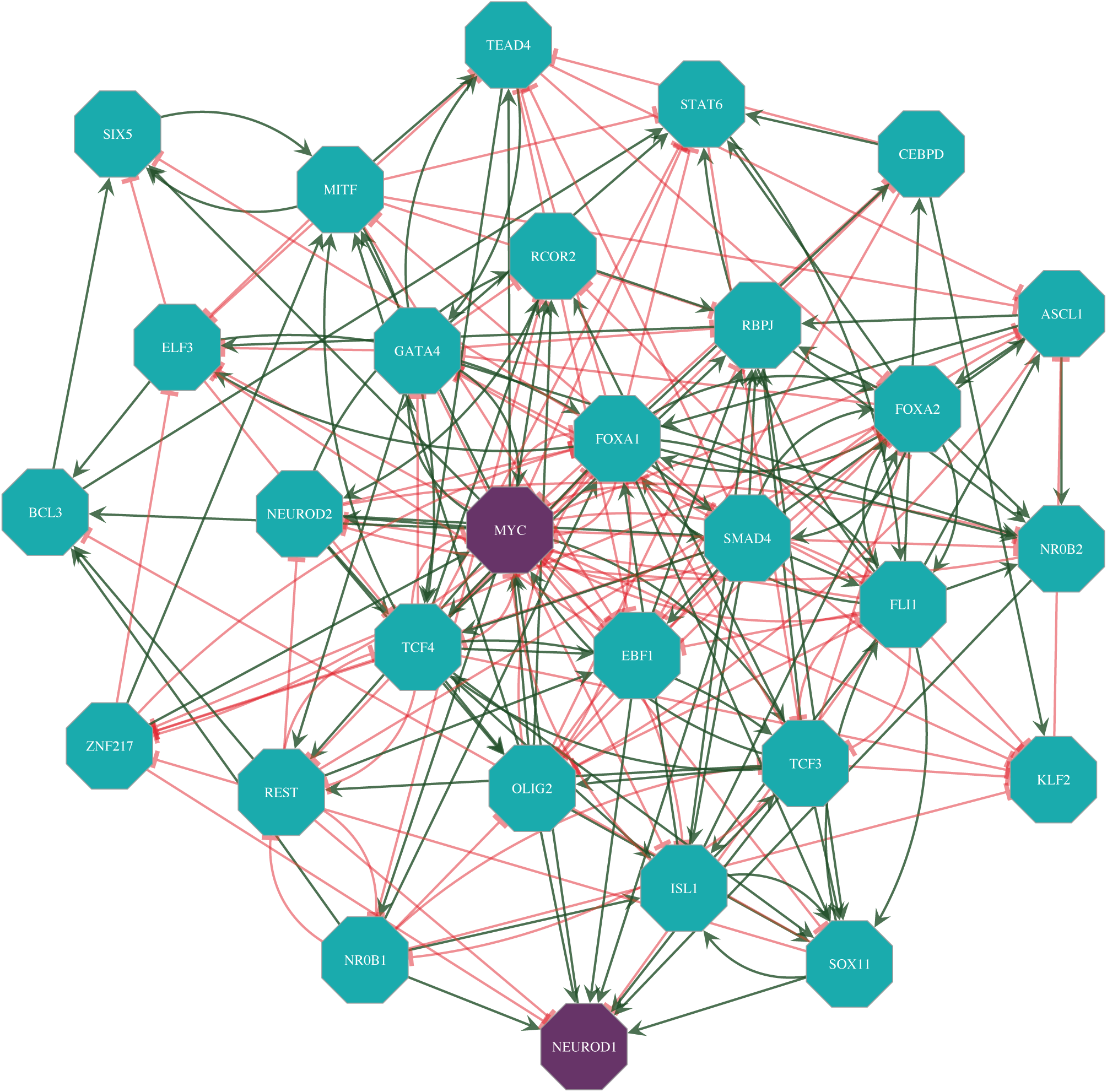
SCLC transcription factor network with MYC and NEUROD1 A. Network adopted from Wooten et al.^28^ to include MYC (and subsequently NEUROD1). Red arrow: inhibiting; Green arrow: activating.

## Supplemental Tables

Supplemental Table 1: Enriched subtype labels (from hierarchical clustering) at each archetype location.

Supplemental Table 2: Table of q values for gene set/task enrichment at archetypes for bulk RNA-seq data. Q <.1 is considered significant; Mann-Whitney test.

Supplemental Table 3: Table of q values for Hallmark Gene Set enrichment at archetypes for bulk RNA-seq data. Q <.1 is considered significant; Mann-Whitney test.

Supplemental Table 4: Table of q values for cancer hallmark gene set enrichment at archetypes for bulk RNA-seq data. Q <.1 is considered significant; Mann-Whitney test.

Supplemental Table 5: Table of q values for gene enrichment at archetypes for bulk RNA-seq data. Q <.1 is considered significant; Mann-Whitney test.

Supplemental Table 6: Archetype gene signature expression data.

## References

1. Sutherland et al. *Cell* of Origin of Small Cell Lung Cancer: Inactivation of Trp53 and Rb1 in Distinct Cell Types of Adult Mouse Lung. Cancer Cell 19, 754–764 (2011).

2. Park, K.-S. et al. Characterization of the cell of origin for small cell lung cancer. Cell Cycle 10, 2806– 2815 (2014).

3. Song, H. et al. Functional characterization of pulmonary neuroendocrine cells in lung development, injury, and tumorigenesis. Proc National Acad Sci 109, 17531–17536 (2012).

4. Park, J. W. et al. Reprogramming normal human epithelial tissues to a common, lethal neuroendocrine cancer lineage. Science 362, 91–95 (2018).

5. Huang, Y.-H. et al. POU2F3 is a master regulator of a tuft cell-like variant of small cell lung cancer. Gene Dev 32, 915–928 (2018).

6. Semenova, E. A., Nagel, R. & Berns, A. Origins, genetic landscape, and emerging therapies of small cell lung cancer. Gene Dev 29, 1447–62 (2015).

7. Gazdar, A. F., Bunn, P. A. & Minna, J. D. Small-cell lung cancer: what we know, what we need to know and the path forward. Nat Rev Cancer 17, 725 (2017).

8. George, J. et al. Comprehensive genomic profiles of small cell lung cancer. Nature 524, 47 (2015).

9. Borromeo, M. D. et al. ASCL1 and NEUROD1 Reveal Heterogeneity in Pulmonary Neuroendocrine Tumors and Regulate Distinct Genetic Programs. Cell Reports 16, 1259–1272 (2016).

10. Gazdar, A. F., Carney, D. N., Nau, M. M. & Minna, J. D. Characterization of variant subclasses of cell lines derived from small cell lung cancer having distinctive biochemical, morphological, and growth properties. vol. 45 (1985).

11. Rudin, C. M. et al. Molecular subtypes of small cell lung cancer: a synthesis of human and mouse model data. Nat Rev Cancer 19, 289–297 (2019).

12. Mollaoglu, G. et al. MYC Drives Progression of Small Cell Lung Cancer to a Variant Neuroendocrine Subtype with Vulnerability to Aurora Kinase Inhibition. Cancer Cell 31, 270–285 (2017).

13. Horie, M., Saito, A., Ohshima, M., Suzuki, H. I. & Nagase, T. YAP and TAZ modulate cell phenotype in a subset of small cell lung cancer. Cancer Sci 107, 1755–1766 (2016).

14. Altschuler, S. J. & Wu, L. F. Cellular Heterogeneity: Do Differences Make a Difference? vol. 141 (2010).

15. Brady, S. W. et al. Combating subclonal evolution of resistant cancer phenotypes. Nat Commun 8, 1231 (2017).

16. Marusyk, A., Almendro, V. & Polyak, K. Intra-tumour heterogeneity: a looking glass for cancer? vol. 12 (2012).

17. Pisco, A. O. & Huang, S. Non-genetic cancer cell plasticity and therapy-induced stemness in tumour relapse: ‘What does not kill me strengthens me.’ Brit J Cancer 112, 1725–1732 (2015).

18. Lawrence, M. S. et al. Mutational heterogeneity in cancer and the search for new cancer-associated genes. Nature 499, 214 (2013).

19. Tirosh, I. et al. Dissecting the multicellular ecosystem of metastatic melanoma by single-cell RNA-seq. Science 352, 189–196 (2016).

20. Sáez-Ayala, M. et al. Directed Phenotype Switching as an Effective Antimelanoma Strategy. Cancer Cell 24, 105–119 (2013).

21. Arozarena, I. & Wellbrock, C. Phenotype plasticity as enabler of melanoma progression and therapy resistance. Nat Rev Cancer 1–15 (2019) doi:10.1038/s41568-019-0154-4.

22. Su, Y. et al. Phenotypic heterogeneity and evolution of melanoma cells associated with targeted therapy resistance. Plos Comput Biol 15, e1007034 (2019).

23. Howard, G. R., Johnson, K. E., Ayala, A. R., Yankeelov, T. E. & Brock, A. A multi-state model of chemoresistance to characterize phenotypic dynamics in breast cancer. Sci Rep-uk 8, 12058 (2018).

24. Gupta, P. B. et al. Stochastic State Transitions Give Rise to Phenotypic Equilibrium in Populations of Cancer Cells. Cell 146, 633–644 (2011).

25. Jia, D., Jolly, M. K., Kulkarni, P. & Levine, H. Phenotypic Plasticity and Cell Fate Decisions in Cancer: Insights from Dynamical Systems Theory. Cancers 9, 70 (2017).

26. Lim, J. S. et al. Intratumoural heterogeneity generated by Notch signalling promotes small-cell lung cancer. Nature 545, 360 (2017).

27. Udyavar, A. R. et al. Novel Hybrid Phenotype Revealed in Small Cell Lung Cancer by a Transcription Factor Network Model That Can Explain Tumor Heterogeneity. Cancer Res 77, 1063–1074 (2017).

28. Wooten, D. J. et al. Systems-level network modeling of Small Cell Lung Cancer subtypes identifies master regulators and destabilizers. Plos Comput Biol 15, e1007343 (2019).

29. Newman, A. M. et al. Robust enumeration of cell subsets from tissue expression profiles. Nat Methods 12, nmeth.3337 (2015).

30. Stewart, C. A. et al. Single-cell analyses reveal increased intratumoral heterogeneity after the onset of therapy resistance in small-cell lung cancer. Nature Cancer (2020) doi:10.1038/s43018-019-0020-z.

31. Zhang, W. et al. Small cell lung cancer tumors and preclinical models display heterogeneity of neuroendocrine phenotypes. Transl Lung Cancer Res 7, 32–49 (2018).

32. McColl, K. et al. Reciprocal expression of INSM1 and YAP1 defines subgroups in small cell lung cancer. Oncotarget 5, 73745–73756 (2015).

33. Poirier, J. et al. DNA methylation in small cell lung cancer defines distinct disease subtypes and correlates with high expression of EZH2. Oncogene 34, 5869–5878 (2015).

34. Krushkal, J. et al. Epigenome-wide DNA methylation analysis of small cell lung cancer cell lines suggests potential chemotherapy targets. Clin Epigenetics 12, 93 (2020).

35. Kalari, S., Jung, M., Kernstine, K. H., Takahashi, T. & Pfeifer, G. P. The DNA methylation landscape of small cell lung cancer suggests a differentiation defect of neuroendocrine cells. Oncogene 32, 3559 (2013).

36. Krohn, A. et al. Tumor Cell Heterogeneity in Small Cell Lung Cancer (SCLC): Phenotypical and Functional Differences Associated with Epithelial-Mesenchymal Transition (EMT) and DNA Methylation Changes. Plos One 9, e100249 (2014).

37. Mørup, M. & Hansen, L. K. Archetypal analysis for machine learning and data mining. Neurocomputing 80, 54–63 (2012).

38. Shoval, O., et al. Evolutionary Trade-Offs, Pareto Optimality, and the Geometry of Phenotype Space. Science 336, 1157–1160 (2012).

39. Aubert, O. et al. Archetype Analysis Identifies Distinct Profiles in Renal Transplant Recipients with Transplant Glomerulopathy Associated with Allograft Survival. Journal of the American Society of Nephrology 30, (2019).

40. Adler, M., Kohanim, Y. K., Tendler, A., Mayo, A. & Alon, U. Continuum of Gene-Expression Profiles Provides Spatial Division of Labor within a Differentiated Cell Type. Cell Syst 8, 43–52.e5 (2019).

41. Tsai, J. H. & Yang, J. Epithelial–mesenchymal plasticity in carcinoma metastasis. Gene Dev 27, 2192–2206 (2013).

42. Rueffler, C., Hermisson, J. & Wagner, G. P. Evolution of functional specialization and division of labor. Proc National Acad Sci 109, E326–E335 (2012).

43. Sheftel, H., Szekely, P., Mayo, A., Sella, G. & Alon, U. Evolutionary trade-offs and the structure of polymorphisms. Philosophical Transactions Royal Soc B 373, 20170105 (2018).

44. Korem, Y. et al. Geometry of the Gene Expression Space of Individual Cells. Plos Comput Biol 11, e1004224 (2015).

45. Hausser, J. et al. Tumor diversity and the trade-off between universal cancer tasks. Nature Communications 10, 5423 (2019).

46. Cutler & Breiman. Archetypal Analysis. (1994).

47. Ireland, A. S. et al. MYC Drives Temporal Evolution of Small Cell Lung Cancer Subtypes by Reprogramming Neuroendocrine Fate. Cancer Cell (2020) doi:10.1016/j.ccell.2020.05.001.

48. Hatzikirou, H., Basanta, D., Simon, M., Schaller, K. & Deutsch, A. “Go or Grow”: the key to the emergence of invasion in tumour progression? Math Medicine Biology 29, 49–65 (2010).

49. Gallaher, J. A., Brown, J. S. & Anderson, A. R. A. The impact of proliferation-migration tradeoffs on phenotypic evolution in cancer. Sci Rep-uk 9, 2425 (2019).

50. Hanahan, D. & Weinberg, R. A. Hallmarks of Cancer: The Next Generation. Cell 144, 646–674 (2011).

51. Hanahan, D. & Weinberg, R. A. The Hallmarks of Cancer. Cell 100, 57–70 (2000).

52. Calbo, J. et al. A Functional Role for Tumor Cell Heterogeneity in a Mouse Model of Small Cell Lung Cancer. Cancer Cell 19, 244–256 (2011).

53. Carney, D. N. et al. Establishment and identification of small cell lung cancer cell lines having classic and variant features. Cancer Res 45, 2913–23 (1985).

54. Bepler, G., Jaques, G., Koehler, A., Gropp, C. & Havemann, K. Markers and characteristics of human SCLC cell lines. J Cancer Res Clin 113, 253–259 (1987).

55. Hart, Y. et al. Inferring biological tasks using Pareto analysis of high-dimensional data. Nat Methods 12, 233–235 (2015).

56. Liberzon, A. et al. Molecular signatures database (MSigDB) 3.0. Bioinformatics 27, 1739–1740 (2011).

57. Liberzon, A. et al. The Molecular Signatures Database Hallmark Gene Set Collection. Cell Syst 1, 417–425 (2015).

58. Subramanian, A. et al. Gene set enrichment analysis: A knowledge-based approach for interpreting genome-wide expression profiles. P Natl Acad Sci Usa 102, 15545–15550 (2005).

59. Zhang, D. et al. CHG: A Systematically Integrated Database of Cancer Hallmark Genes. Frontiers Genetics 11, 29 (2020).

60. Sen, T., Gay, C. M. & Byers, L. A. Targeting DNA damage repair in small cell lung cancer and the biomarker landscape. Transl Lung Cancer Res 7, 50–68 (2018).

61. Alam, Sk. K., et al. ASCL1-regulated DARPP-32 and t-DARPP stimulate small cell lung cancer growth and neuroendocrine tumour cell proliferation. Brit J Cancer 123, 819–832 (2020).

62. Kuo, C. S. & Krasnow, M. A. Formation of a Neurosensory Organ by Epithelial Cell Slithering. Cell 163, 394–405 (2015).

63. Li, N. & Li, S. Epigenetic inactivation of SOX1 promotes cell migration in lung cancer. Tumor Biol 36, 4603–4610 (2015).

64. Garcia, I. et al. Oncogenic activity of SOX1 in glioblastoma. Sci Rep-uk 7, 46575 (2017).

65. Venere, M. et al. Sox1 marks an activated neural stem/progenitor cell in the hippocampus. Development 139, 3938–3949 (2012).

66. Elkouris, M. et al. Sox1 Maintains the Undifferentiated State of Cortical Neural Progenitor Cells via the Suppression of Prox1-Mediated Cell Cycle Exit and Neurogenesis. Stem Cells 29, 89–98 (2011).

67. Brägelmann, J. et al. Family matters: how MYC family oncogenes impact small cell lung cancer. Cell cycle (Georgetown, Tex.) 16, 0 (2017).

68. Kuleshov, M. V. et al. Enrichr: a comprehensive gene set enrichment analysis web server 2016 update. Nucleic Acids Res 44, W90–W97 (2016).

69. Chen, E. Y. et al. Enrichr: interactive and collaborative HTML5 gene list enrichment analysis tool. Bmc Bioinformatics 14, 128 (2013).

70. Zhang, S., Li, M., Ji, H. & Fang, Z. Landscape of transcriptional deregulation in lung cancer. Bmc Genomics 19, 435 (2018).

71. Mu, P. et al. *SOX2* promotes lineage plasticity and antiandrogen resistance in *TP53*- and *RB1*-deficient prostate cancer. Science 355, 84–88 (2017).

72. Karachaliou, N., Rosell, R. & Viteri, S. The role of SOX2 in small cell lung cancer, lung adenocarcinoma and squamous cell carcinoma of the lung. Transl Lung Cancer Res 2, 172–9 (2013).

73. Rudin, C. M. et al. Comprehensive genomic analysis identifies SOX2 as a frequently amplified gene in small-cell lung cancer. Nat Genet 44, 1111 (2012).

74. Simpson, K. L. et al. A biobank of small cell lung cancer CDX models elucidates inter- and intratumoral phenotypic heterogeneity. Nat Cancer 1–15 (2020) doi:10.1038/s43018-020-0046-2.

75. Wang, T. et al. Transcription factor E2F1 promotes EMT by regulating ZEB2 in small cell lung cancer. Bmc Cancer 17, 719 (2017).

76. Gardner, E. E. et al. Chemosensitive Relapse in Small Cell Lung Cancer Proceeds through an EZH2-SLFN11 Axis. vol. 31 (2017).

77. Tripathi, S. C. et al. MCAM Mediates Chemoresistance in Small-Cell Lung Cancer via the PI3K/AKT/SOX2 Signaling Pathway. Cancer Res 77, 4414–4425 (2017).

78. Tendler, A., Mayo, A. & Alon, U. Evolutionary tradeoffs, Pareto optimality and the morphology of ammonite shells. Bmc Syst Biol 9, 12 (2015).

79. Szekely, P., Korem, Y., Moran, U., Mayo, A. & Alon, U. The Mass-Longevity Triangle: Pareto Optimality and the Geometry of Life-History Trait Space. Plos Comput Biol 11, e1004524 (2015).

80. Manno, G. L. et al. RNA velocity of single cells. Nature 560, (2018).

81. Bergen, V., Lange, M., Peidli, S., Wolf, F. A. & Theis, F. J. Generalizing RNA velocity to transient cell states through dynamical modeling. Nat Biotechnol 1–7 (2020) doi:10.1038/s41587-020-0591-3.

82. Hausser, J. & Alon, U. Tumour heterogeneity and the evolutionary trade-offs of cancer. Nat Rev Cancer 20, 247–257 (2020).

83. Huch, M. & Rawlins, E. L. Cancer: Tumours build their niche. Nature 545, 292 (2017).

84. Tammela, T. et al. A Wnt-producing niche drives proliferative potential and progression in lung adenocarcinoma. Nature 545, 355 (2017).

85. Williamson, S. C. et al. Vasculogenic mimicry in small cell lung cancer. Nat Commun 7, 13322 (2016).

86. Seftor, R. E. B. et al. Tumor Cell Vasculogenic Mimicry From Controversy to Therapeutic Promise. Am J Pathology 181, 1115–1125 (2012).

87. Chan, J. M. et al. Single cell profiling reveals novel tumor and myeloid subpopulations in small cell lung cancer. doi:10.1101/2020.12.01.406363.

88. Chen, H. J. et al. Generation of pulmonary neuroendocrine cells and SCLC-like tumors from human embryonic stem cellshESC-derived pulmonary neuroendocrine cells and tumors. J Exp Med 216, jem.20181155 (2019).

89. Yao, E. et al. Notch Signaling Controls Transdifferentiation of Pulmonary Neuroendocrine Cells in Response to Lung Injury. Stem Cells 36, 377–391 (2018).

90. Reynolds, S. D. et al. Conditional Clara cell ablation reveals a self-renewing progenitor function of pulmonary neuroendocrine cells. Am J Physiol-lung C 278, L1256–L1263 (2000).

91. Ouadah, Y. et al. Rare Pulmonary Neuroendocrine Cells Are Stem Cells Regulated by Rb, p53, and Notch. Cell 179, 403–416.e23 (2019).

92. Gu, X. et al. Chemosensory Functions for Pulmonary Neuroendocrine Cells. Am J Resp Cell Mol 50, 637–646 (2014).

93. Garg, A., Sui, P., Verheyden, J. M., Young, L. R. & Sun, X. Consider the lung as a sensory organ: A tip from pulmonary neuroendocrine cells. Curr Top Dev Biol (2019) doi:10.1016/bs.ctdb.2018.12.002.

94. Lommel, V. A. Pulmonary neuroendocrine cells (PNEC) and neuroepithelial bodies (NEB): chemoreceptors and regulators of lung development. Paediatric Respiratory Reviews 2, 171–176 (2001).

95. Domnik, N. J. & Cutz, E. Pulmonary neuroepithelial bodies as airway sensors: putative role in the generation of dyspnea. Curr Opin Pharmacol 11, 211–217 (2011).

96. Dean, C. H. & Snelgrove, R. J. New Rules for Club Development: New Insights into Human Small Airway Epithelial Club Cell Ontogeny and Function. Am J Resp Crit Care 198, 1355–1356 (2018).

97. Zuo, W.-L. et al. Ontogeny and Biology of Human Small Airway Epithelial Club Cells. Am J Resp Crit Care 198, 1375–1388 (2018).

98. Montoro, D. T. et al. A revised airway epithelial hierarchy includes CFTR-expressing ionocytes. Nature 560, 319–324 (2018).

99. Reynolds, S. D. & Malkinson, A. M. Clara cell: Progenitor for the bronchiolar epithelium. Int J Biochem Cell Biology 42, 1–4 (2010).

100. Widdicombe, J. G. & Pack, R. J. The Clara cell. Eur J Respir Dis 63, 202–20 (1982).

101. Aiello-Couzo, N. M. & Kang, Y. A bridge between melanoma cell states. Nat Cell Biol 22, 913–914 (2020).

102. Su, Y. et al. Multi-omic single-cell snapshots reveal multiple independent trajectories to drug tolerance in a melanoma cell line. Nat Commun 11, 2345 (2020).

103. Li, F. Z., Dhillon, A. S., Anderson, R. L., McArthur, G. & Ferrao, P. T. Phenotype Switching in Melanoma: Implications for Progression and Therapy. Frontiers Oncol 5, 31 (2015).

104. Kemper, K., Goeje, P. L. de, Peeper, D. S. & Amerongen, R. van. Phenotype Switching: Tumor Cell Plasticity as a Resistance Mechanism and Target for Therapy. Cancer Res 74, 5937–5941 (2014).

105. Ghandi, M. et al. Next-generation characterization of the Cancer Cell Line Encyclopedia. Nature 569, 503–508 (2019).

106. Barretina, J. et al. The Cancer Cell Line Encyclopedia enables predictive modelling of anticancer drug sensitivity. Nature 483, 603–607 (2012).

107. Cerami, E. et al. The cBio Cancer Genomics Portal: An Open Platform for Exploring Multidimensional Cancer Genomics Data. Cancer Discov 2, 401–404 (2012).

108. Gao, J. et al. Integrative Analysis of Complex Cancer Genomics and Clinical Profiles Using the cBioPortal. Sci Signal 6, pl1–pl1 (2013).

109. Leek, J. T. & Storey, J. D. Capturing Heterogeneity in Gene Expression Studies by Surrogate Variable Analysis. Plos Genet 3, e161 (2007).

110. Johnson, W. E., Li, C. & Rabinovic, A. Adjusting batch effects in microarray expression data using empirical Bayes methods. Biostatistics 8, 118–127 (2007).

111. Brock, G., Pihur, V., Datta, S. & Datta, S. clValid: An R Package for Cluster Validation Brock Journal of Statistical Software. (2008).

112. Klein, A. M. et al. Droplet Barcoding for Single-Cell Transcriptomics Applied to Embryonic Stem Cells. Cell 161, 1187–1201 (2015).

113. Petukhov, V. et al. dropEst: pipeline for accurate estimation of molecular counts in droplet-based single-cell RNA-seq experiments. Genome Biol 19, 78 (2018).

114. Dobin, A. et al. STAR: ultrafast universal RNA-seq aligner. Bioinformatics 29, 15–21 (2013).

115. Wolf, F. A., Angerer, P. & Theis, F. J. SCANPY: large-scale single-cell gene expression data analysis. Genome Biol 19, 15 (2018).

116. Hie, B., Bryson, B. & Berger, B. Efficient integration of heterogeneous single-cell transcriptomes using Scanorama. Nat Biotechnol 37, 685–691 (2019).

117. Wolock, S. L., Lopez, R. & Klein, A. M. Scrublet: Computational Identification of Cell Doublets in Single-Cell Transcriptomic Data. Cell Syst 8, 281–291.e9 (2019).

118. Roweis, S. T. & Saul, L. K. Nonlinear Dimensionality Reduction by Locally Linear Embedding. Science 290, 2323–2326 (2000).

119. Huang, S. Cell Lineage Determination in State Space: A Systems View Brings Flexibility to Dogmatic Canonical Rules. Plos Biol 8, e1000380 (2010).

120. Wahl, G. M. & Spike, B. T. Cell state plasticity, stem cells, EMT, and the generation of intra-tumoral heterogeneity. Npj Breast Cancer 3, 14 (2017).

121. Yadav, T., Quivy, J.-P. & Almouzni, G. Chromatin plasticity: A versatile landscape that underlies cell fate and identity. Science 361, 1332–1336 (2018).

122. Ahmed, E. et al. Lung development, regeneration and plasticity: From disease physiopathology to drug design using induced pluripotent stem cells. Pharmacol Therapeut 183, 58–77 (2018).

123. Scheel, C. & Weinberg, R. A. Phenotypic plasticity and epithelial-mesenchymal transitions in cancer and normal stem cells? Int J Cancer 129, 2310–2314 (2011).

124. Tata, P. R. & Rajagopal, J. Plasticity in the lung: making and breaking cell identity. Development 144, 755–766 (2017).

125. Wagers, A. J. & Weissman, I. L. Plasticity of Adult Stem Cells. Cell 116, 639–648 (2004).

126. Burclaff, J. & Mills, J. C. Plasticity of differentiated cells in wound repair and tumorigenesis, part I: stomach and pancreas. Dis Model Mech 11, dmm033373 (2018).

127. Hurley, K. et al. Reconstructed Single-Cell Fate Trajectories Define Lineage Plasticity Windows during Differentiation of Human PSC-Derived Distal Lung Progenitors. Cell Stem Cell 26, 593–608.e8 (2020).

128. Risom, T. et al. Differentiation-state plasticity is a targetable resistance mechanism in basal-like breast cancer. Nat Commun 9, 3815 (2018).

129. Marjanovic, N. D. et al. Emergence of a High-Plasticity Cell State during Lung Cancer Evolution. Cancer Cell (2020) doi:10.1016/j.ccell.2020.06.012.

130. Flavahan, W. A., Gaskell, E. & Bernstein, B. E. Epigenetic plasticity and the hallmarks of cancer. Science 357, eaal2380 (2017).

131. Neftel, C. et al. An Integrative Model of Cellular States, Plasticity, and Genetics for Glioblastoma. Cell 178, 835–849.e21 (2019).

132. Marjanovic, N. D., Weinberg, R. A. & Chaffer, C. L. Cell Plasticity and Heterogeneity in Cancer. Clin Chem 59, 168–179 (2013).

133. Quintanal-Villalonga, Á. et al. Lineage plasticity in cancer: a shared pathway of therapeutic resistance. Nat Rev Clin Oncol 1–12 (2020) doi:10.1038/s41571-020-0340-z.

134. Wooten, D. J. & Quaranta, V. Mathematical models of cell phenotype regulation and reprogramming: Make cancer cells sensitive again! Biochim Biophys Acta 1867, (2017).

135. Weinreb, C., Wolock, S., Tusi, B. K., Socolovsky, M. & Klein, A. M. Fundamental limits on dynamic inference from single-cell snapshots. Proc National Acad Sci 115, 201714723 (2018).

136. Weinreb, C., Rodriguez-Fraticelli, A., Camargo, F. D. & Klein, A. M. Lineage tracing on transcriptional landscapes links state to fate during differentiation. Sci New York N Y eaaw3381 (2020) doi:10.1126/science.aaw3381.

137. Tritschler, S. et al. Concepts and limitations for learning developmental trajectories from single cell genomics. Development 146, dev170506 (2019).

138. Hochgerner, H., Zeisel, A., Lönnerberg, P. & Linnarsson, S. Conserved properties of dentate gyrus neurogenesis across postnatal development revealed by single-cell RNA sequencing. Nat Neurosci 21, 290–299 (2018).

139. Bastidas-Ponce, A. et al. Comprehensive single cell mRNA profiling reveals a detailed roadmap for pancreatic endocrinogenesis. Development 146, dev173849 (2019).

140. Pirttimaki, T. M. & Parri, H. R. Astrocyte Plasticity. Neurosci 19, 604–615 (2013).

141. Zhou, Y. et al. Dual roles of astrocytes in plasticity and reconstruction after traumatic brain injury. Cell Commun Signal 18, 62 (2020).

142. Migliorini, A., Bader, E. & Lickert, H. Islet cell plasticity and regeneration. Mol Metab 3, 268–274 (2014).

143. Habener, J. F. & Stanojevic, V. α-cell role in β-cell generation and regeneration. Islets 4, 188–198 (2012).

144. Furuyama, K. et al. Diabetes relief in mice by glucose-sensing insulin-secreting human α-cells. Nature 567, 43–48 (2019).

